# Persistent trade-offs balance competition and colonization across centuries

**DOI:** 10.1101/2025.11.16.688059

**Authors:** Talia Backman, Jiajun Cui, Emma Caullireau, Ella Bleak, Ilja Bezrukov, Patricia Girardi, Aubrey Hawks, Jesse R. Lasky, Sergio M. Latorre, Joel M. Erberich, Lua Lopez, Manuela Neumann, Allison M. Perkins, Efthymia Symeonidi, Parastoo Azadi, Martin P. Horvath, Artur Muszyński, Patricia L. M. Lang, Talia L. Karasov, Hernán A. Burbano

**Author notes:** Equal contribution (Talia Backman, Jiajun Cui, Emma Caullireau, Talia L. Karasov, Hernán A. Burbano). Corresponding author: TLK, HAB. **Classification:** Biological Sciences/Evolution. ORCID: TB, JC, EC, EB, IB, PG, AH, JL, SL, JE, LL, MN, AP, ES, PA, MPH, AM, PL.

## Abstract

Microbial competition drives rapid adaptation, often forcing organisms to specialize in new ecological niches. Adaptations that improve competitive ability can reduce performance in other environments creating trade-offs. Whether such trade-offs persist in nature—or are eroded as lineages adapt through compensatory changes—remains largely unknown. Here we show that a trade-off between competitive ability and host colonization has been stably maintained in natural *Pseudomonas* populations for centuries. Wild plant-pathogenic *Pseudomonas* compete using tailocins—phage-derived molecular weapons that bind to specific cell-surface receptors. Genomic surveys and functional assays reveal that the most broadly lethal tailocins remain rare—while the tailocin’s production increases competitive killing, it also compromises plant colonization. We determine that the polymorphisms behind this trade-off are not transient — historical genomes spanning two centuries show that the trade-off has been maintained for at least 10⁵–10⁶ generations. Our results demonstrate that, in natural populations, a trade-off between competition and pathogenicity is fundamental and not easily overcome.

**Significance:** When a microbe colonizes a host, it must both establish infection and outcompete other organisms. Short-term experiments show that gains in competitive ability can reduce colonization, creating trade-offs, but whether microbes resolve these conflicts over long evolutionary timescales is unknown. We show that a trade-off between competitive killing and host colonization has been stably maintained for centuries in natural *Pseudomonas* populations infecting *Arabidopsis thaliana*. Tailocins—phage-derived weapons—provide strong competitive advantages, yet their production reduces colonization success, explaining why the most broadly lethal variants remain rare. Genomic surveys and historical genomes spanning two centuries reveal that the polymorphisms underlying this trade-off have persisted across 10⁵-10⁶ generations. Understanding such long-lived constraints can inform antimicrobial strategies that exploit evolutionary trade-offs.

## Introduction

Competition between microbes is an ecological and evolutionary force that shapes community structure, resource use, and diversification ^1^. Experimental evolution has shown that competition can drive rapid adaptation to new ecological niches ^2^, yet these adaptive gains often come at a cost in other contexts. The resulting trade-offs—between competitive ability, growth, and colonization—are hypothesized to be pervasive in microbial communities ^3^. However, most evidence for such trade-offs comes from experimental evolution in laboratory systems ^4^, leaving open the question of whether they constrain populations in wild environments and over longer, evolutionary timescales. Do microbes in the wild overcome these trade-offs, optimizing performance across environments, or do they remain trapped by enduring conflicts between competition and colonization?

We address this question in *Pseudomonas viridiflava*, a common plant-pathogenic bacterium that infects *Arabidopsis thaliana* across Europe ^5,6^. Within these populations, a pathogenic phylotype named ATUE5 forms a genetically diverse metapopulation in which no single strain dominates ^6^. Intraspecific competition within ATUE5 is an important selective pressure ^7^ mediated by molecular weapons termed tailocins. Tailocins are diffusible protein complexes that resemble phage tails and kill susceptible strains by puncturing their cell membranes ^8^. Bacteria are resistant to their own tailocins upon encounter in the environment, allowing producer clones to suppress non-self competitors while sparing their own lineage ^9^. Because tailocins are released through cell lysis, the producing cell dies. Consequently, only a fraction of a clonal population produces tailocins, thereby protecting clonemates and suppressing competing strains ^9^.

The tailocin gene cluster in ATUE5 consists of ∼24 collinearly arranged genes, integrated at the same genomic location in all strains, which together assemble the functional tailocin. Each ATUE5 strain encodes one of a few tailocin variants, distinguished by the tail fiber (HTF), a receptor-binding protein that dictates killing specificity ^7,8^. The HTF binds to the outer membrane of the target cell, specifically recognizing the O-antigen, a highly variable component of the bacterial lipopolysaccharide (LPS) layer. In ATUE5, *HTF* genes vary in length (1245–1830 bp) and killing spectrum, with distinct length classes corresponding to unique O-antigen biosynthesis gene profiles, suggesting tight linkage between *HTF* variants and their O-antigen receptors within each tailocin-producing bacterium.

Paradoxically, bacteria that produce the most lethal tailocin variant—capable of killing ∼80% of other strains and resistant to ∼70% of others—remain rare in natural populations. Why has the most competitive variant not spread to fixation? We hypothesized that its persistence at low frequency may reflect a trade-off between competitive dominance and host colonization. The O-antigen provides a mechanistic link between these processes ^10^. O-antigen diversity influences surface recognition, virulence, and susceptibility to tailocins ^11,12^ potentially generating an evolutionary tension between success within the host and resistance to microbial attack.

To test for this competition-colonization trade-off, we combined saturation mutagenesis with functional assays and *in planta* infections. The genetic mutations most important for surviving in the presence of the tailocin also compromised growth in the plant. This experiment showed a short-term trade-off acting on contemporary bacterial strains. We then examined whether this trade-off showed evolutionary persistence by analyzing 49 historical bacterial genomes recovered from herbarium specimens spanning more than two centuries. Together, the modern and historical population datasets provide a uniquely powerful framework to identify the association between tailocin and O-antigen polymorphisms underlying the competition-colonization trade-off, and then to assess whether the trade-off is transient or an enduring feature of bacterial evolution.

Here, we show that O-antigen–dependent trade-offs between pathogenicity and interbacterial competition have persisted for over two centuries in natural *P. viridiflava* populations. O-antigen production enhances virulence but increases susceptibility to tailocin attack. By integrating historical genomics with functional microbiology, we reveal the genetic basis, ecological significance, and remarkable temporal stability of this trade-off. These findings demonstrate that structural constraints on microbial competition and colonization can maintain genetic diversity across evolutionary timescales, illuminating why the most competitive strains do not necessarily dominate in nature.

## Results

### The growth-defense trade-off: the O-antigen biosynthesis cluster (OBC) reduces resistance to interbacterial killing by tailocins but enhances colonization *in planta*

Our previous work established four major *HTF* length variants in *P. viridiflava* populations (1245, 1383, 1803, and 1830 bp) ^7^. These *HTF* length variants correlate, within the same bacterial genome, with the presence or absence of specific genes in the O-antigen biosynthesis cluster (OBC) that are critical for tailocin binding. The tight linkage between *HTF* length variants and OBC haplotypes within a bacterial genome suggests that the tail fiber and its O-antigen receptor are evolving in concert in natural populations of *P. viridiflava*. To determine the killing spectrum for each of these *HTF* length variants, we conducted killing assays that included at least one representative from each variant. The individual tailocin preparations were applied to a panel of 50 *P. viridiflava* tester strains that captured the diversity of the focal OBC presence and *HTF* length variants in the metapopulation (**Fig. 1A**). As shown previously, the 1245 bp and 1803 bp *HTF* variants are correlated in the genome of modern strains with OBC⁺, whereas the 1383 bp and 1830 bp variants occur only in OBC⁻ strains ^7^. The resulting killing spectra revealed a strong pattern: tailocins from OBC⁻ strains killed on average 79% of tester strains, while those from OBC⁺ strains killed 51% (**Fig. 1B**). These results provide preliminary evidence that OBC^-^ strains are more successful interbacterial competitors. Despite this apparent advantage, the OBC^-^ strains represent only ∼16% of the population. We thus hypothesized that loss of O-antigen in the OBC⁻ genotypes leads to reduced fitness in other natural conditions such as during plant infection ^13^.

**Figure 1:**
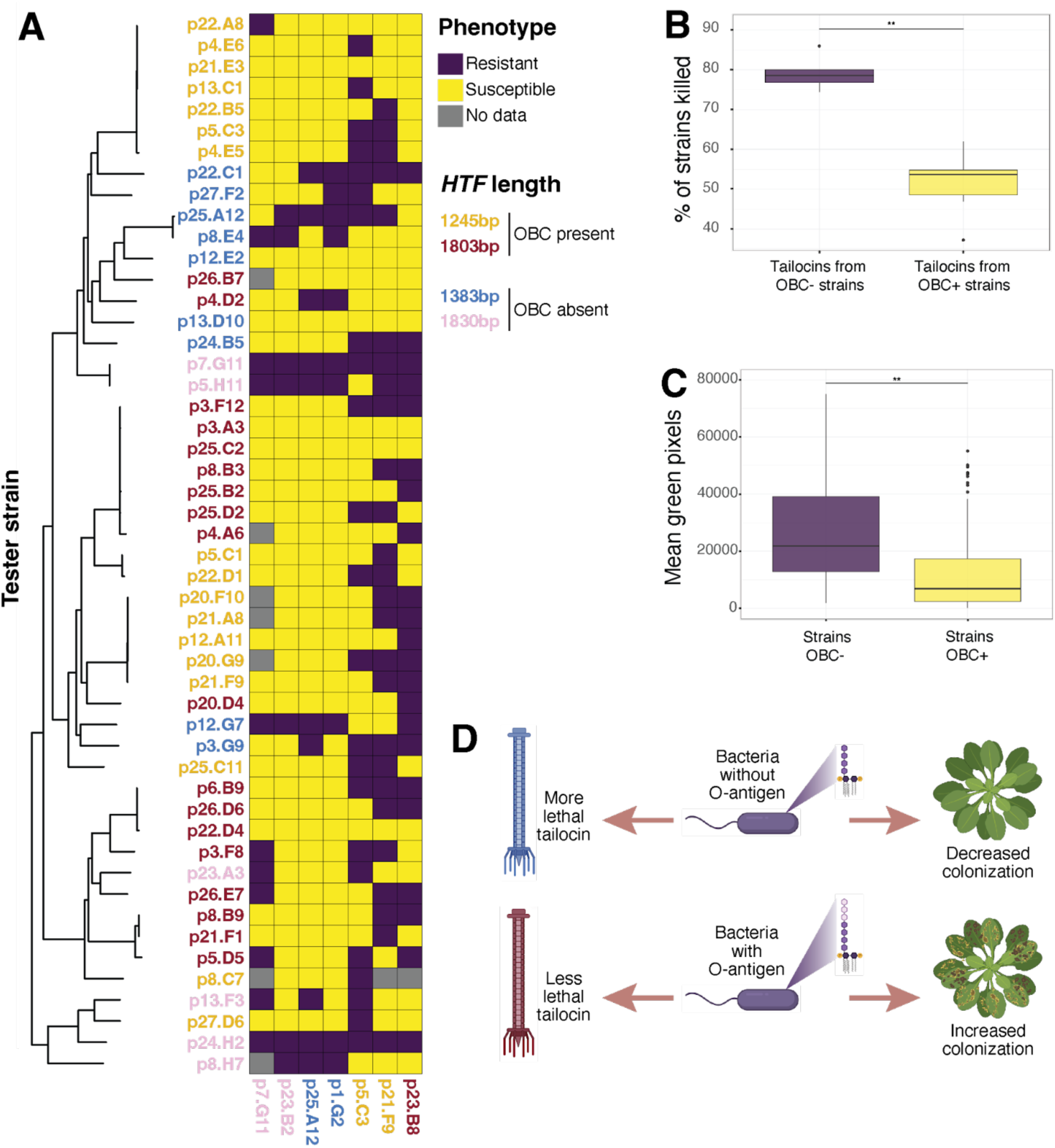
Decreased tailocin activity and increased virulence are associated with the presence of the focal O-antigen Biosynthesis Cluster (OBC) in wild populations of *P. viridiflava*. **A.** Tailocin killing matrix showing sensitivity (yellow), resistance (purple), or missing data (gray) across a panel of 50 *P. viridiflava* strains. Each row is a tester strain, and each column represents a tailocin from a naturally co-occurring producer strain, annotated below the heatmap and colored by its tail fiber (HTF) length variant. There are representative tailocins from all four HTF length variants previously characterized. **B.** Tailocins from OBC- strains exhibit significantly broader killing spectra than those from OBC+ strains, killing a higher proportion of tester strains (two-sided Wilcoxon rank-sum test, p < 0.01; data derived from A). Boxes display the median, interquartile range, and individual outliers. **C.** Infections with OBC- strains resulted in significantly higher mean green pixel counts compared to OBC+ strains, indicating reduced disease severity (n = 77 strains; two-sided Wilcoxon rank-sum test, W = 574, p = 0.009963). Boxes display the median, interquartile range, and outliers. **D.** Schematic describing the hypothesis for why we see variation in the success of competitors and colonizers in wild plant populations. Created in https://BioRender.com.

To test the colonization hypothesis, we infected *A. thaliana* seedlings with a subset of wild *P. viridiflava* strains that varied in *HTF* length and, therefore, OBC presence. Plant health was quantified using green pixel counts from high-quality plant images, where lower green pixel counts corresponded to a more diseased plant. We found that OBC^+^ strains (associated with the 1245 bp and 1803 bp *HTF* haplotypes) caused more severe disease phenotypes than OBC⁻ strains (associated with the 1383 bp and 1830 bp *HTF* haplotypes) (**Fig. 1C**). These results provide preliminary data that suggest that OBC^+^ strains may have enhanced aggressiveness in this experimental context. Together, these results suggest a trade-off between interbacterial competition and host colonization: OBC⁻ strains gain a competitive edge through potent tailocins but suffer reduced virulence *in planta*, whereas OBC⁺ strains colonize hosts more effectively but are more vulnerable to tailocin-mediated killing (**Fig. 1D**).

### OBC genes mediate a trade-off between tailocin resistance and plant colonization

We next examined whether disrupting O-antigen biosynthesis within a single genetic background reproduces the population-level trade-off between tailocin resistance and host colonization. We chose to focus on the model *P. viridiflava* strain p25.C2 ^7,14^ which is OBC^+^ and susceptible to the tailocin produced by strain p25.A12 ^7^. To systematically identify genes involved in both tailocin susceptibility and fitness *in planta*, we employed high-throughput Random-Barcoded Transposon (Tn) Sequencing (RB-TnSeq) ^15^. This approach uses uniquely barcoded transposon mutants to measure each gene’s contribution to fitness across different growth conditions. We previously performed RB-TnSeq fitness in liquid culture of p25.C2 treated with strain p25.A12’s tailocin to identify mutations (Tn-insertions) that render p25.C2 resistant to p25.A12’s tailocin ^7^. Here, we performed RB-TnSeq with the same library in plant infections (*A. thaliana* ecotypes Col-0 and Eyach 1.5-2) to identify mutations that influence fitness in the plant hosts. We tracked the abundance of mutants after growth either in a tailocin environment or in the plants, which provided relative fitness in the presence of tailocins versus within the plant host (**Fig. 2A**). As reported previously, mutants in the OBC significantly increased in abundance following tailocin application, indicating that disruption of these genes confers resistance (**Fig. 2B**, log fold change tailocin > 0) ^7^.

**Figure 2:**
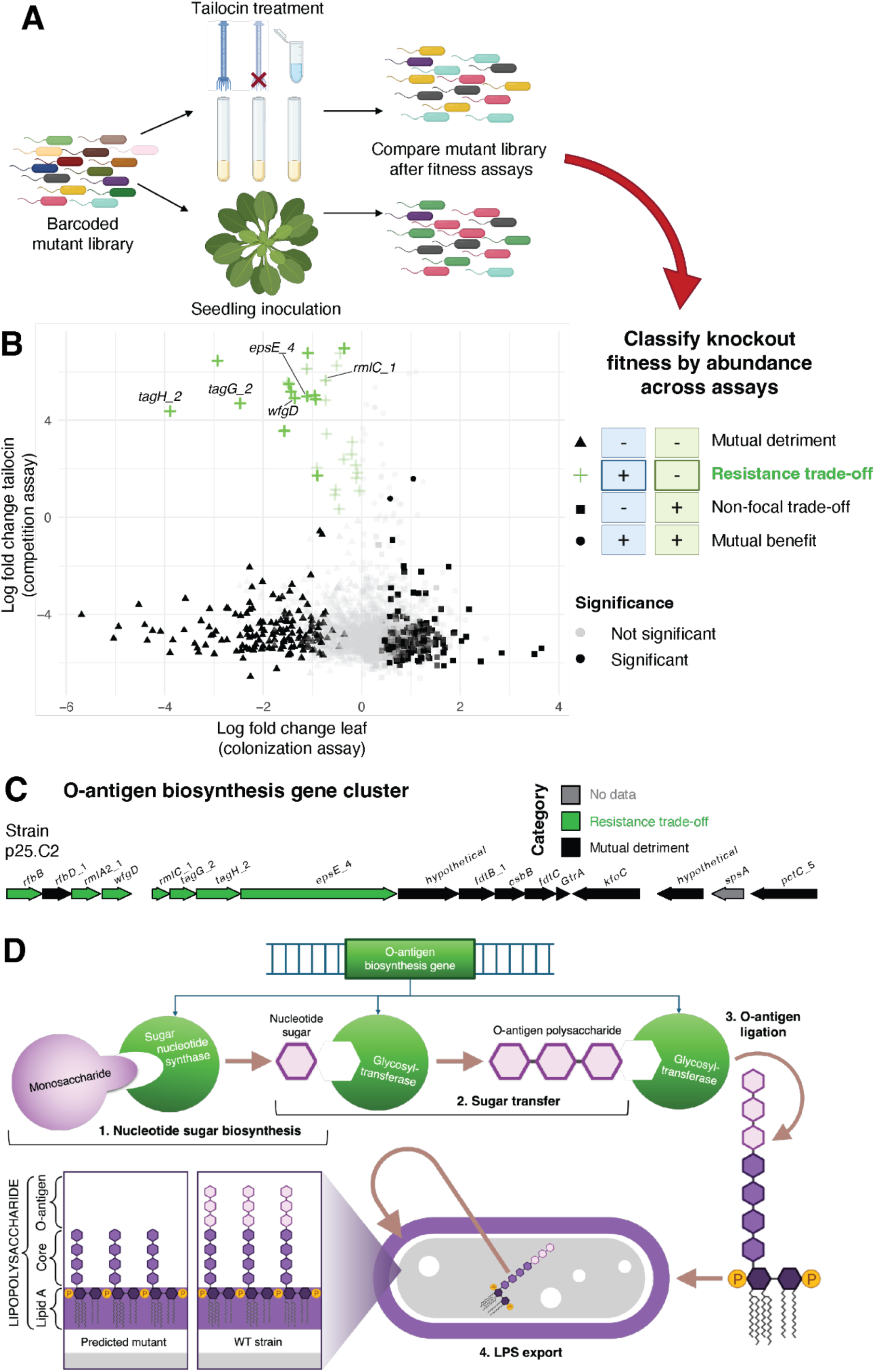
Tailocin resistance mutations in the OBC impair bacterial colonization in *Arabidopsis thaliana*. **A.** Schematic of the RB-TnSeq experimental design. A barcoded transposon mutant library of *P. viridiflava* strain p25.C2 was subjected to two separate treatments: (i) tailocin exposure in liquid culture (data from Backman et al. 2024), and (ii) leaf infiltration into *A. thaliana* ecotypes Eyach 1.5-2 and Col-0. Mutant fitness was inferred from barcode abundance post-selection, and each gene was categorized based on relative fitness across both conditions together: –/– = mutually detrimental, +/+ = mutually beneficial, +/– = resistance trade-off, –/+ = mon-focal trade-off. **B.** Scatterplot of the log_2_ fold change in barcode abundance during leaf colonization in Eyach 1.5-2 (x-axis) versus tailocin treatment (y-axis). Each point represents one gene (mean across barcodes). Significance is indicated by opacity: opaque points mark genes significant in both ecotypes (p < 0.05), while lighter points indicate nonsignificance in at least one condition. Shapes denote fitness outcome categories (mutual benefit, mutual detriment, non-focal trade-off, or resistance trade-off in both ecotypes). Labeled genes correspond to focal O-antigen biosynthesis cluster resistance trade-off genes. **C.** Gene diagram of the OBC on the *P. viridiflava* strain p25.C2 chromosome. Genes are colored based on statistical significance in each assay: resistance trade-off (green), mutual detriment (black), or no data (grey). **D.** Simplified schematic of O-antigen biosynthesis. (1) Nucleotide sugars are synthetized from phosphorylated monosaccharide precursors. (2) Sugars are sequentially transferred onto a lipid carrier, forming the O-antigen repeat unit and eventually the O-antigen polymer. (3) The O-antigen is ligated to the lipid A-core oligosaccharide to produce lipopolysaccharide (LPS), which is transported across the outer membrane (4). Resistance trade-off genes encode sugar nucleotide synthases and glycosyltransferases involved in these steps, indicating that disrupting any part of this pathway can confer tailocin resistance at the cost of reduced colonization. Created in https://BioRender.com.

To determine how these tailocin resistance mutations influence fitness in the host, we plotted each mutant fold-change in abundance during plant colonization (in Eyach 1.5-2; x-axis) against its abundance change under tailocin application (y-axis; **Fig. 2B**). Among the 70 mutations (putative gene knockouts) associated with tailocin resistance ^7^, 21 significantly reduced bacterial fitness in at least one ecotype, with 13 (61%) showing reduced fitness across both ecotypes (**Fig. 2B**, **Table S1**). We focus on this set of 21 genes, which constitute the “trade-off” genes, for which transposon insertions increase fitness upon tailocin challenge, but decrease fitness during colonization.

We then examined the predicted function of these 21 trade-off candidate genes: seven are found within a single 18 kb operon predicted to synthesize the focal OBC (**Fig. 2C**). These genes encode sugar nucleotide synthases (*rfbB*, *rmlA2_1*, *wfgD*, *rmlC_1*) and glycosyltransferases (*tagG_2*, *tagH_2*, *epsE_4*) responsible for the sequential transfer of sugars involved in O-antigen initiation, elongation, polymerization, or ligation (**Fig. 2D**). Putative knockouts in genes involved in this pathway increased survival under tailocin treatment but reduced fitness during leaf colonization, supporting the hypothesized competition-versus-colonization trade-off, regardless of the phase in O-antigen biosynthesis. The same pattern emerged, although with smaller effect sizes, for additional LPS- and O-antigen-related genes elsewhere in the genome, suggesting that LPS and O-antigen modifications are common for tailocin-resistant mutations that result in reduced colonization (**Table S1**).

The overlap in the RB-TnSeq results in two *A. thaliana* ecotypes validates the predicted trade-off, suggesting that the OBC resistance mechanism is associated with impaired plant colonization, thereby prioritizing the OBC for further functional analysis.

### O-antigen mutants alter LPS structure and reduce susceptibility to tailocins and bacterial load *in planta*

To confirm the contribution of the O-antigen deletion on *P. viridiflava* growth in the plant niche, we generated site-directed deletion mutants. We generated clean deletion mutants via homologous recombination for six candidate OBC gene mutants. These included putative nucleotide sugar biosynthetic enzymes (*wfgD*, *rmlC_1*), and glycosyltransferases (*tagG_2*, *tagH_2*, *epsE_4*, *spsA*; **Fig. 2C**). *spsA* was included in the mutant analysis given its position within the same predicted operon and its prior functional annotation as an O-antigen biosynthetic gene as previously reported ^7^.

These gene disruptions altered LPS structure, as shown by the electrophoretic profiles of purified LPS (**Fig. 3A**). Five of six mutants (*ΔwfgD*, *ΔtagG_2*, *ΔtagH_2*, *ΔepsE_4*, and *ΔspsA)* lack high-molecular-weight bands relative to WT, consistent with a modification or absence of O-antigen polymer (silver-stained DOC-PAGE, **Fig. 3A**). In contrast, *ΔrmlC_1,* a deletion of a gene with a sequence homolog in the same genome, retained a higher-molecular-weight band, as observed in WT. Chemical compositional analysis further supported these LPS modifications, showing a shift in sugar residues with a marked loss of rhamnose in all mutants except *ΔrmlC_1* (**Table S2**).

**Figure 3:**
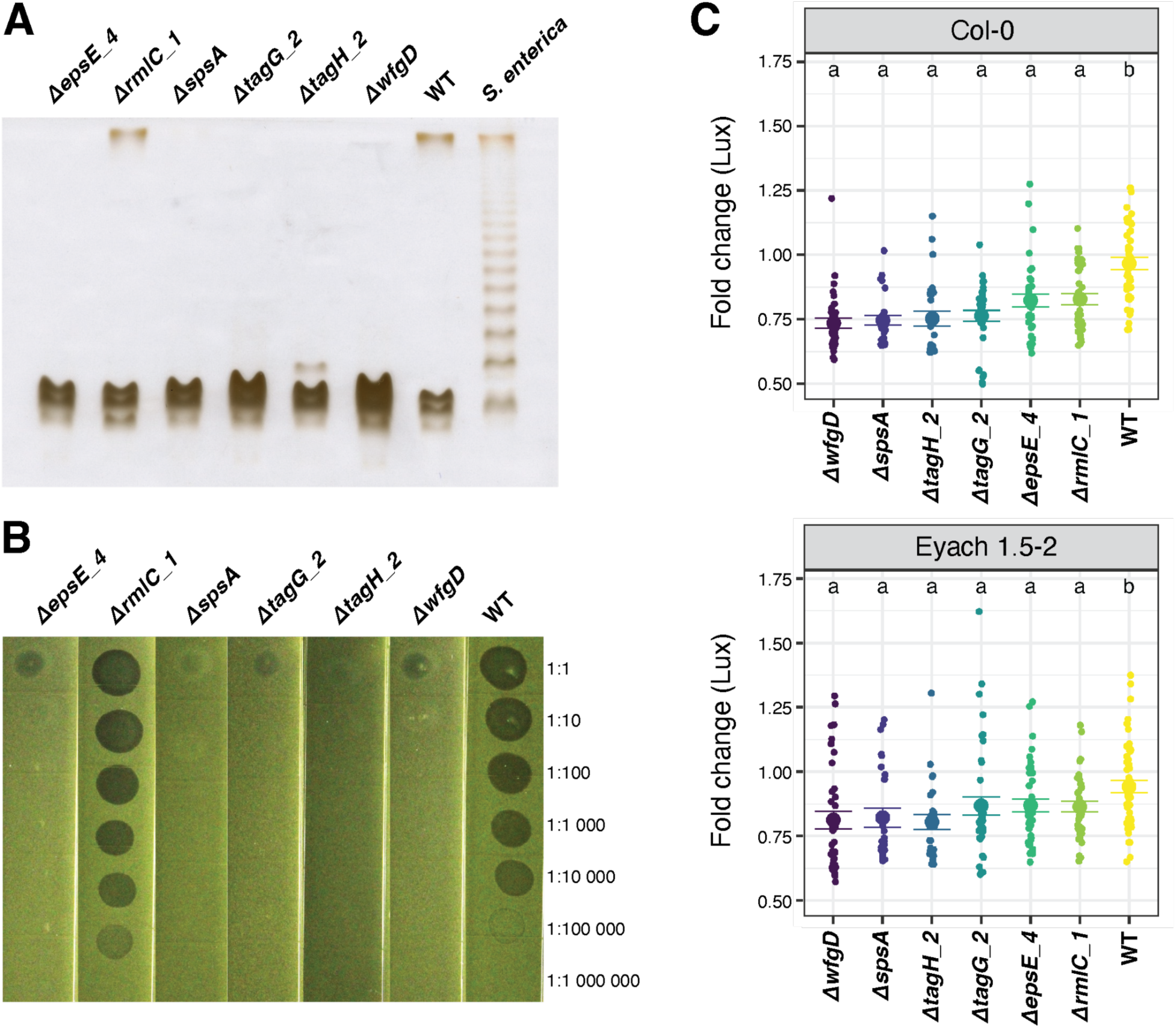
O-antigen mutants lack O-antigen chains, proliferate less *in planta*, and improve plant health outcomes. **A.** The LPS of the O-antigen knockouts were extracted and visualized with a silver-stained DOC-PAGE gel. Some mutants lack the high molecular weight O-antigen band, while *ΔrmlC_1* still has the O-antigen band. **B.** Tailocin killing assay using partially purified tailocin from p25.A12 applied to soft agar with each O-antigen mutant or wild-type (WT). Killing is shown by zones of inhibition. Six mutants exhibit resistance to tailocin-mediated killing. *ΔrmlC_1* appears to be even more sensitive than WT. Representative results from three GRGbiological replicates are shown. **C.** *Pseudomonas* strains tagged with luciferase were flood-inoculated onto *Arabidopsis thaliana* seedlings (Col-0 or Eyach 1.5-2 background). Bacterial load was quantified by luminescence 7 days post-infection (dpi). Points represent replicate measurements of fold change in Lux signal from 0 dpi to 7 dpi (log-transformed). Mutants generally showed reduced bacterial growth compared to wild-type (WT). Horizontal bars indicate mean ± standard error; letters indicate statistical groupings by a linear mixed model with random intercept for batch.

Next, tailocin susceptibility of the mutants was assessed using a soft agar overlay assay with partially purified tailocin from p25.A12 (**Fig. 3B**). The five mutants (*ΔwfgD*, *ΔtagG_2*, *ΔtagH_2*, *ΔepsE_4*, and *ΔspsA)* exhibited resistance to tailocin-mediated killing, while *ΔrmlC_1* showed killing zones similar to WT. We then tested whether O-antigen loss affects bacterial growth and disease *in planta*, by flood-inoculating *A. thaliana* seedlings with WT and O-antigen mutants. We additionally assessed bacterial population size dynamics as well as plant health 7 days post-inoculation (dpi). Across both *Arabidopsis* ecotypes, O-antigen mutants generally exhibited reduced bacterial load compared to WT at 7 dpi, consistent with a fitness cost of resistance to tailocin (**Fig. 3C**). In Col-0, all six mutants showed significantly lower bacterial load than WT, with highly significant differences observed for all strains after FDR correction (all p < 0.0001). In Eyach 1.5-2, three mutants (*ΔspsA*, *ΔtagH_2*, and *ΔwfgD*) exhibited strongly reduced growth relative to WT (FDR-adjusted p < 0.005), while *ΔepsE_4*, *ΔrmlC_1,* and *ΔtagG_2* were moderately reduced (FDR-adjusted p ≈ 0.05). Notably, these results were independent of the inoculation method, as similar outcomes were observed following the syringe infiltration of one O-antigen mutant into plant leaves (**Fig. S1**), suggesting that the role of the O-antigen for colonization extends beyond the leaf surface (epiphytic phase) to the inside of the plant tissue (endophytic phase). Deletion of O-antigen synthesis genes conferred improved plant health outcomes relative to wild-type (WT): in both ecotypes, infections with WT strain frequently resulted in chlorosis or necrosis, whereas infections with O-antigen mutants more often led to asymptomatic, healthy plants (**Fig. S2**). Furthermore, plant health tended to decline overall with increasing bacterial load (**Fig. S3**). Together, these findings validate the function of the insertion mutant candidate genes for an O-antigen-mediated trade-off between tailocin resistance and host colonization with robust and consistent effects across multiple genetic backgrounds.

### O-antigen loss affects growth properties *in vitro* but does not affect tolerance to stresses nor host immunity activation

We investigated the biological basis of the reduced bacterial fitness observed *in planta* by testing several hypotheses. Specifically, we considered whether this trade-off could result from (i) a general physiological defect affecting bacterial growth properties, (ii) a higher susceptibility to antimicrobials or reduced stress tolerance due to membrane permeability, and/or (iii) a stronger activation of host immunity due to a faster pathogen recognition.

We first compared the *in-vitro* growth of the WT and O-antigen deletion mutant strains and observed that all mutants, except *ΔrmlC_1*, displayed a growth defect in liquid cultures grown in rich medium (**Fig. S4**). This defect was statistically significant based on several growth parameters, including the growth rates (*k* or *r*), generation time, and the area under the curve (AUC; **Table S3**). We next assessed surface-associated growth of the WT and the O-antigen mutant strains using a 96-well static assay. The WT and *ΔrmlC_1* strains consistently showed no visible aggregate formation, whereas all other O-antigen mutants formed robust surface-associated aggregates in nearly every well after 48 hours of incubation (≥ 95% of replicates, Pearson’s χ² = 241.5, p < 2 × 10^-16^; **Fig. S5**). All O-antigen mutants except *ΔrmlC_1* produced significantly more aggregates than the WT (pairwise Fisher’s exact test, adjusted p < 10^-16^).

Secondly, to test whether O-antigen acts as an antimicrobial barrier during plant colonization, we compared the tolerance of tailocin-resistant O-antigen mutants and the WT to two stressors: the cationic peptide polymyxin B (PMB) and hydrogen peroxide (H_2_O_2_). No significant difference in growth inhibition was observed between the WT and the O-antigen *P. viridiflava* mutants in PMB disk diffusion assays, regardless of the compound concentration (**Fig. S6A**). Similar results were also observed with another polymyxin (colistin sulfate, **Fig. S6B**). These results indicate that the O-antigen does not contribute detectably to polymyxin tolerance in our strains, in contrast to previous reports describing high sensitivity to PMB of O-antigen mutants in *P. aeruginosa^16^*. Furthermore, growth curves across a gradient of H_2_O_2_ concentrations revealed that all strains shared the same inhibitory profile, with indistinguishable minimum inhibitory concentrations (MICs; MIC/WT ≈ 1.0 for all strains; **Table S4**) and parallel declines in AUC as H_2_O_2_ increased (**Fig. S7**). Mixed-effects modeling of five independent replicates confirmed that O-antigen mutants did not differ significantly from WT in their AUC responses or MIC values after accounting for replicate-level variation.

Finally, we tested whether the O-antigen influences host immunity activation. Focusing on six marker genes associated with early (*FRK1*, *WRKY29*, *NHL10*, *PAD3*) or late (*PDF1.2*, *PR1*) plant defense responses, we monitored their expression from 0 to 72 hours post-infection with either the WT strain or selected O-antigen mutants. We found no evidence of a stronger or faster plant defense activation that could account for the reduced fitness of the O-antigen mutants *in planta* (**Fig. S8**).

Together, these data indicate that the loss of O-antigen (i) generally affects bacterial growth *in vitro* and promotes surface-associated aggregate formation, but is not relevant for (ii) tolerance to reactive oxygen species or cationic peptides under laboratory conditions, and (iii) activation of plant immunity. This suggests that the ecological trade-offs of O-antigen loss observed *in planta* likely result from general physiological changes that lead to altered bacterial growth properties.

### Herbarium-derived genomes reveals the persistence of the trade-off across centuries

Together, our functional and genetic results reveal a clear trade-off: loss of O-antigen confers resistance to tailocins but reduces bacterial fitness during plant colonization. This raises a key evolutionary question: is this trade-off transient, or has it constrained *P. viridiflava* populations over longer, ecological and historical timescales? To investigate this, we used ancient DNA (aDNA) techniques ^17^ to sequence 49 metagenomic libraries from *A. thaliana* herbarium specimens collected across Europe between 1817 and 2015 (**Fig. 4A**). This ∼200 year period corresponds to an estimated 10^5^-10^6^ generations for *P. viridiflava ^18^*. On average, 5.0% (0.2–36.3%) of metagenomic reads mapped to the *P. viridiflava* ATUE5 reference genome (**Fig. S9A**). The herbarium-derived reads covered, on average, 82.5% (59.8–95.1%) of the ATUE5 genome, with an average sequencing depth of 14.7x (1.1-137.4x) (**Fig. S9B**). The *Pseudomonas*-mapped reads showed patterns of DNA damage and fragmentations typical of aDNA ^19^ (**Fig. S9C; Fig. S10; Fig. S11**) and the distribution of *k*-mer coverage revealed that most historical infections are dominated by a single strain (**Fig. S9D**).

**Figure 4.**
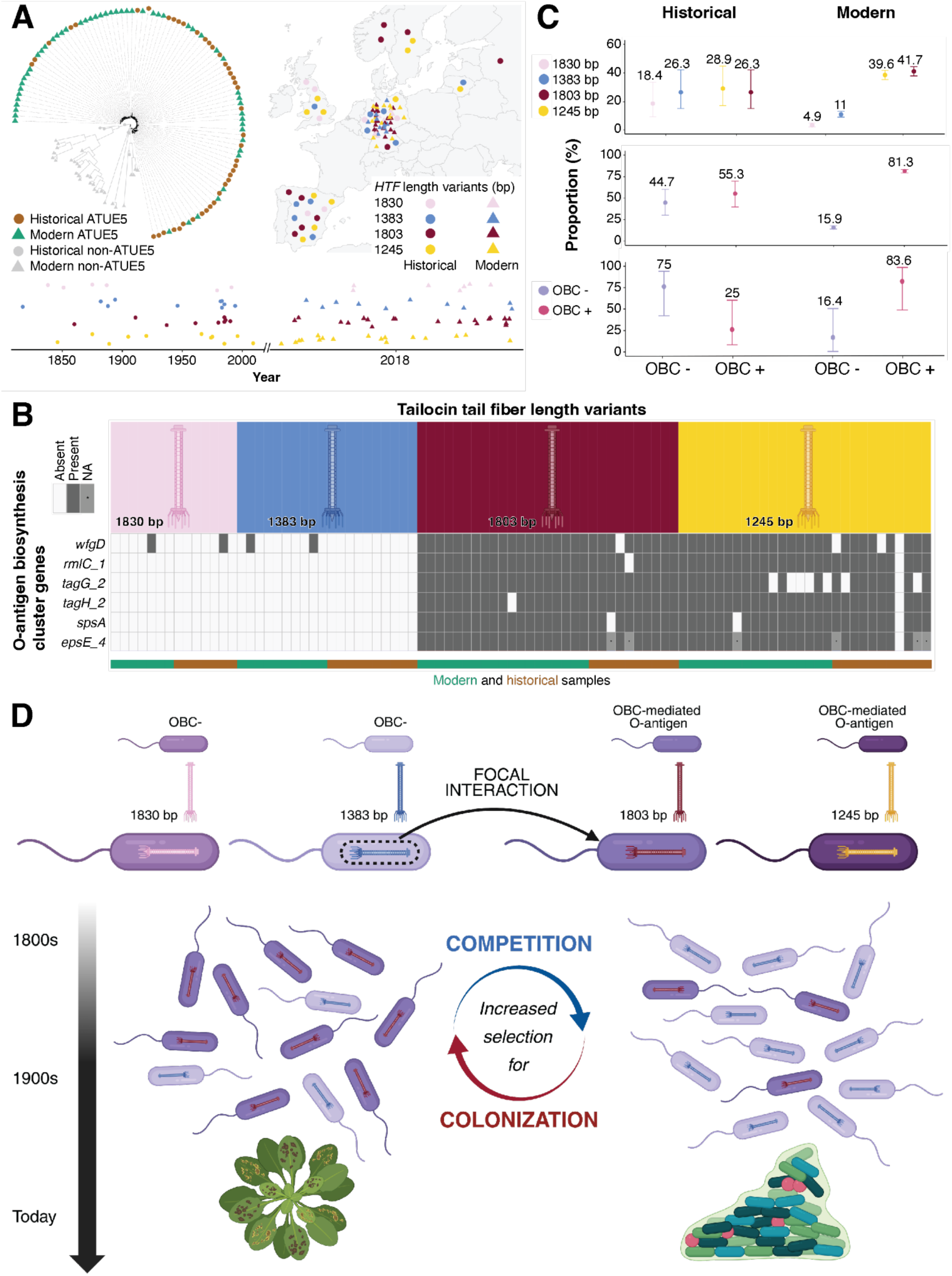
Century-long association between tailocin and O-antigen variants in *P. viridiflava*. **A.** Temporal and geographic distribution of historical and modern *Pseudomonas* genomes. The maximum-likelihood tree includes 83 modern *Pseudomonas* strains representing the known diversity segregating in wild populations, together with 49 historical herbarium-derived strains. The type of strain and its phylogenetic classification are indicated in the inset. The map shows the country of origin of each strain, with point locations placed arbitrarily within each country. The type of strain and its corresponding tail fiber (*HTF)* length variant are also indicated in the inset. The temporal distribution of historical and modern samples is shown below, with shape and color codes matching those in the map. Historical strains are positioned according to their collection dates, whereas a horizontal jitter was applied to modern samples (collected in 2018) for visualization. A random vertical jitter was applied to all samples for visual clarity. **B.** Patterns of presence and absence of six OBC genes and their association with *HTF* length variants in historical (brown) and modern (green) genomes. **C.** Proportion of *HTF* length variants and their association with the presence or absence of the O-antigen biosynthesis cluster (OBC). The upper panel shows the frequency of *HTF* length variants in historical and modern strains, the middle panel groups *HTF* length haplotypes according to their association with the presence or absence of the OBC, while the lower panel compares the distribution of *HTF* length variants between German historical strains (n = 8) and modern strains downsampled to the same sample size. **D.** Conceptual model of long-term HTF-O-antigen coevolution in natural *Pseudomonas* populations. This schematic illustrates how fluctuating selection between competitive and colonization contexts may maintain diversity in tailocin tailfiber (HTF) and O-antigen variants over time. Four major *HTF* length variants (1245 bp, 1383 bp, 1803 bp, and 1830 bp) co-occur in natural populations, each represented by a distinct color (yellow, blue, dark red, and pink, respectively). Strains with 1245 bp and 1803 bp *HTF* possess the OBC-mediated O-antigen, while 1383 bp and 1830 bp variants lack OBC genes, reflecting differences in tailocin susceptibility and colonization potential. The lower panel depicts how ecological conditions cyclically favor different strategies: under competitive conditions, strains producing tailocins with the 1383-bp *HTF* variant gain advantage, whereas under host-associated or colonization-favoring conditions, variants with the OBC-mediated O-antigen perform better. These opposing pressures have likely acted for at least 200 years, maintaining polymorphism in *HTF* length and associated O-antigen. Created in https://BioRender.com.

To place the 49 historical *Pseudomonas* draft genomes in the context of contemporary diversity, we constructed a phylogenetic tree that included a subset of 83 contemporary genomes representing the known diversity of *Pseudomonas* segregating in the wild ^7^. The tree topology revealed that 43 of the historical draft genomes belong to the ATUE5 phylotype and are broadly distributed across the ATUE5 phylogeny, indicating that historical diversity mirrors contemporary diversity and demonstrating the temporal continuity of ATUE5 over centuries (**Fig. 4A; Fig. S12**).

We then sought to determine the presence of the tailocin cluster and, when present, identify the corresponding *HTF* length variant. The tailocin cluster was detected in all 43 historical genomes (average coverage depth of 17.4x and average proportion of 86.0% of the cluster covered), supporting the idea of a single ancestral insertion shared across the entire ATUE5 population. Due to the high sequence divergence among *HTF* variants, a mapping-based approach was not feasible. We therefore identified the variants using local assembly for high-coverage samples and a unique *k*-mer approach for low-coverage samples. Both approaches yielded congruent results when tested on contemporary ATUE5 short-read data and validated against their corresponding whole-genome assemblies (**Fig. S13A**). Out of the 43 historical *P. viridiflava* draft genomes, we had enough coverage to retrieve reliable *HTF* variants for 38 of the genomes. The local assembly approach permitted the ascertainment of 23 historical *HTF* variants, whereas the approach based on unique *k*-mers allowed us to retrieve 15 additional *HTF* variants (**Fig. S13B**). Our analysis revealed that the four *HTF* length variants have been maintained in European populations over the past 200 years (**Fig. 4A**), suggesting that these variants are subject to long-term selective pressures that contribute to the stability of tailocin-mediated interactions.

Finally, we sought to determine whether the association between *HTF* length variants and O-antigen haplotypes has been maintained over century timescales. To address this, we annotated the presence or absence of six genes within the O-antigen biosynthesis cluster (OBC) — previously identified as key determinants of tailocin susceptibility in *P. viridiflava* strains ^7^. We found that the 1803 bp and 1245 bp *HTF* variants were almost exclusively associated with the presence of the six OBC genes (OBC^+^), with only one historical sample deviating from this pattern. Meanwhile, the 1830 bp and 1383 bp *HTF* variants were exclusively associated with the absence of OBC (OBC^-^) (**Fig. 4B**). The association between *HTF* variants and OBC haplotypes would be expected if the bacterial chromosome were in strong linkage disequilibrium (LD), but would be unexpected if horizontal gene transfer (HGT) were pervasive and routinely broke down long-range LD. To evaluate this, we calculated pairwise LD between ATUE5 single nucleotide polymorphisms (SNPs) and measured LD decay as a function of the genetic distance between SNPs. We found that LD decays to 50% of its maximum value after only 78 bp (**Fig. S14**). Using an orthogonal, mixture-model–based approach to estimate the relative contributions of recombination and mutation ^20^, we further estimate that, on average, 96% of SNPs in ATUE5 arise by recombination (**Fig. S15**). These results indicate that HGT-driven recombination is pervasive in ATUE5, making the strong association between *HTF* variants and OBC haplotypes particularly striking, given that the tailocin cluster and the OBC cluster are separated by 1.7 Mb. Our results thus demonstrate that the coupling between tailocin *HTF* length variants and O-antigen haplotypes in *P. viridiflava* has been maintained over a century timescale, likely reflecting the requirement for bacterial self-immunity to their own tailocins. This enduring association suggests that the trade-off between competitive ability and host colonization has persisted for centuries, driven by structural and molecular constraints that restrict the diversification of compatible *HTF*-O-antigen pairs. These constraints preserve functional specificity and prevent deleterious mismatches over evolutionary time. Contemporary data show that *HTF* length variants associated with the absence of O-antigen persist at low frequencies (**Fig. 4C**), likely reflecting the fitness cost imposed by the competition–colonization trade-off. Analysis of historical *HTF* length haplotype frequencies further reveals that none have approached fixation and that their relative abundances fluctuate over time in both the European and German populations (**Fig. 4A-C**), suggesting that the magnitude of the fitness cost associated with the trade-off may vary across ecological or evolutionary contexts.

## Discussion

Our results demonstrate that an O-antigen biosynthesis cluster (OBC) mediates a persistent trade-off between tailocin-dependent microbial competition and successful host colonization in *P. viridiflava*. OBC^+^ strains colonize *A. thaliana* more successfully but remain more vulnerable to tailocin killing, while OBC^-^strains resist tailocins but exhibit reduced growth *in planta*. Previous work has shown O-antigen-dependent costs of phage or tailocin resistance through experimental evolution ^21,22^. However, it remains unclear whether such constraints persist in natural populations over extended evolutionary timescales, where environments and clade abundances fluctuate. Here, we extend this body of work, using a time-series of herbarium-derived genomes, to show that this trade-off is present in natural populations, and long-lived, persisting for more than two centuries or hundreds of thousands of generations.

Tailocin tail fibers recognize O-antigen sugar motifs, with length variation modulating their breadth of killing. The persistence of linkage in OBC-tail fiber associations across both modern isolates and herbarium samples indicates that compensatory evolution to escape this trade-off is constrained, and if it occurs at all, is likely infrequent. It has been hypothesized that evolutionary systems can overcome trade-offs when conflicts are regulatory or ecological—when organisms evolve conditional expression, gene duplication, or context-dependent specialization that separates competing functions ^23^. In contrast, structural conflicts embedded in shared molecular architecture are predicted to be less flexible ^24^. The O-antigen exemplifies the latter: the same surface structure underlies both host colonization and susceptibility to tailocins, creating an antagonistic pleotropy that makes the trade-off difficult to decouple without compromising either function. We hypothesize that the long-term stability of OBC variants therefore illustrates a case where natural selection is persistently trapped by a structural constraint.

Similar constraints are seen in other systems, such as the conserved flagellin epitope flg22, where mutations that evade host recognition compromise essential motility ^25,26^. Our findings suggest that the costs of O-antigen-mediated resistance are lasting features of bacterial physiology rather than short-term fitness deficits that are easily overcome by compensatory mutations.

The continued presence of OBC⁻ strains in wild populations despite reduced colonization success suggests that ecological factors buffer the costs of O-antigen loss. OBC^-^ strains may rely on other surface glycans or exopolysaccharides or gain advantages in niches where competition outweighs colonization ^27^. Unlike host-microbe interactions in which frequency-dependent selection maintains resistance variation ^28^, in the tailocin system there is no known frequency dependence of the selection. Instead, variation in host genotype, community composition, or niche context may periodically favor OBC⁻ lineages, sustaining ATUE5 diversity alongside the structural constraint itself, because no single trade-off state maximizes fitness across all environments ^24^. Together, these findings show how enduring molecular trade-offs can be stabilized by ecological feedback, producing long-term coexistence rather than evolutionary escape. Understanding such stable constraints could inform antimicrobial strategies that align with, rather than oppose, the predictable evolutionary dynamics of bacterial populations.

Our conceptual model (**Fig. 4D**) synthesizes these results for long-term coevolution between tailocin tail fiber (*HTF*) length and O-antigen structure. *HTF* variants co-occur in wild *Pseudomonas* populations and are associated with differences in tailocin susceptibility and host colonization. We propose that fluctuating ecological contexts of competition versus colonization drive alternating selection for O-antigen variation. Under competitive conditions, tailocin-producing strains with reduced O-antigen gain the fitness advantage, whereas under host-associated contexts, O-antigenic variants perform better. Over time, these opposing forces maintain polymorphism in both *HTF* length and O-antigen composition. This dynamic balance may explain the remarkable temporal persistence of these variants and show how ecological feedback loops can stabilize structural diversity without requiring continuous molecular innovation.

## Methods

### Bacterial strains, plasmids and growth conditions

The strains used in this study are from Karasov et al. 2018 ^6^ (ENA: PRJEB24450). Bacteria were grown on Luria-Bertani broth agar (LB agar) and in Luria-Bertani (LB) medium at 28°C for *Pseudomonas spp.* strains and 37°C for *Escherichia coli* strains. All liquid cultures were incubated with shaking at 180 rpm. Antibiotics were used when appropriate at the following concentrations: 50, 100, and 10 μg/ml for kanamycin (Km), nitrofurantoin (NFN), and tetracycline (Tc), respectively. All strains were stored at −80°C in 30% glycerol [v/v]. For plasmid preparation, *E. coli* transformants were cultured in LB. For the selection of the second *P. viridiflava* homologous recombination event, LB supplemented with sucrose (10 g/L tryptone, 5 g/L yeast extract, 10% (W/V) filtered sucrose, 15 g/L agar) plates were used.

### Tailocin extraction and partial purification

Methods were adapted from Backman et al. 2024. Overnight cultures were back-diluted into 50 ml of fresh LB to extract and isolate tailocins from the *P. viridiflava* strains. When cultures reached an exponential growth phase [an optical density at 600 nm (OD600) of 0.4 to 0.6), 5 μg/ml MMC (Selleck, catalog no. S8146) was added to induce the bacterial SOS response and tailocin induction. The cultures were then incubated at 28°C for a minimum of 18 hours and then centrifuged at 4°C for 1 hour at 1,400×g to pellet cell debris. The supernatants were sterilized by filtration with 0.2-μm cellulose acetate filters. To precipitate tailocins, 40% (w/v) of ammonium sulfate was slowly added to the filter-sterilized tailocin lysate while stirring on ice. The lysate was left stirring on ice for at least 18 hours. After 18 hours at 4°C, the ammonium sulfate was pelleted by centrifugation at 2,090 × g for 2 hours at 4°C. The supernatant was discarded, and the pellet was resuspended in 500 μl of cold P-buffer (100 mM NaCl, 8 mM MgSO4, 50 mM Tris-HCl, pH 7.5) and stored at 4°C. Tailocin lysates were used fresh or for up to 2 weeks and stored at 4°C.

### Testing bacterial sensitivity to tailocins with spot test phenotypic assays

Methods were adapted from Backman et al. 2024. Briefly, to test the sensitivity of the different bacterial strains to the tailocins, soft agar assays were performed using an adaptation of a protocol from Vacheron et al.^29^. Overnight cultures (750 μl) of each strain were mixed with 25 ml of LB soft agar (0.8%), and the mixture was poured into a square plate and left to harden. Then, aliquots of 3 μl of concentrated tailocins suspension were applied to the agar, along with a control of P-buffer alone and noninduced cultures in serial dilutions. The plates were incubated overnight at 28°C. Bacterial sensitivity or resistance to the tailocins was assessed after 24 hours. No plaques were observed when testing serially diluted tailocins, suggesting that the killing agent is nonreplicative and not a phage. The p25.A12 tailocin can kill p25.C2 WT, and was used for mutant spot test phenotypic assays. There were three biological replicates per condition and eight serial dilutions.

### Plant infections of *Pseudomonas* natural isolates

*A. thaliana* genotypes Eyach 1.5-2 (15-2) and Col-0 were grown axenically and infected with single *Pseudomonas* strains as described in^6^. In brief, seeds were sterilized by overnight incubation at −80°C, followed by 4 hours of bleach treatment at room temperature (seeds in an open 2 mL tube in a desiccator containing a beaker with 40 mL Chlorox and 1 mL HCl (32%)). Seeds were then stratified for three days at 4°C in the dark on ½ MS media. Plants were grown in 3-4 mL ½ MS medium in six-well plates in long-day (16 hours) at 23°C. 12-14 days after stratification, plants were infected with single bacterial strains.

*Pseudomonas* overnight cultures were diluted at 1:10 in 5 mL of selective media and grown for an additional three hours. Bacteria were then centrifuged at 3500 g and the pellets diluted to an optical density (OD600) of 0.01 in 10 mM MgSO_4_. A hundred microliters of this bacterial suspension were used to drip-inoculate plants, distributing the volume over the whole rosette. Plants were infected with 10 mM MgSO_4_ as a control. The plates were sealed with parafilm and then returned to the growth chamber. Seven days after infection, pictures of rosettes were taken for green pixel quantification. At least three biological replicates were performed per infection.

### Plant health quantification with green pixels

For plant health quantification, plates were photographed seven days post-infection, with a tripod-mounted Canon PowerShot G12 digital camera. Individual plants were extracted from whole-plate images as described in ^14^. The number of green pixels was determined for each plant and used as a proxy for plant fresh mass. The segmentation of the plant from the background was performed by applying thresholds in LAB color space, followed by a series of morphological operations to remove noise and non-plant objects. Finally, a GrabCut-based postprocessing was applied, and CSV files with plant IDs and green pixel counts were created. The workflow was implemented in Python 3.6 and bash using OpenCV 3.1.0 and scikit-image 0.13.0 for image processing operations^14^.

### Competitive mutant fitness assays *in planta*

Columbia-0 (Col-0) and Eyach 1.5-2 WT *A. thaliana* plants were grown under long-day conditions (16 hours light, 8 hours dark) in an AR41L3 Percival detector with 60% intensity of the SciWhite LED lights. Seedlings were grown in 24-well plates (Greiner Bio-One, catalog no. 6621665). Thirteen-day-old seedlings were used for the infections. Plants were infected with bacterial suspension of the *P. viridiflava* p25.C2 saturation mutagenesis library at an OD_600_ of 0.02. Each plant was infected with 200 μL of bacterial suspension. Each biological sample contained two infected plants. Three days after infection, samples were collected in 2-mL deep-well plates and snap-frozen. Material was ground using two 5-mm glass beads and DNA was extracted using the Puregene extraction kit (QIAGEN Gmbh, Hilden, Germany).

### RB-TnSeq and analysis of RB-TnSeq data

Methods were adapted from Backman et al. 2024. Briefly, genomic DNA was extracted and barcode PCR was performed as described in^15^. DNA extractions were quantified with NanoDrop 1000 (ThermoFisher Scientific). Barcode sequence data were obtained by multiplexing on a partial NovaSeq X plus lane (Illumina, San Diego, United States) at Novogene (Novogene Corporation Inc., Sacramento, United States). Fitness data were calculated and analyzed from these reads with the DESeq2 R package^30^, and scripts can be found on our GitHub page.

### Construction of the p25.C2 *Pseudomonas* gene deletions

The p25.C2 mutant strains (Δ*OBC*, Δ*wfgD*, Δ*rmlC_1*, Δ*tagG_2*, Δ*tagH_2,* Δ*epsE_4*, and Δ*spsA*) were obtained using the Gateway cloning system with the donor vector pDONR1K18ms (Addgene plasmid no. 72644) and the destination vector pDEST2T18ms (Addgene plasmid no. 72647). Briefly, attB-flanked upstream and downstream gene fragments were ordered from TWIST^31^ (**Table S5**). BP and LR reactions were conducted in the one-tube format using the attB-flanked fragments. The resulting expression clone was transformed into competent *E. coli* DH5α and then introduced into recipient cells (*P. viridiflava* p25.C2) by conjugation. Recombinant cells (merodiploids) were selected for with the antibiotic resistance conferred by the expression clone and verified with PCR and gel electrophoresis. Positive colonies were purified by streaking onto new plates. Purified merodiploids were grown overnight without the antibiotic conferred by the expression clone to allow for another recombination event and then plated on 10% D-sucrose for *sacB* counterselection. Colonies were examined by PCR and verified by DNA sequencing (GeneWiz). The plasmids for *tagH_2* and *spsA* were ordered directly from TWIST.

### LPS isolation and purification

Bacteria were grown overnight at 28 °C under vigorous agitation (200 rpm). The suspensions were centrifuged, and the pellets were washed three times in PBS. Wet cell pastes were stored at -70 °C until LPS extraction. The cells were uniformly suspended in deionized water (ratio 1:5 w/v) and the suspensions were brought up to 68 °C with gentle stirring. The extraction was performed by adding an equal volume of preheated to 68 °C 90% liquefied phenol (w/w) and stirring for 20 min at 68 °C, following the Westphal procedure^32^ and as previously described^7^. Nucleic acids and proteins were removed by treatment with Benzonase (15 U/mL of LPS stock for 18 h, 37 °C, in the 50 mM MgCl_2_ and 20 mM NaOAc buffer at the pH 7.6), followed by Proteinase K (0.66U/ mL, 18 h, 37 °C), and dialysis (12-14,000 MWCO) at 4 °C against several exchanges of deionized water. Dialyzed fractions were freeze-dried, dissolved, and precipitated three times in cold (-20 °C) 90% EtOH. LPS pellets were freeze-dried again and dissolved in deionized water, and additionally purified by ultracentrifugation at 100,000 × g at 4 °C for 16 h. The enzymatic purification and the ultracentrifugation were repeated twice to obtain ultrapure LPS, which was verified by chemical analysis.

### Analysis of pure LPS by DOC-PAGE

One microgram of purified LPS samples in Laemmli buffer were resolved in 0.75mm-Polyacrylamide gel electrophoresis (PAGE, 4% stacking gel and 18% resolving gel) in the presence of deoxycholic acid (DOC) detergent^33^. for 1 h at 400 V, 30 mA. *Salmonella enterica* ser. *typhimurium* S-type LPS (O-antigen producing strain) was used as a standard. The PAGE was fixed overnight in an aqueous solution of 40% ethanol and 5% acetic acid, and the LPSs were visualized using a silver stain reagent kit (Bio-Rad, CA, USA) after oxidation with sodium periodate.

### Chemical analysis of pure LPS

Analysis of the glycosyl residues constituting LPS was achieved by derivatizing the samples to O-trimethylsilyl (TMS) methyl glycosides. Briefly, samples were methanolyzed with 1 M HCl-methanol at 80 °C for 18 h, re-N-acetylated at 100 °C for 1 h, and O-trimethylsilylated with Tri-Sil reagent (Thermo-Fisher) at 80 °C for 30 min. Each sample was supplemented with an internal standard of myo-inositol ^34,35^. GC-MS analysis of the LPS derivatives was performed on an Agilent AT 7890A GC system interfaced to a 5975B MSD using an Equity-1 (Supelco) fused silica capillary column (30 m length × 0.25 mm ID × 0.25 μm film thickness). The temperature gradient was 80 °C for 2 min, then increased to 140 °C at 20 °C/min with a 2-min hold, followed by an increase to 200 °C at 2 °C/min. Finally, the temperature was increased to 250 °C at 30 °C/min with a 5-min hold. The data were processed using Agilent ChemStation.

### Plant infection by flood inoculation

To evaluate the growth of p25.C2 WT and O-antigen mutants *in planta*, Columbia-0 and Eyach 1.5-2 WT *A. thaliana* plants were grown under long-day conditions (16 hours light, 8 hours dark) in an AR41L3 Percival detector with 60% intensity of the SciWhite LED lights. Seedlings were grown in 24-well plates (Greiner Bio-One, catalog no. 6621665). Thirteen-day-old seedlings were used for the infections. Plants were infected with bacterial suspension of the *P. viridiflava* strain p25.C2, O-antigen mutants, or buffer (10 mM MgSO4). All bacterial strains were tagged with luciferase^14^. Bacteria were grown overnight and diluted 1:10 on the day of the infection. Cells were grown for another 3 hours and then collected and resuspended in 10 mM MgSO4 to a final OD600 of 0.01. The plants were flood-inoculated for 1 min in a randomized manner with 1 mL of treatment (WT, mutant, or buffer).

### Bacterial fitness evaluation

After flood-inoculation, plants were grown for 7 days and then collected. The collected plants were ground in 1 ml of 10 mM MgSO_4_ using the Qiagen tissue lyser II, and 200 μl of the suspension was used to measure luciferase activity with a microplate plate reader (Spark®, TECAN, Switzerland). Plants infected with MgSO_4_ were used as controls. In total, 24 plants were used for each strain or buffer. Plants were blindly scored as healthy (only green, healthy tissue), diseased (chlorosis phenotype, may have some white tissue), or dead (necrosis phenotype, full white tissue).

To assess differences in bacterial growth across strains, we used linear mixed-effects models with batch as a random effect and strain as a fixed effect, using the lmer() function in the lme4 R package. Pairwise comparisons between WT and mutant strains were performed using the emmeans() function from the emmeans package. P-values were adjusted for multiple comparisons using the False Discovery Rate (FDR) method, accounting for all pairwise contrasts among strains. This approach enabled us to test for significant differences in colonization while controlling for inflated Type I error rates resulting from multiple tests.

### Oxidative stress tolerance assays

Methods were adapted from Binesse et al., 2015^36^. To assess sensitivity to reactive oxygen species, we measured growth of WT and O-antigen mutants across a range of hydrogen peroxide (H_2_O_2_) concentrations using a microplate growth assay. Overnight cultures were diluted into fresh medium and inoculated into 96-well plates containing 0–10 mM H_2_O_2_ in technical replicates. Optical density (OD_600_) was recorded every 15 minutes for 24 hours at 28°C using a microplate reader (Spark®, TECAN, Switzerland). Four biological replicates were done.

For each plate (replicate), raw OD_600_ time series were converted to growth area (AUC) using the trapezoidal rule over 0–24 h. AUC values for each strain were normalized to their respective 0 mM control to yield AUC_rel0._ To account for plate-to-plate variation in baseline H_2_O_2_ potency, we estimated a per-plate WT minimum inhibitory concentration (MIC), defined as the first concentration where mean AUC_rel0_ dropped below 0.1. All H_2_O_2_ concentrations on that plate were then scaled by the corresponding WT MIC to generate a standardized dose axis (H_2_O_2__scaled).

To test for differences in dose-response profiles between WT and O-antigen mutants, we fitted a linear mixed-effects model of log-transformed AUC values, log(AUC_rel0_) = strain × H_2_O_2__scaled + (1 | replicate), using the lmerTest package in R. Estimated marginal means (EMMs) and pairwise contrasts versus WT were computed with the emmeans package on the response scale.

To directly compare MIC values, we also estimated a strain-specific MIC for each replicate using linear interpolation of AUC_rel0_ versus H_2_O_2_ concentration. A second mixed model of log-transformed MIC values, log(MIC) = strain + (1 ∣ replicate), was used to test for differences in inhibitory thresholds.

### Disk diffusion assays

Bacteria were grown for 16 h at 28 °C with vigorous agitation (200 rpm). Suspensions were back-diluted 1:10 in 5 mL of selective liquid medium and grown for an additional 3 h. Suspensions were adjusted to OD600nm = 0.6 and streaked as a thin, uniform layer onto selective solid medium using sterile cotton swabs. Sterile 6 mm filter paper disks were saturated with 20 uL of antimicrobial compound solutions (polymyxin B or colistin sulfate) prepared at concentrations ranging from 0 to 500 ug/mL. Once dried, the disks were placed on the bacterial layer, ensuring sufficient distance between them to prevent overlap of inhibition zones. Inhibition radii were measured after overnight static incubation at 28 °C. The absence of significant differences among strains at each compound concentration was assessed using either one-way ANOVA or Kruskal–Wallis tests, as appropriate.

### Aggregation formation assay

Aggregation formation was assessed using a 96-well plate growth assay. Overnight cultures of each strain were grown in LB medium at 28 °C with shaking. The following day, cultures were back-diluted 1:10 in fresh LB and grown to an OD_600_ of approximately 0.5. Each culture was then diluted 1:40, and 200 µL of the resulting suspension was added to wells of a sterile, flat-bottom polystyrene 96-well plate (≥10 technical replicates per strain). Wells containing 200 µL of sterile LB served as negative controls. Plates were incubated statically at 28 °C for 48 h. Biofilm formation assays were done in three biological replicates.

After incubation, wells were visually blindly scored as aggregation (visible surface-associated growth), no aggregation (turbid growth without surface attachment), or clear (no growth). For statistical analyses, aggregation presence/absence was coded as a binary trait. Proportional differences among strains were first tested with a Pearson’s chi-square test, followed by pairwise Fisher’s exact tests comparing each mutant to the WT strain (p25.C2). Resulting p-values were adjusted for multiple testing using the Benjamini-Hochberg procedure.

### Immune gene expression assessment

Following flood-inoculation, rosettes were collected at 0, 2, 6, 10, 24, 48 and 72 hours post-infection. For each treatment modality at each time point, three individual plants were pooled to form one biological replicate, and this was repeated 3 times to obtain the biological replicates. Samples were snap frozen immediately after collection and stored at -80 °C until RNA extraction. Frozen material was ground using the Qiagen tissue lyser II and total RNA was extracted with TRIzol™ reagent (#15596026; Invitrogen, Thermo Fisher Scientific, MA, USA). RNA concentration, purity and integrity were verified spectrophotometrically and by gel electrophoresis. One microgram of RNA was treated with 1 unit of DNase I (#EN052; Thermo Fisher Scientific, MA, USA) and reverse-transcribed using 50 ng of random hexamers and 200 units of SuperScript™ IV (#18090200; Thermo Fisher Scientific, MA, USA). The resulting cDNA was treated with 2 units of *E. coli* RNase H (#18090200; Thermo Fisher Scientific, MA, USA) and diluted 1:20 to serve as a template for *PP2A* amplification (27 cycles), confirming successful cDNA synthesis. Samples were diluted 1:50 and used as templates for qPCR reactions performed with Luna® Universal qPCR Master Mix (New England Biolabs, MA, USA), using 20% of cDNA per reaction. All samples from one biological replicate were run on a single 384-well qPCR plate, with three technical replicates per target gene. Amplification was performed on a CFX Opus 384 Real-time PCR system (**Table S6**) and melt curves quality was assessed using the Maestro CX software (Bio-Rad, CA, USA). Amplification data were analyzed in RStudio using the chipPCR and qpcR packages. PCR efficiencies were calculated for each individual well and incorporated into the computation of mean normalized expression^37,38^. Expression of *FRK1*, *WRKY29*, *NHL10*, *PAD3*, *PDF1.2*, and *PR1* was quantified over time relative to the housekeeping gene *PP2A*, and normalized to the mock-treated samples at each time point. For each target gene, significant differences between the WT and O-antigen mutants at each time point were assessed using one-way ANOVA or Kruskal–Wallis tests, as appropriate.

### DNA extraction, library preparation and sequencing of *A. thaliana* herbarium specimens

Herbarium specimens of *Arabidopsis thaliana* spanning almost two centuries (between 1817 and 2015) were obtained from seven European institutions: Staatliches Museum für Naturkunde Stuttgart (Stuttgart, Germany), the Herbarium Tubingense (Tübingen, Germany), Lund University Botanical Museum (Lund, Sweden), the herbarium at the Biological Museum Oskarshamn (Oskarshamn, Sweden), the herbarium at Real Jardín Botánico (Madrid, Spain), the Herbario de Málaga (Malaga, Spain) and The Natural History Museum’s herbarium (London, United Kingdom). Small tissue samples (∼0.7 cm^2^) were collected for DNA extraction and sequencing, with care taken to minimize visible damage. Each sample specimen was labelled with collection details and internal identifiers, and DNA extraction followed established protocols^17^.

DNA from herbarium specimens was extracted and processed into single-stranded Illumina libraries at the UCSC Ancient and Degraded DNA Processing Center following Kapp et al. ^39^ with suggested modification from Nguyen et al. ^40^. Strict contamination control and cleanroom procedures were implemented. Libraries were initially shallow-sequenced on an Illumina NextSeq 550 to assess DNA quality and plant endogenous DNA content, then sequenced on a NovaSeq X Plus (2x150 bp), while adjusting per-sample concentration in the DNA pools to finally achieve an average depth around 9x for the host *A. thaliana* genome of each sample. All sequencing reads have been deposited in the European Nucleotide Archive (ENA) under project ID: PRJEB98841.

### Historical reads processing, mapping and authentication

We identified *Pseudomonas*-derived reads in 49 *A. thaliana* herbarium specimens collected across Europe between 1817 and 2015, comprising two previously published datasets ^41,42^ and 31 newly sequenced herbarium samples from this study (**Table S7**). Before read mapping, we used AdapterRemoval v2.3.3 ^43^ to trim adapters, remove low-quality bases, and collapse overlapping paired-end reads. Host-derived reads were removed by mapping all merged reads to the *A. thaliana* TAIR10 reference genome ^44^ using BWA aln v0.7.17 ^45^, with the seed disabled to improve alignment of damaged historical reads ^17^. For each library, reads that could not be merged due to long insert sizes were mapped independently as forward and reverse reads. Subsequently, merged and unmerged reads were combined into a single BAM file per library. The remaining unmapped reads were then mapped to the *P. viridiflava* ATUE5 p25.C2 reference genome^6^ using the same approach. PCR duplicates were removed, and mapped reads were filtered for mapping quality ≥ 20 using samtools v1.11^46^. To authenticate the historical nature of both *A. thaliana*- and *Pseudomonas*-mapped reads we used mapDamage2 v2.20 ^47^, which generated misincorporation patterns and length distributions typical of ancient DNA.

### Phylogenetic placement of historical *Pseudomonas* in the context of modern diversity

To determine the phylogenetic placement of the 49 historical *Pseudomonas* strains in the context of modern diversity, we used a set of 83 contemporary genomes representing the known diversity of *Pseudomonas* segregating in wild populations. Single Nucleotide Polymorphisms (SNPs) were identified using bcftools v1.11 with the parameter -ploidy 1 ^46^, filtered for quality (QUAL >= 20), merged across samples, and restricted to biallelic sites with no missing data. A maximum-likelihood tree was constructed from 3,095 biallelic SNPs detected in at least 95% of isolates using IQ-TREE v2.1.4 ^48^ applying TVM+F+ASC+R3 as the best substitution model ^49^. Historical strains clustering within the modern ATUE5 clade were classified as ATUE5. To examine the diversity within the ATUE5 lineage, we constructed a phylogeny using 53 modern and 43 historical ATUE5 genomes. SNPs were identified and filtered as described above, yielding a total of 8,184 biallelic SNPs. A maximum likelihood tree was constructed also using IQ-TREE v2.1.4 ^48^ applying TVM+F+ASC+R4 as the best substitution model ^49^.

### Ascertainment of tail fiber assembly (TFA) and hypothetical tail fiber (HTF) haplotypes in historical samples

To capture reads representing the full spectrum of tailocin genetic diversity, we extended the *P. viridiflava* p25.C2 reference genome with six major haplotypes of the highly polymorphic tail fiber (*HTF*) and tail fiber assembly (*TFA*) genes ^7^. Historical *P. viridiflava* reads were then mapped to this extended reference as described above. Reads mapping to the tailocin cluster between the flanking genes *trpE* and *trpG*, as well as all reads mapping to any haplotype of the *HTF* and *TFA* genes were used for *de novo* assembly using SPAdes v3.15.0 ^50^. Collinearity between the resulting assemblies and the reference *HTF* and *TFA* genes was assessed using Minimap2 v2.28 ^50,51^, and the best-matching *HTF* and *TFA* variants for each assembly were identified by ranking the proportion of coverage obtained from the Minimap2 mapping. To determine *HTF* haplotypes in historical strains that had insufficient coverage for *de novo* assembly of the *HTF* gene, we performed *k*-mer–based classification of *HTF* sequences. Reference sets of unique *k*-mers (k = 31) were constructed from the seven *HTF* variants (**Fig. S16**). Only *k*-mers exclusive to any of the *HTF* haplotypes and absent from whole-genome backgrounds were retained. To enhance the discriminatory power of our analysis, we filtered *k*-mers across haplotypes to ensure a minimum edit distance of ≥2 between them. The only exception was the haplotype with the smallest *k*-mer set (HTF_p23.B8), for which *k*-mers were removed only from the larger overlapping sets. Using this set of filtered *HTF* haplotype *k*-mers, we queried *k*-mer match counts for each historical isolate with Jellyfish v2.2.10 ^52^ (**Fig. S17**) (all additional *k*-mer profiles are available on GitHub). Additionally, we removed *k*-mers with a fold coverage less than the genome-wide mean minus 0.5 standard deviations. For each isolate, we calculated the proportion of *k*-mers matching each haplotype and assigned the haplotype with the highest proportion as the dominant one (**Table S8**). In cases where multiple haplotypes displayed similar dominant proportions, potential co-infections were recorded and confirmed through inspection of genome-wide *k*-mer profiles (**Fig. S17**). Modern high-quality reads were used as benchmarks to validate the *k*-mer approach and compare it with the local assembly method (**Fig. S13**).

### O-antigen biosynthesis cluster (OBC) gene presence/absence analysis

We determined the presence of six O-antigen biosynthesis genes (*wfgD, rmlC_1, tagG_2, tagH_2, spsA,* and *epsE_4*) ^7^. A gene was considered present if it met both of the following criteria: (i) a breadth of coverage ≥ 65% and (ii) a mean depth ≥ the genome-wide average minus 0.25 standard deviations of each isolate; otherwise, it was considered absent.

The *epsE_4* locus required additional analysis because of its extended length (∼4.5 kb) and high allelic divergence (**Table S8; see GitHub**). For a fraction of modern strains, the mapping to the *P. viridiflava* p25.C2 *epsE_4* reference resulted in the full recovery of the *epsE_4* gene. In other modern strains, we resorted in the previously published de novo assemblies ^7^ to extend from the conserved mapped region on the assembled contig. This process generated a set of diverse modern *epsE_4* variants. For historical isolates lacking whole-genome assemblies, the modern haplotype set was used as a reference for local assembly. Reads mapping to any variant were extracted and assembled locally using SPAdes v3.15.0. ^50^. Assembled contigs were then aligned against the modern haplotypes with Minimap2 v2.28 ^51^ to identify the best-matching variant.

For both modern and historical datasets, rescued *espE4* sequences were translated into amino acids using ExPASy ^53^ and EMBOSS Transeq ^54^. Amino acid sequences were then aligned with Clustal Omega ^55^. The resulting multiple sequence alignments and allele length distributions are provided in the supplementary materials (**Table S8; see GitHub**).

### Linkage disequilibrium and recombination analysis

To evaluate the impact of recombination on the association between *HTF* length variants and OBC haplotypes, we quantified genome-wide linkage disequilibrium (LD) and recombination across *Pseudomonas viridiflava* ATUE5. Variant calling was performed on 53 modern ATUE5 genomes using bcftools v1.11 ^46^, following the same procedure used for phylogenetic reconstruction. A total of 298,977 SNPs were retained and reformatted to include only the genotype (GT) field for LD estimation. Pairwise LD (r²) was computed using VCFtools v0.1.17 with the --hap-r2 function ^56^. LD decay was calculated for SNP pairs separated by 0–2 kb using all SNPs, whereas for 2–2000 kb using a random subset of 2% to reduce computation time. The median *r²* values were calculated after binning SNP pairs by distance (5 bp bins for short-range and 10 kb bins for long-range intervals).

To further quantify genome-wide recombination, the same 53 modern genomes were analyzed using Recophy ^20^ with the *P. viridiflava* p25.C2 reference genome under default settings. Recophy mixture-model–based approach to estimate whether SNPs across all pairwise genome comparisons arise via mutation (clonal fraction) or recombination (recombining fraction), from which the ratio of recombined to clonal SNPs was calculated.

### Plant infection by syringe-infiltration and bacterial population size assessment

Bacteria were grown for 16 h at 28 °C with vigorous agitation (200 rpm). Suspensions were back-diluted 1:10 in 5 mL of selective liquid medium and incubated for an additional 3 h. Cells were centrifuged and pellets were washed 3 times with 10 mM MgSO4 (5 mL). Suspensions were adjusted to OD600nm = 0.1 (10^8 CFU/mL), 0.01 (10^7 CFU/mL) and 0.001 (10^6 CFU/mL). Ten-week-old *Arabidopsis thaliana* plants of the Eyach 1.5-2 and Col-0 ecotypes, grown under long-day conditions (16 h light, constant 23 °C), were used for the infiltration, selecting two leaves at the same phenological stage from 3 individual plants for each treatment. Using a tip-less 1mL-syringe, half of each leaf was infiltrated on the abaxial face with either mock solution (10 mM MgSO4) or bacterial suspension (final volume = *ca.* 5-15 uL). Infiltrated plants were returned to the growth chamber and incubated under a transparent plastic lid to maintain high humidity. At two and five days post-infiltration, infiltrated leaves were collected and ground in 1 mL of 10 mM MgSO₄. Homogenates were serially diluted (five 1:10 dilutions), and three 10 µL drops of each dilution were spotted onto selective solid medium. After 48 h of static incubation at 28 °C, colonies were counted in each drop to estimate bacterial population size. Significant differences between the WT and O-antigen mutant for each condition were assessed using a Student’s t-test, Welch’s t-test, or Wilcoxon rank-sum test, as appropriate.

### Bacterial growth *in vitro*

Bacteria were grown overnight at 28 °C with vigorous agitation (200 rpm). Cells were centrifuged and overnight pellets were washed three times with PBS. Suspensions were adjusted to OD600nm = 0.01 in liquid LB. Two hundred microliters of suspensions were aliquoted in a 96-well flat transparent plate, considering 4 technical replicates per strain. LB media with no inoculum (sterile) provided the reference for optical density measurements The plate was placed in a humidity cassette and then loaded into the microplate reader (Spark®, TECAN, Switzerland) where it was continuously shaken and incubated at 28°C, and optical density at 600 nm was measured every 15 minutes for 24 h. Statistical analysis was performed with three independent biological replicates. Growth curve parameters were extracted using the growthcurver R package and differences were assessed by one-way analysis of variance (ANOVA1) followed by Tukey’s post hoc test for each parameter.

### Contributions

TB, JC, EC, MH, TLK and HAB devised the study. IB and MN developed the pipeline for analyzing plant growth. TB, EC, EB, ES, PG and AP performed plant growth and tailocin treatment experiments. TB, JC, EC, SL, TLK and HAB analyzed the data. SL, LL, JL, JE, GM, HAB and PL generated and provided the historical genomes data. EC, PA and AM designed the LPS assays and conducted the characterization. AH developed the RB-TnSeq pipeline. TB, JC, EC, MH, TLK and HAB wrote the paper.

## Acknowledgements

Haim Ashkenazy and Detlef Weigel for discussions and for providing access to the genome data. We are also grateful to the Max Planck Society for supporting the infection experiments, and to Jiyeon Hyun and Heejin Yoo for providing the primers used for PR1 amplification in qPCR. We thank Dominique Bergmann (Stanford) for hosting and supporting PL and JE’s generation of historical *A. thaliana* genomes. JRL and some herbarium sequencing was supported by NIH R35GM138300. We thank Aida Andrés, Gemma Murray and members of their labs for input in data analyses and valuable discussions. We further thank Joy Bergelson, Sophien Kamoun, Johannes Krause, Dmitri Petrov and Max Reuter for valuable discussions. This work would not be possible without historical herbarium collections and their curators. We are grateful to Anette Rosenbauer and Mike Thiv at the Staatliches Museum für Naturkunde Stuttgart (Stuttgart, Germany), Oliver Bossdorf and Uta Grünert at the Herbarium Tubingense (Tübingen, Germany), Arne Thell and Ulf Arup from Lund University Botanical Museum (Lund, Sweden), Åke Rühling at the herbarium at the Biological Museum Oskarshamn (Oskarshamn, Sweden), Leopoldo Medina from the herbarium at Real Jardín Botánico (Madrid, Spain), Jose García Sánchez at the Herbario de Málaga (Malaga, Spain) and John Hunnex, Fred Rumsey and Mark Carine from The Natural History Museum’s herbarium (London, United Kingdom) for their generous specimen loans and the kind permission to sample specimens. We are further grateful for all help received over the last five years from the California Academy of Sciences Botany Department staff, and for their support with obtaining loans from numerous international institutions, in particular to Collection Manager Emily Magnaghi, past Collection Manager Debra Trock, and past McAllister Chair of Botany Nathalie Nagalingum.

## Data and materials availability

All data and code are available in the supplementary materials and have been deposited in the following repositories. Historical sequencing read data have been deposited in the NCBI Sequence Read Archive under BioProject accession PRJEB98841 and in two public datasets ^41,42^. Code has been deposited to GitHub: (https://github.com/JiajunCui-jjc/HTF_OBC_historical_analysis.git, https://github.com/talia-backman/Ps1524_tailocin_tradeoffs).

---

**Figure S1:**
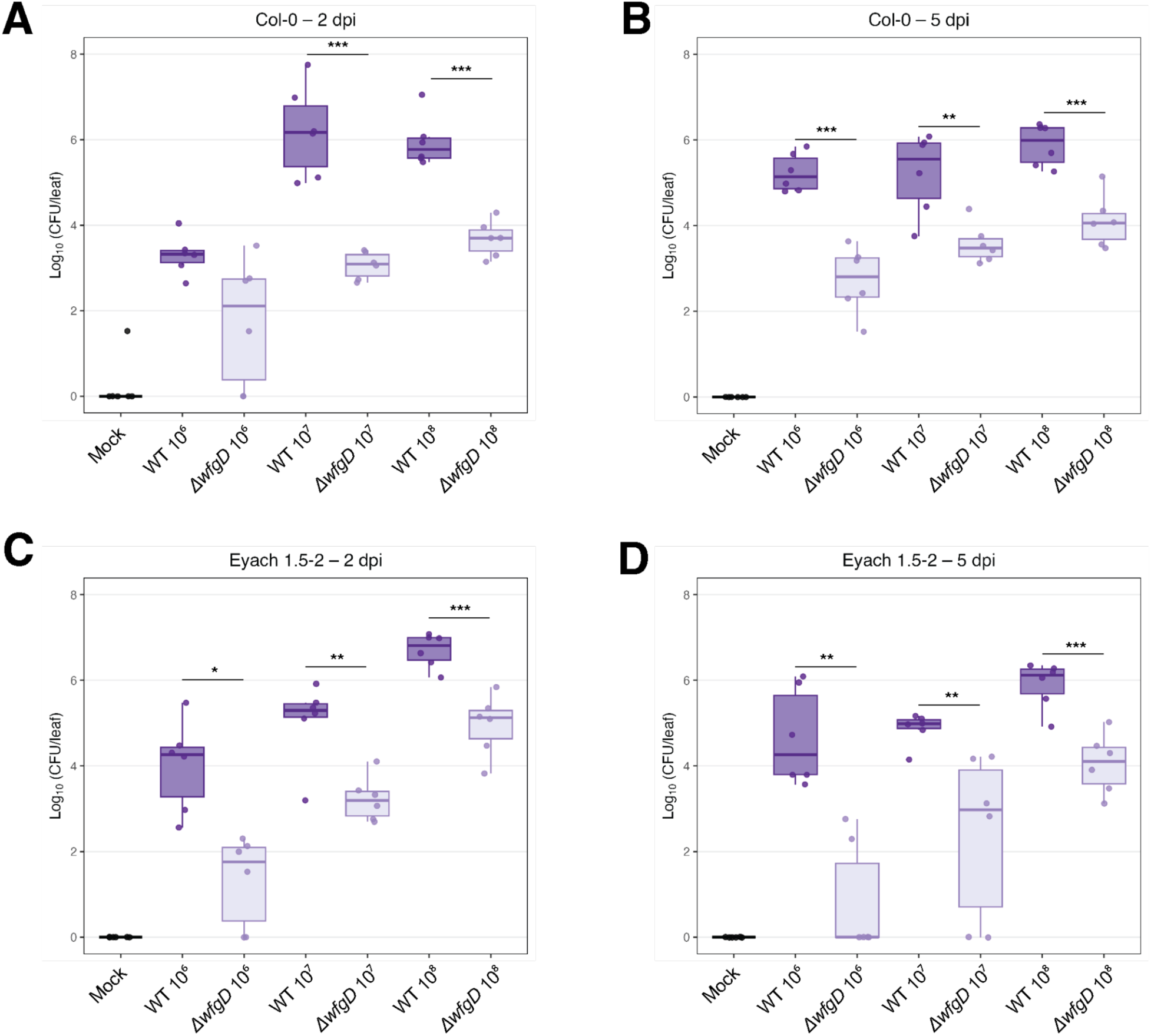
Bacterial population size of p25.C2 WT and O-antigen mutant *ΔwfgD* in Col-0 and Eyach 1.5-2 at 2 and 5 days post-infiltration (dpi) at different inoculum concentrations. The population size of the WT strain was significantly smaller than that of the *ΔwfgD* mutant in all but one condition. Differences between the two strains were assessed using a Student’s t-test, Welch’s t-test, or Wilcoxon rank-sum test, as appropriate (p-values: *** < 0.001, ** < 0.01, * < 0.05). Data represent 3 technical replicates per condition across 6 biological replicates.

**Figure S2:**
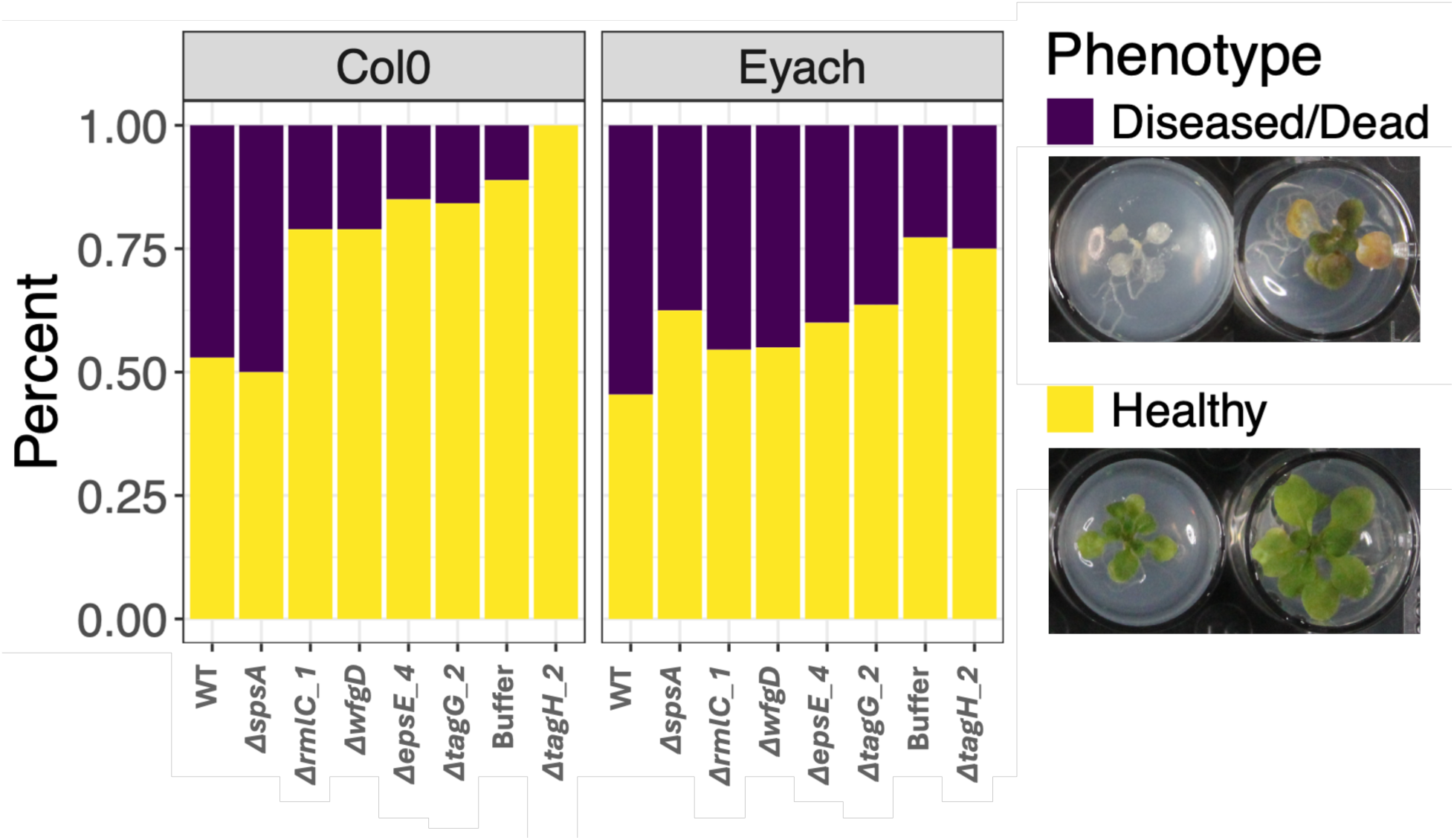
Proportion of plants exhibiting diseased/dead (purple) or healthy (yellow) phenotypes 7 dpi. Across both ecotypes, O-antigen mutants tended to show a higher proportion of healthy plants than WT, although differences were not statistically significant after multiple testing corrections. WT caused disease more frequently but not uniformly, indicating variability in pathogenic outcomes or flood inoculation. Photos of representative diseased, dead, and healthy plants are shown below the legend. Each well is 1.5 cm across.

**Figure S3:**
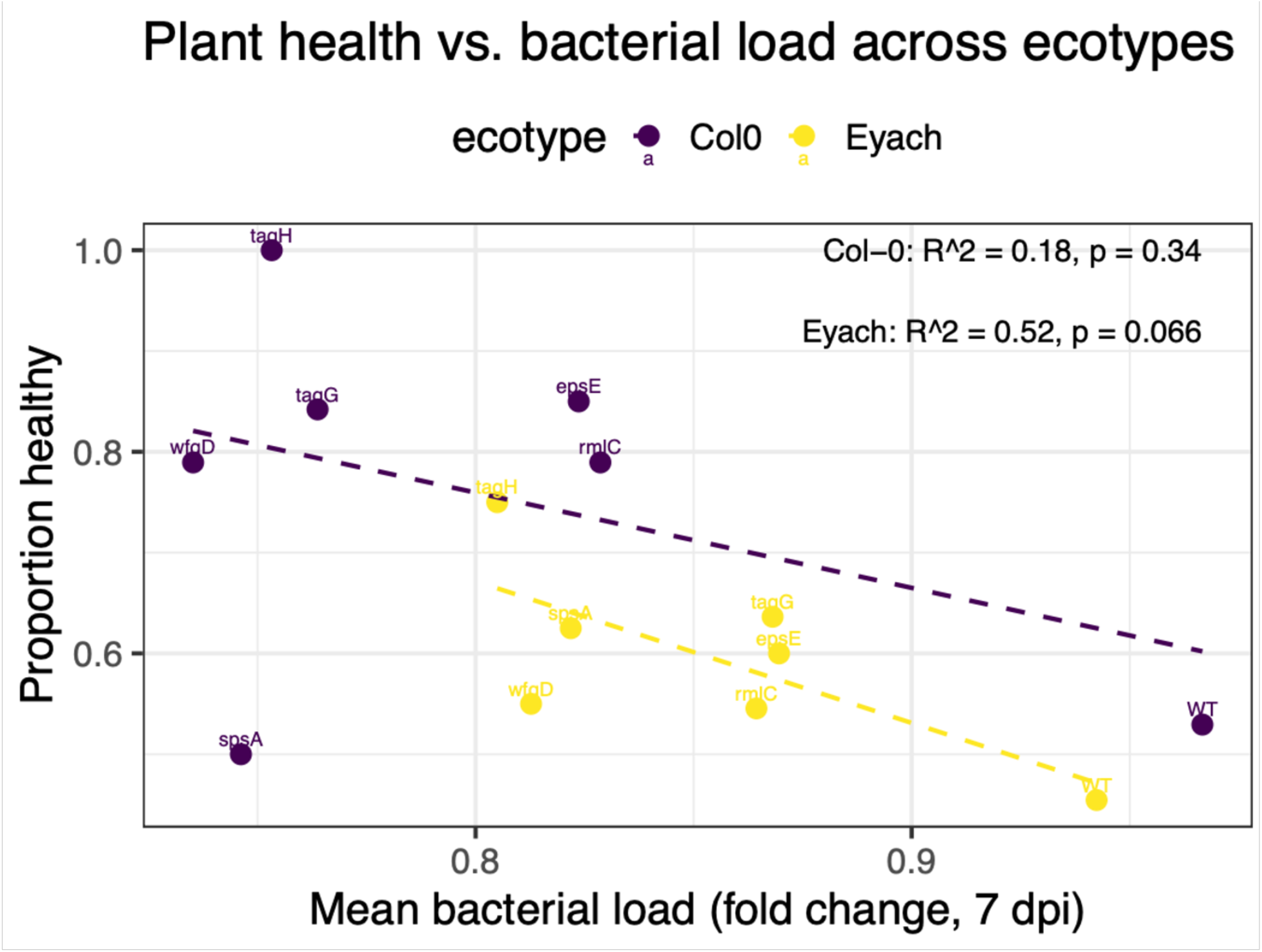
Mean bacterial load (x-axis) is negatively associated with plant health (y-axis) across strains. Each point represents a strain’s average 7 dpi fold change (from Fig. 4C) and the corresponding proportion of healthy plants (from Fig. S2). Linear regressions are shown for each ecotype. Across all strains, higher bacterial load tended to predict reduced plant health (pooled model R^2^ = 0.45, p = 0.037; slope p = 0.086), although this trend was not significant within Col-0 (R^2^ = 0.18, p = 0.34) and only slightly in Eyach 1.5-2 (R^2^ = 0.52, p = 0.066), reflecting background-specific variation in the relationship between colonization and disease severity.

**Figure S4:**
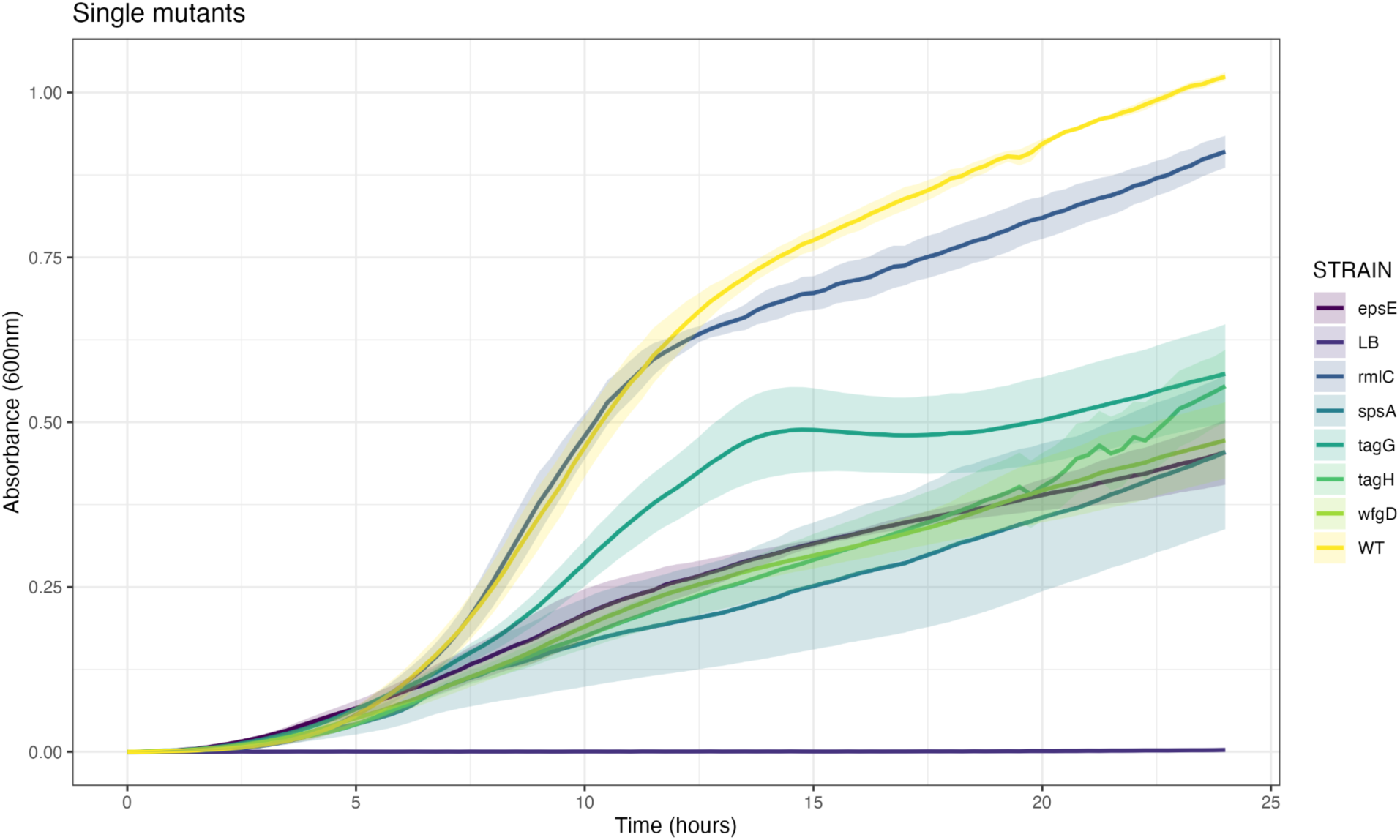
Growth curves *in vitro* of p25.C2 (WT) and the O-antigen mutants. Overnight cultures were adjusted to OD600nm = 0.01 in rich medium (LB) and absorbance was measured for 24 h. Data represent 4 technical replicates per strain across 3 independent biological replicates. Significant differences among strains are reported in Table S3.

**Figure S5:**
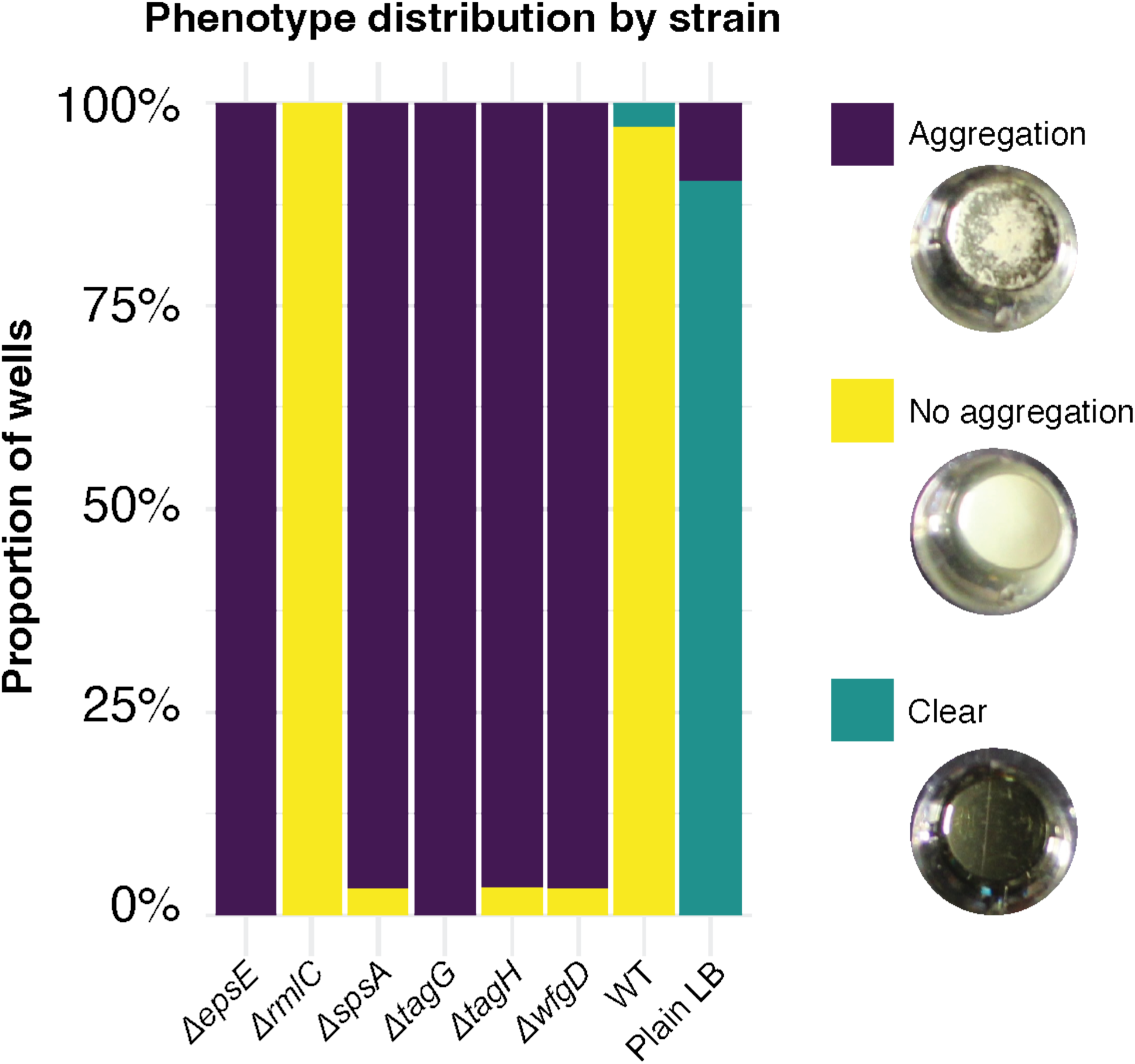
Proportion of wells displaying visible aggregation phenotypes after 48 hours of static incubation at 28°C. Wells were categorized as clear (no growth), growth with no aggregation, or growth with aggregation. All O-antigen mutants except *ΔrmlC* showed significantly higher aggregation frequency than the WT strain (p25.C2; Fisher’s exact test, Benjamini–Hochberg-adjusted p < 10^-16^). Data represent 10 technical replicates per strain across 3 biological replicates. Representative photographs illustrate each phenotype category: clear, growth without aggregation, and growth with aggregation.

**Figure S6:**
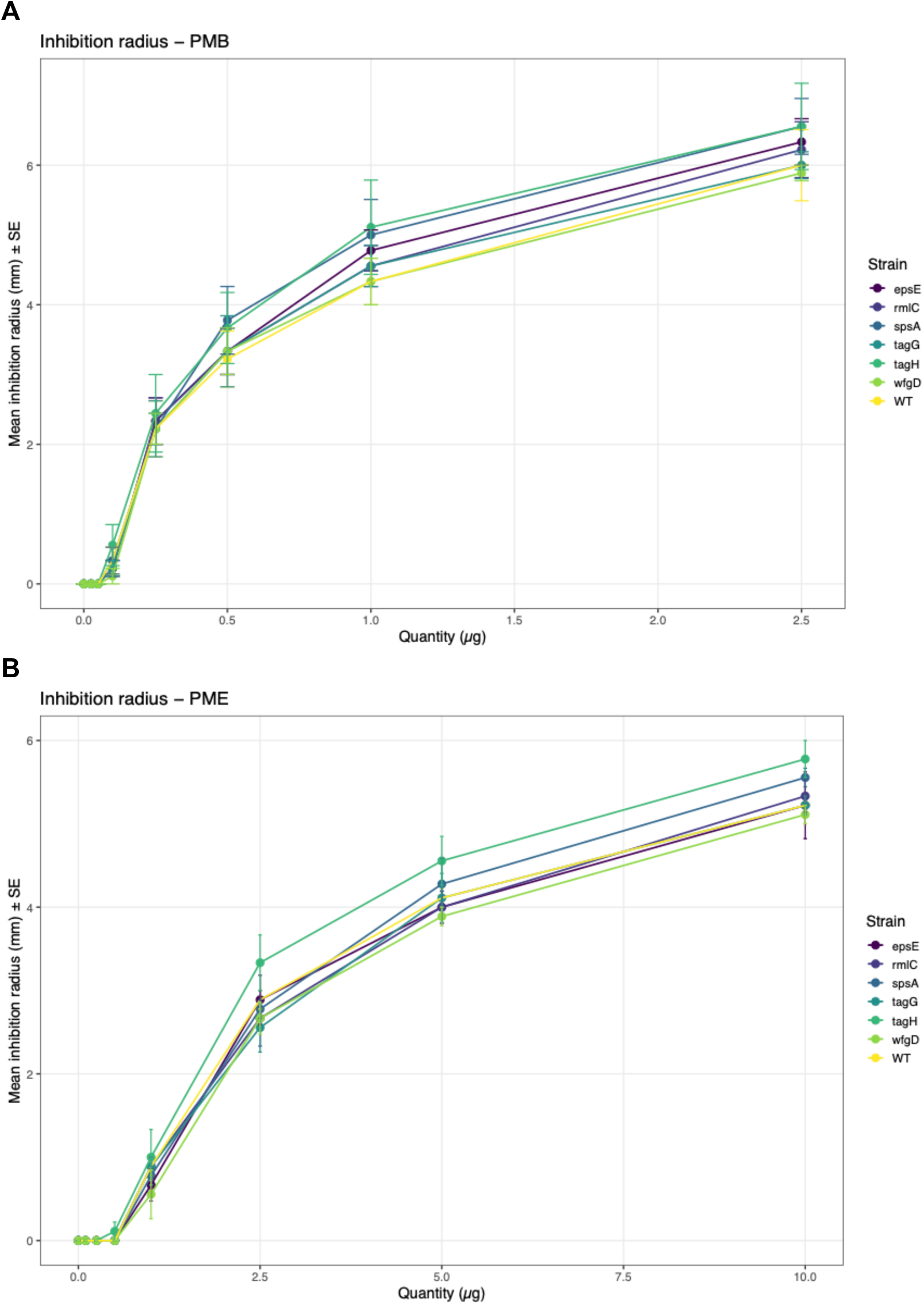
Sensitivity to polymyxin B (A) and polymyxin E (B) in p25.C2 (WT) and the O-antigen mutant. Filter paper disks were saturated with varying quantities of each polymyxin and placed onto freshly streaked bacterial lawns on rich agar medium (LB). Inhibition radii were measured after overnight static incubation at 28 °C. Data represent 3 technical replicates per condition across 3 biological replicates. The absence of significant difference among strains at each compound concentration was assessed using one-way ANOVA or Kruskal–Wallis tests, as appropriate.

**Figure S7:**
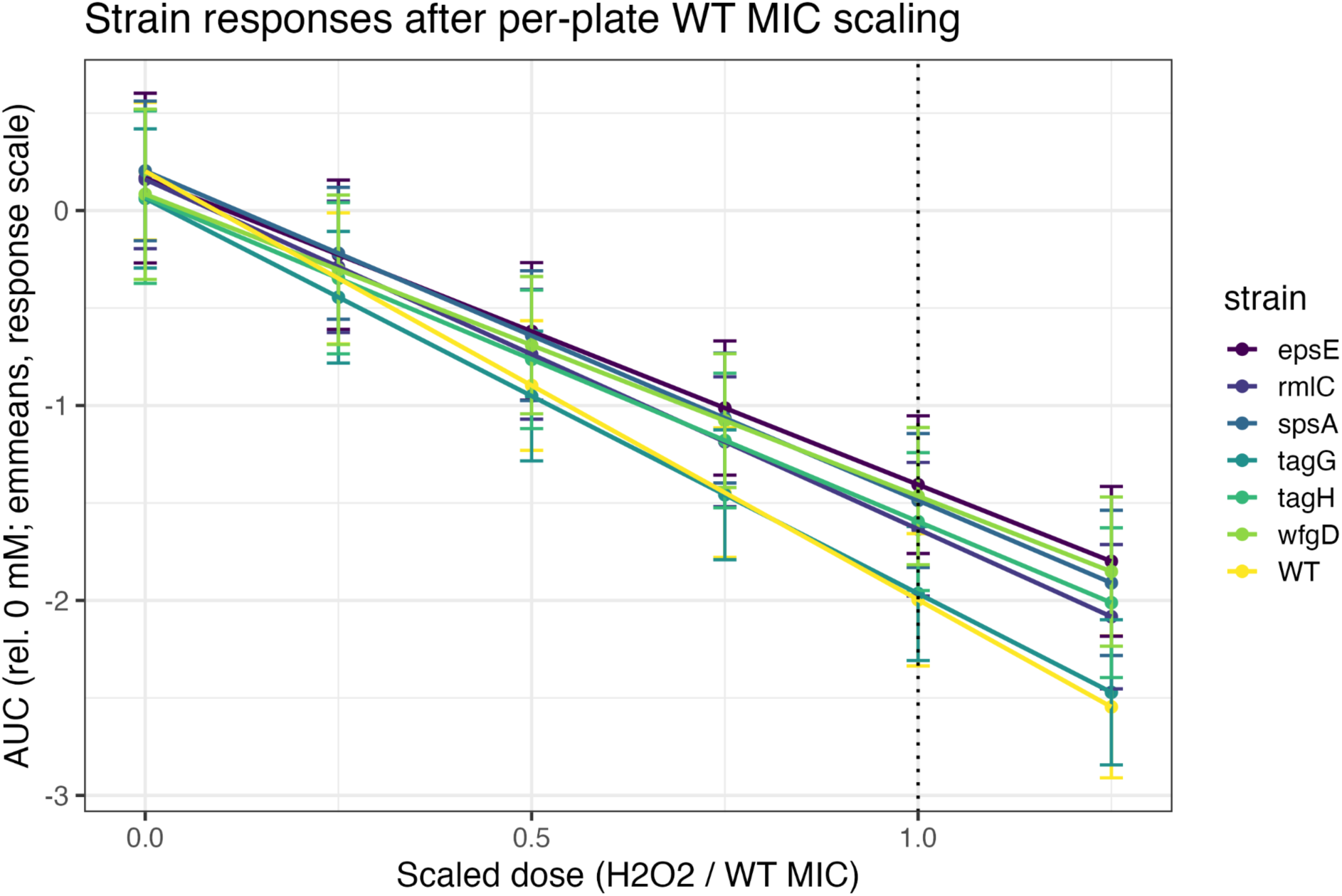
O-antigen mutants and WT exhibit equivalent tolerance to oxidative stress. Area-under-the-curve (AUC) values from growth curves at increasing H_2_O_2_ concentrations were normalized to 0 mM for each strain and replicate. Each plate’s doses were scaled by its WT minimum inhibitory concentration (MIC) to correct for between-experiment variation. Mixed-effects modeling with emmeans contrasts revealed no significant differences between any OPS mutant and WT (p > 0.05). Lines show estimated marginal means ± 95% confidence intervals from four replicates.

**Figure S8:**
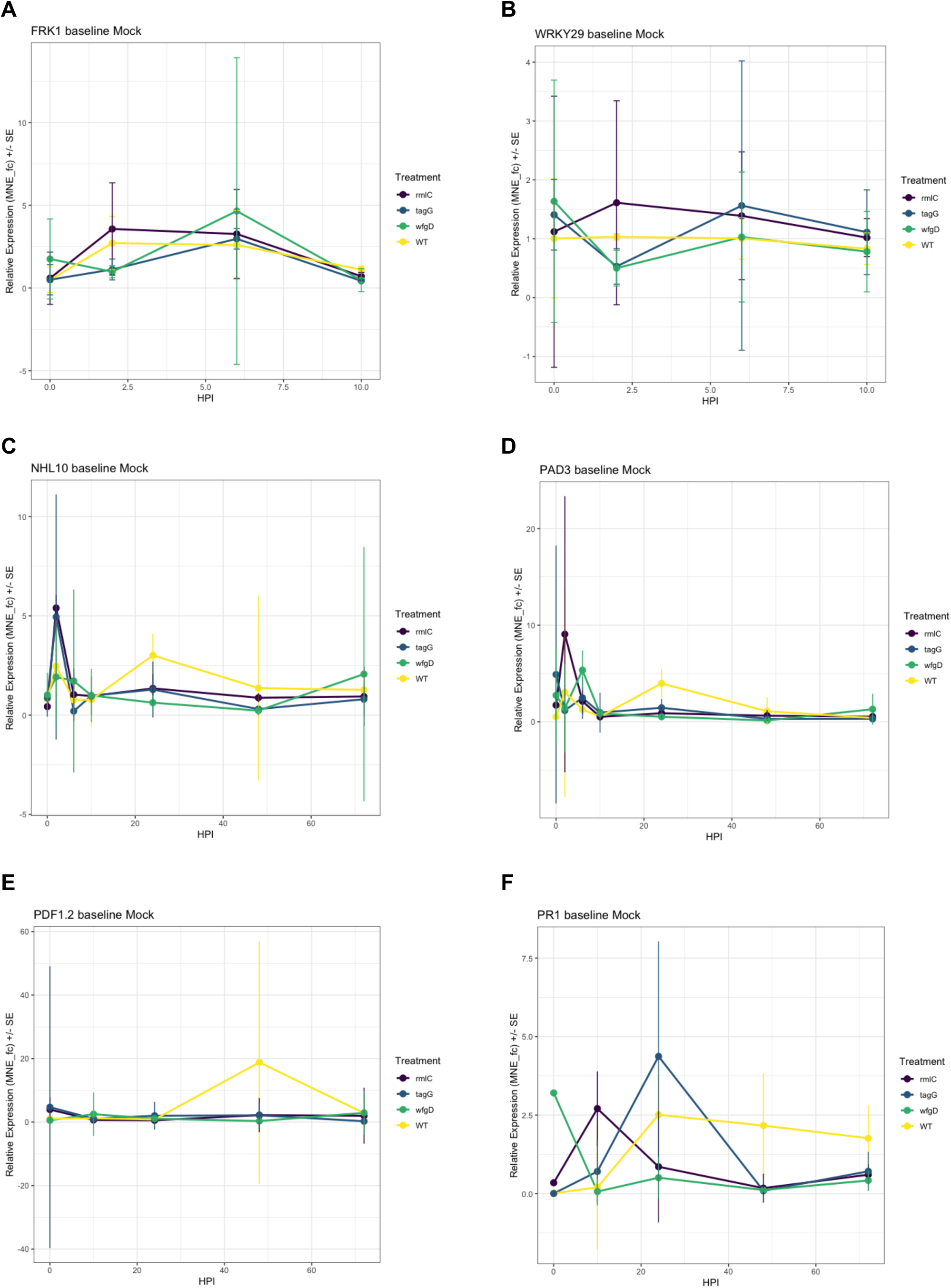
Immunity induction in *Arabidopsis thaliana* Col-0 following infection with p25.C2 (WT) and O-antigen mutants. Plants were flood-inoculated, and expression of immune marker genes *FRK1* (A), *WRKY29* (B), *NHL10* (C), *PAD3* (D), *PDF1.2* (E), and *PR1* (F) was assessed at 0, 2, 6, 10, 24, 48, and 72 hours post-infection (HPI). Expression was normalized to the housekeeping gene *PP2A* and expressed as fold change (fc, relative to mock) of the mean normalized expression (MNE), accounting for PCR efficiency. Data represent three technical replicates per condition across three biological replicates. The absence of significant difference among strains at each time point was assessed for each target using one-way ANOVA or Kruskal–Wallis tests, as appropriate.

**Figure S9.**
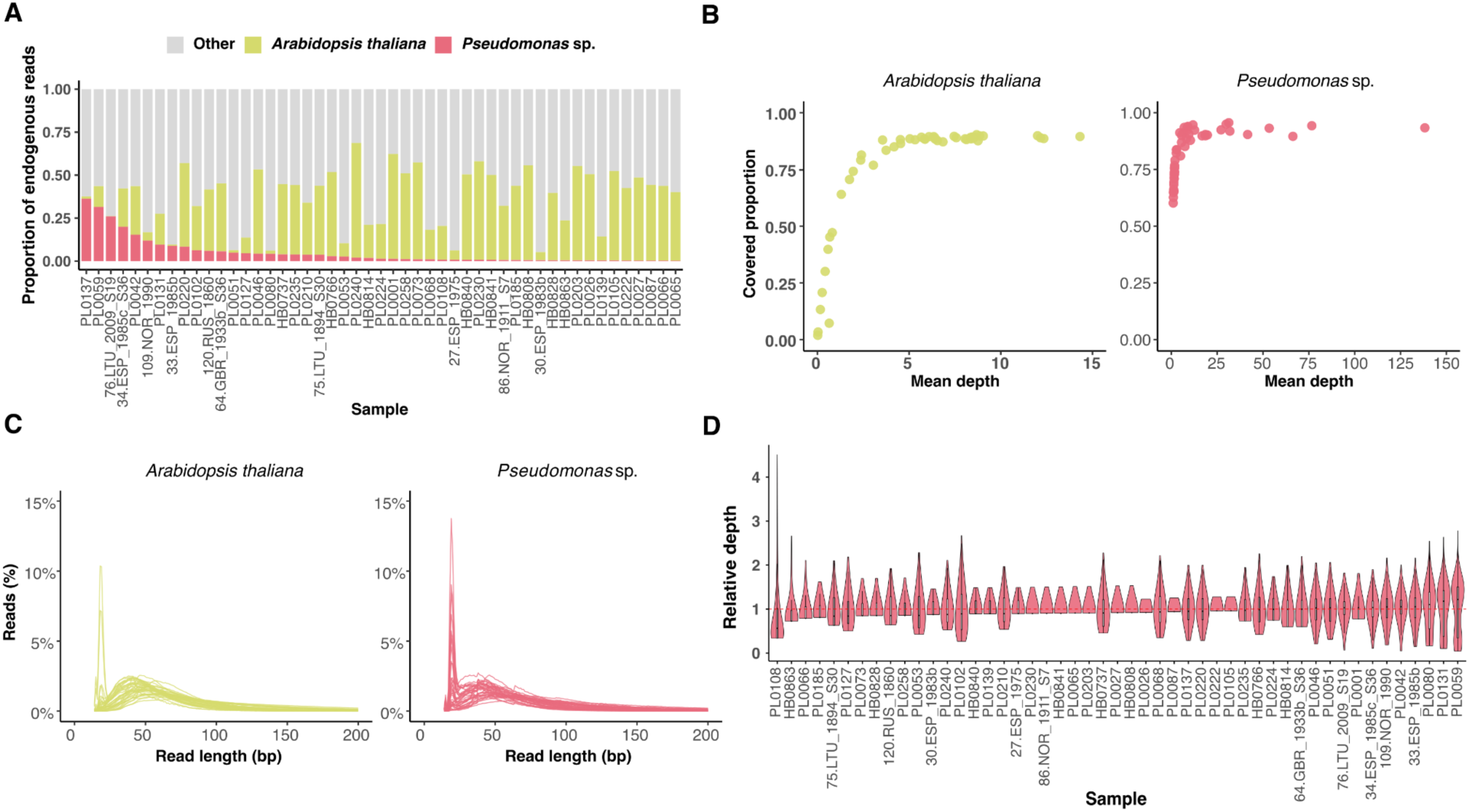
Endogenous DNA profiles reveal variation in historical pathogen representation. **A.** Endogenous read composition across all 49 historical metagenomic libraries. Stacked bars represent the proportion of reads assigned to *Arabidopsis thaliana* (olive-yellow), *Pseudomonas* sp. (red), and other taxa (grey) within each metagenome. **B.** A scatterplot showing the genome-wide mean read depth (x-axis) vs the breadth of genomic coverage (y-axis) for the *A. thaliana* (olive-yellow) and *Pseudomonas* sp. (red) genomes. Each point represents one historical sample. **C.** Read length distributions of *A. thaliana* (olive-yellow) and *Pseudomonas* sp. (red) reads from 39 historical samples. X-axis was limited to read lengths ≤ 200 bp to provide higher resolution. The remaining ten samples from Lopez et al. (2025) are shown separately in Fig. S10. **D.** Violin plots of *Pseudomonas* sp. whole-genome k-mer depth across historical isolates. The x-axis represents individual samples, and the y-axis shows k-mer depth normalized to each genome’s mean depth (mean = 1). The distributions reveal that most historical infections were dominated by a single *Pseudomonas* strain.

**Figure S10.**
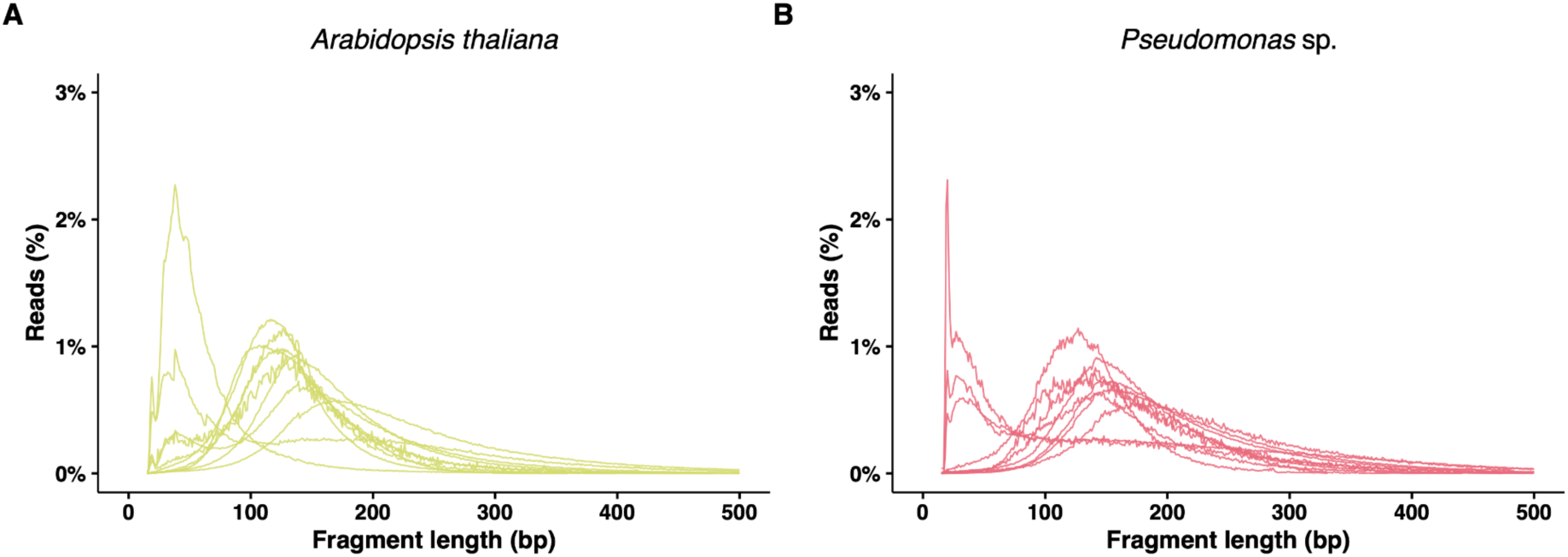
**A-B.** Read length distributions of reads mapped to *Arabidopsis thaliana* (olive-yellow) and *Pseudomonas* sp. (red) for the ten historical metagenomes from Lopez et al. (2025). X-axis was limited to read lengths ≤ 500 bp to provide higher resolution.

**Figure S11.**
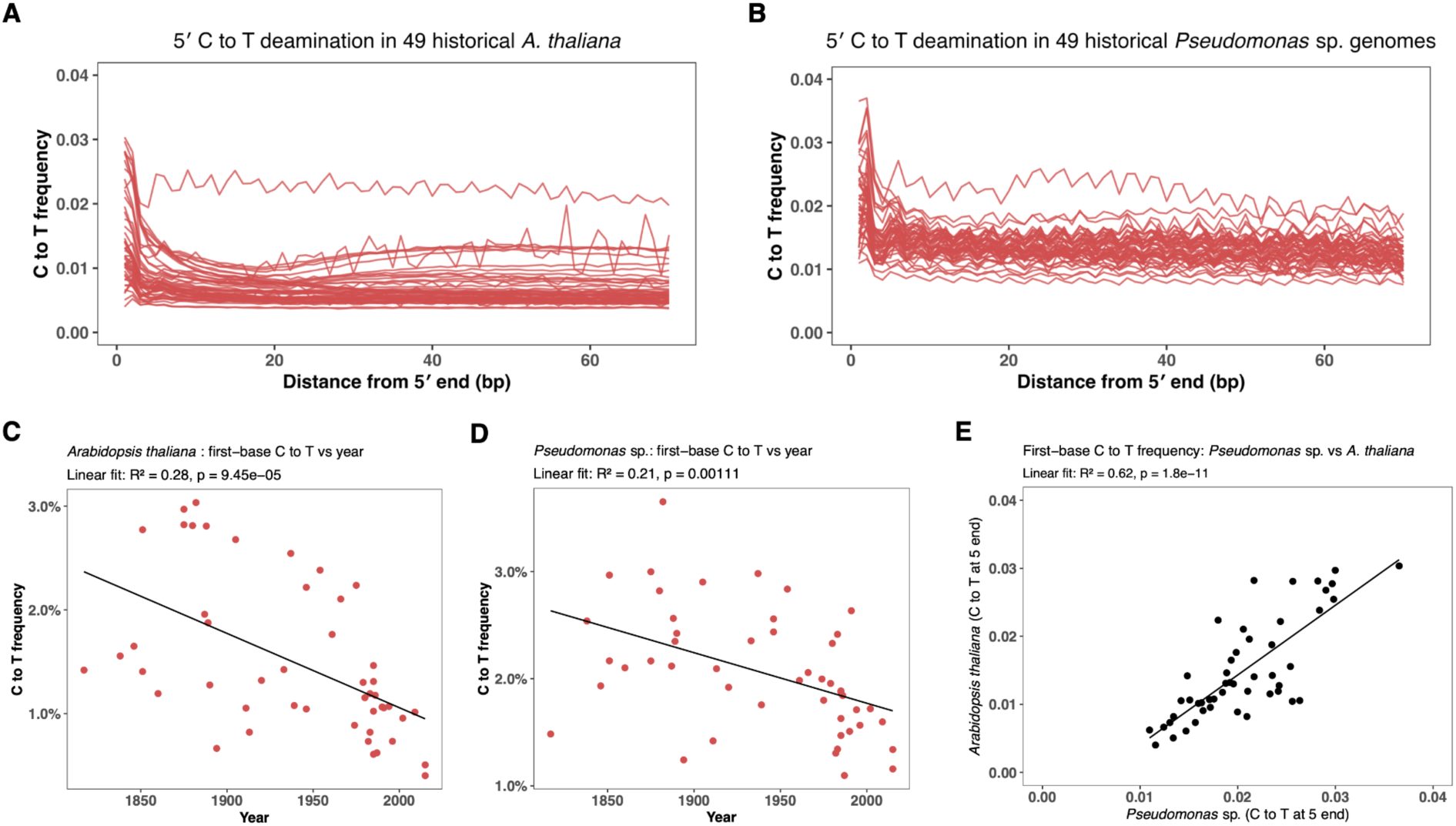
DNA damage patterns in host and pathogen reads support the authenticity of historical metagenomes. Accumulated C-to-T frequencies at the 5′ ends of sequencing reads for 49 *Arabidopsis thaliana* (**A.**) and 49 *Pseudomonas* sp. genomes (**B.**). The y-axis represents C-to-T substitution frequency, and the x-axis shows positions in base pairs from the 5′ end of each read. Both datasets exhibit a typical C-to-T substitutions pattern decaying from the 5′ ends, consistent with historical DNA damage. **C-D.** Scatter plots showing the relationship between the proportion of C-to-T substitutions at the first base (y-axis) and the collection year of each sample (x-axis) for *A. thaliana* (**C.**) and *Pseudomonas* sp. (**D.**). Linear regressions reveal significant negative correlations (*A. thaliana*: R²=0.28, p value=9.45×10⁻⁵; *Pseudomonas* sp.: R²=0.21, p value=1.11×10⁻^3^), indicating that older samples exhibit higher levels of cytosine deamination. Sample PL0087 was excluded due to missing year information. **E.** Correlation between C-to-T substitution frequencies at the 5′ first bases of reads mapped to *Pseudomonas* sp. vs *A. thaliana*. A significant positive correlation confirms that DNA damage signatures are acquired at similar rates between hosts and pathogens.

**Figure S12.**
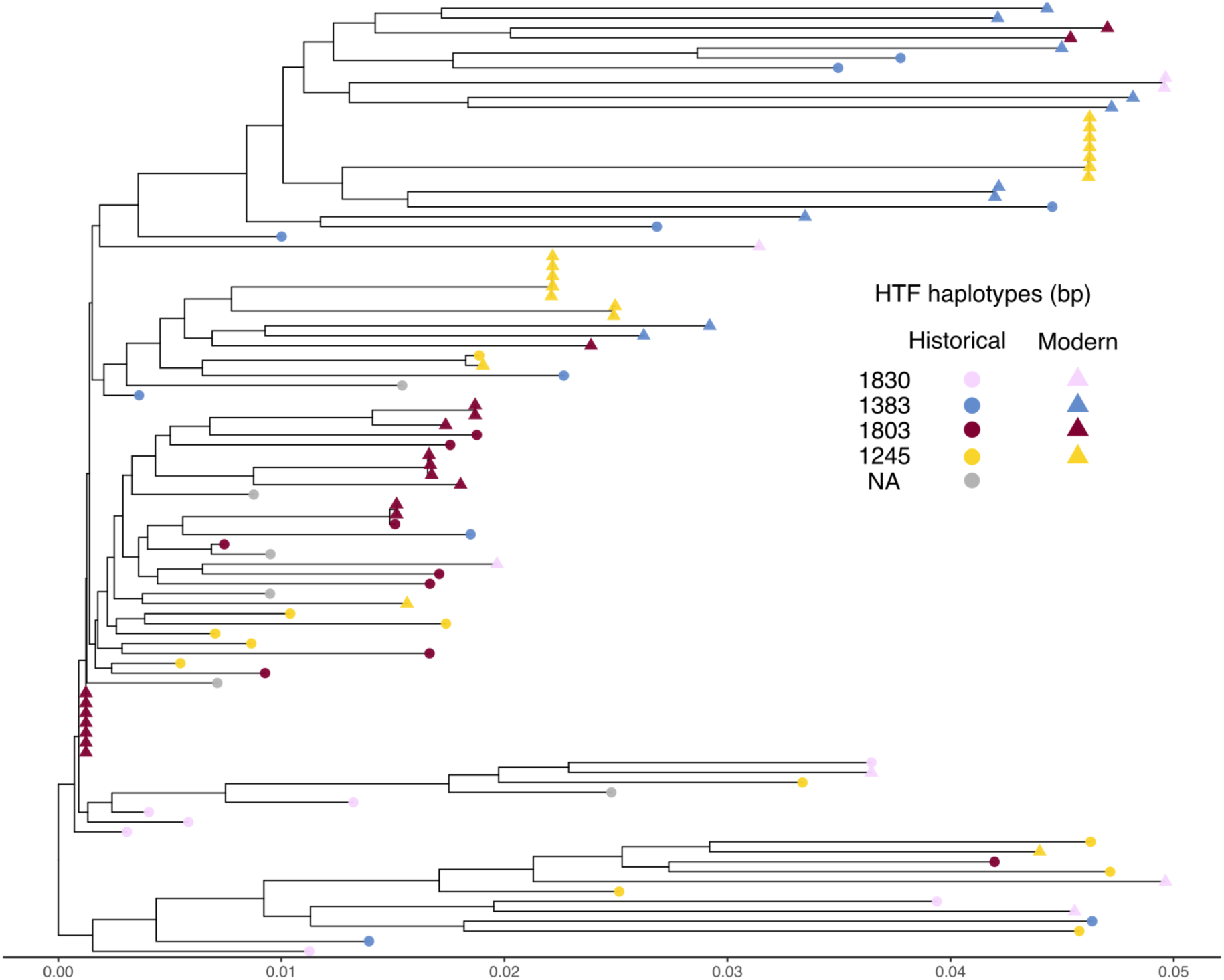
Historical and modern ATUE5 reveal phylogenetic continuity over centuries. Midpoint-rooted maximum-likelihood tree of 43 historical (circles) and 53 modern (triangles) *Pseudomonas viridiflava* ATUE5 genomes. Tip colours indicate the four dominant *HTF* length variants: 1830 bp (pink), 1383 bp (blue), 1803 bp (red) and 1245 bp (yellow). Historical *HTF* variants are broadly distributed across all major branches of the ATUE5 phylogeny, indicating long-term temporal continuity within the lineage.

**Figure S13.**
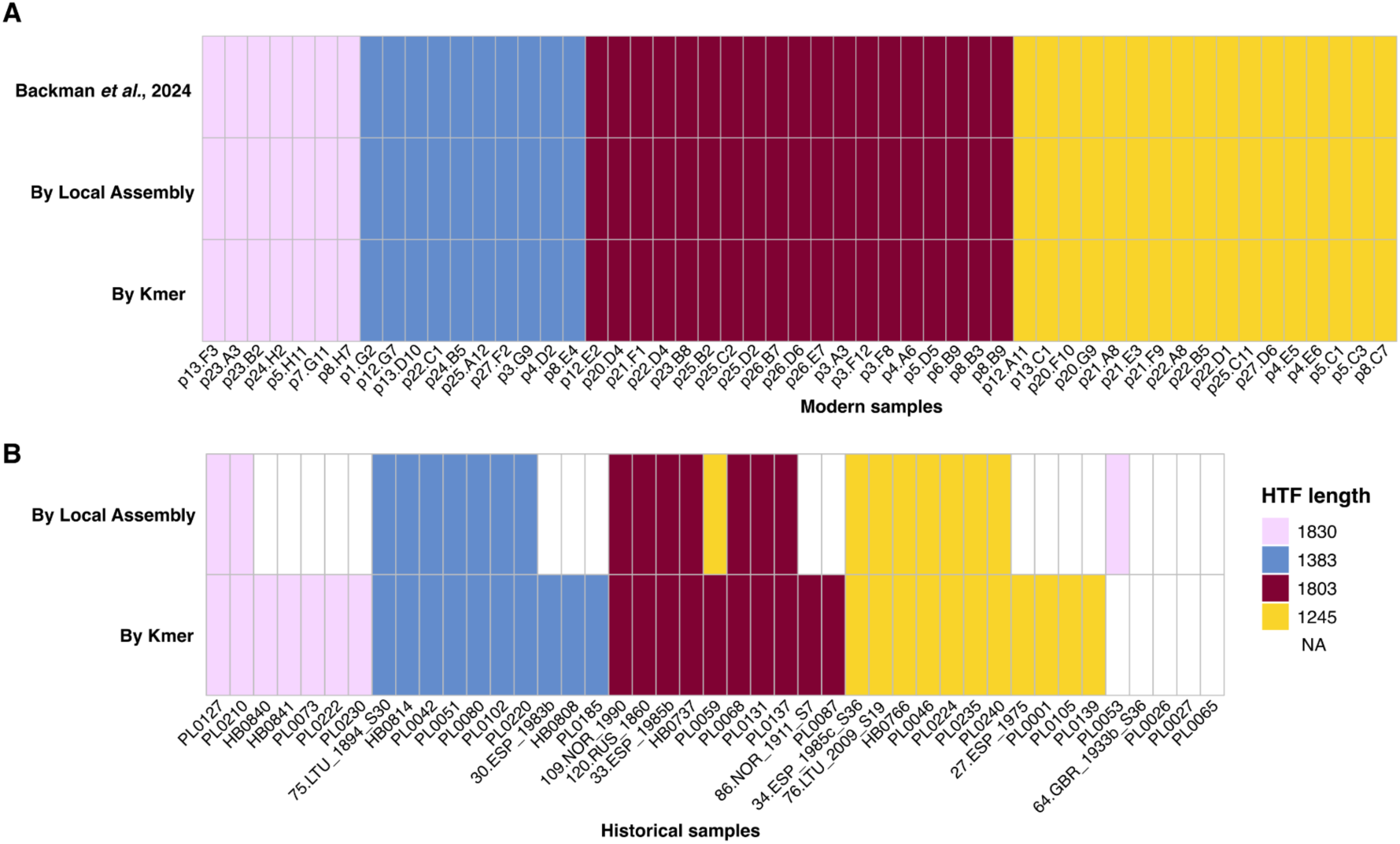
High concordance between k-mer and local assembly methods validates HTF haplotype reconstructions in historical genomes. **A.** All 53 modern genomes yielded fully concordant haplotype calls across both methods, consistent with previous assignments (Backman *et al*., 2024), which validated the robustness of our approach. **B.** Of 43 historical genomes, 25 haplotypes were assigned by local assembly, and 38 by the k-mer method. Concordance among haplotypes assigned by both methods was 96% (23 of 24), with one sample (PL0059) showing conflicting haplotypes (1245 bp vs. 1803 bp), likely due to coinfection (Fig. S17). We used the 23 concordant and 15 additional k-mer–derived haplotypes for downstream analyses.

**Figure S14.**
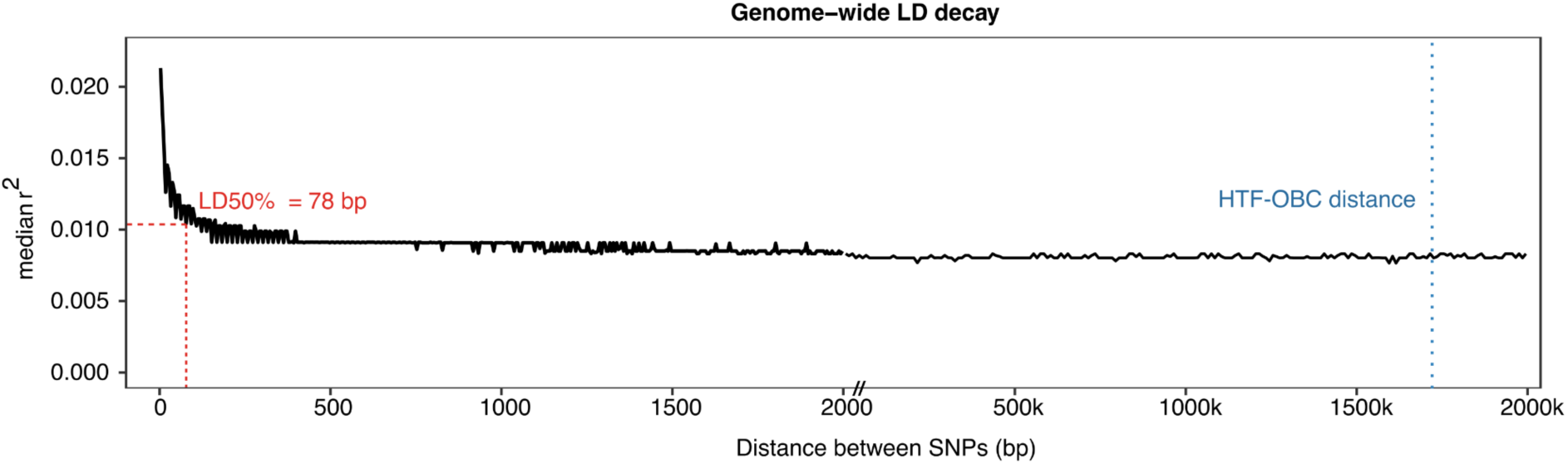
Genome-wide LD decay in *Pseudomonas viridiflava* ATUE5. Pairwise linkage disequilibrium (LD; median r²) between biallelic SNPs was calculated across 53 modern ATUE5 genomes and plotted against SNP distance. LD was estimated for SNP pairs spanning 0–2 kb using all SNPs, and for 2–2000 kb using a random 2% subset to reduce computation time. LD decays to 50% of its maximum value at 78 bp (LD50; red dashed lines) and reaches background levels by < 2 kb. The genetic distance between the HTF and OBC loci (1.7 Mb; blue dashed line) is shown for reference.

**Figure S15.**
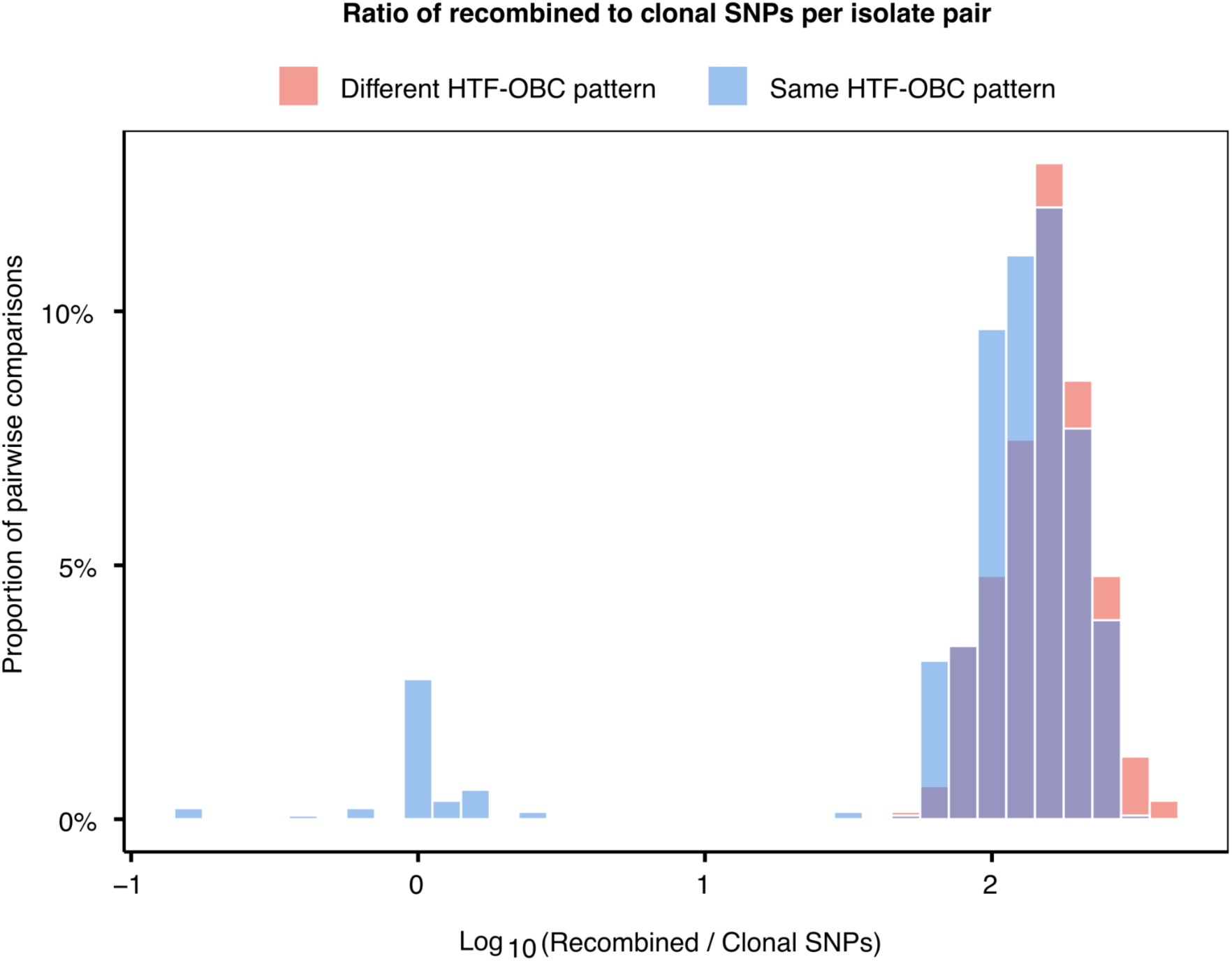
Ratio of recombined to clonal SNPs in pairwise comparisons of *Pseudomonas viridiflava* ATUE5. Distribution of log₁₀(Recombined / Clonal SNPs) across 1,378 pairwise comparisons among 53 modern ATUE5 genomes. Bars indicate the proportion of pairs sharing the same (blue) or different (red) HTF–OBC pattern. 1,318 pairs (95.6%) exhibited log₁₀(Recombined / Clonal SNPs) > 1, indicating that recombination generated more than tenfold the number of clonal SNPs. On average, recombination accounted for 96% of SNPs per isolate.

**Figure S16.**
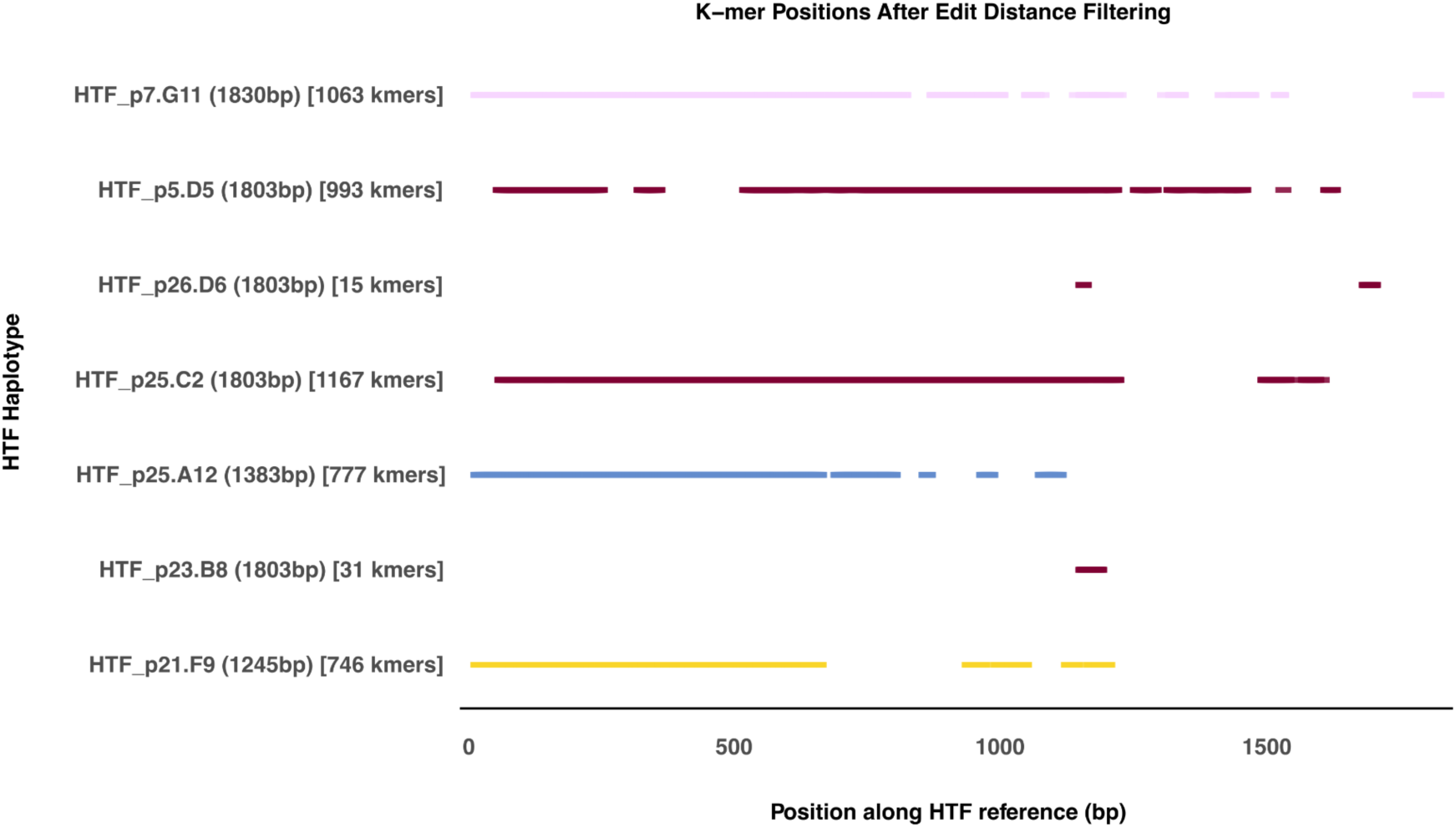
Distribution of diagnostic k-mers across HTF reference sequences. Each tick represents the position of a k-mer along the *HTF* nucleotide sequences (x-axis) after applying an edit-distance filter (>=2). Y-axis labels show *HTF* haplotype names, and the total number of retained k-mers per haplotype.

**Figure S17.**
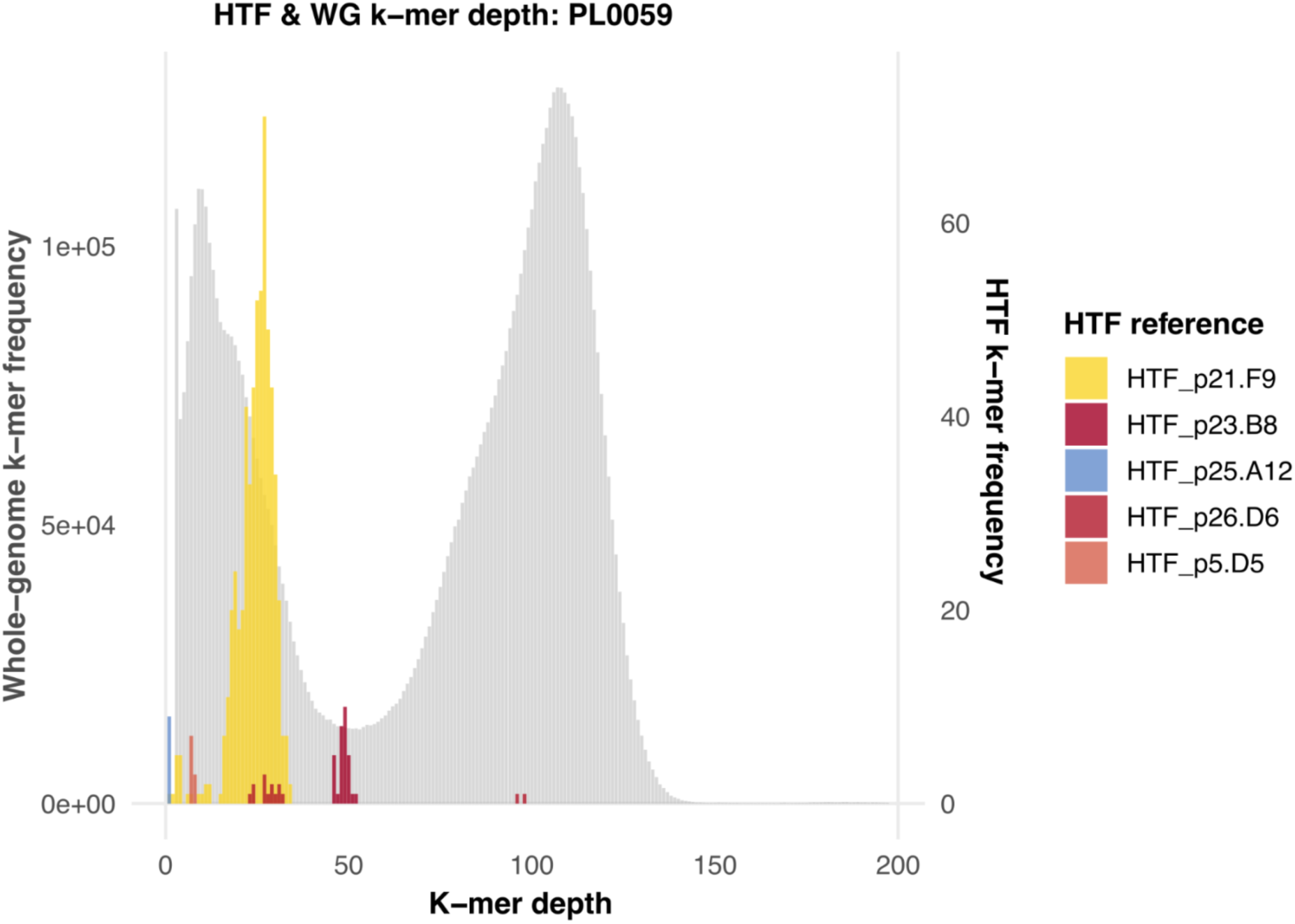
Bimodal k-mer depth distribution of PL0059 suggests coinfection. The histograms represent k-mer depth distributions. The grey background shows the whole-genome distribution, and the colored histograms represent the distribution for each of the diagnostic *HTF* haplotype k-mers. The bimodal pattern in the whole genome *k*-mer distribution indicates potential coinfection in PL0059. *K*-mer distributions for other isolates are available on GitHub.

**Table S1:** RB-TnSeq significant resistance trade-off genes.

| Barcode ID | Log<sub>2</sub><br>(fold change)<br>in Col-0 | p-value<br>Col-0 | Log<br>(fold change)<br>after tailocin<br>treatment | p-value<br>tailocin<br>assay | Log<sub>2</sub><br>(fold change)<br>in Eyach1.5-2 | p-value<br>Eyach<br>1.5-2 | Name gene or<br>product | Significance<br>in both<br>ecotypes |
| --- | --- | --- | --- | --- | --- | --- | --- | --- |
| BJEIHDPM_02277 | -2.284 | 1.09E-06 | 3.560 | 1.38E-15 | -1.570 | 7.8E-04 | *chpB* | Both |
| BJEIHDPM_03520 | -0.790 | 0.023 | 0.633 | 0.035 | 0.140 | 0.685 | *yciF* | Col-0 only |
| BJEIHDPM_02879 | -1.281 | 0.023 | 2.396 | 1.69E-09 | -0.365 | 0.512 | BolA-like protein | Col-0 only |
| BJEIHDPM_04292 | -0.092 | 0.759 | 3.436 | 7.25E-16 | -0.710 | 0.025 | Hypothetical protein | Eyach 1.5-2 only |
| BJEIHDPM_01314 | -1.152 | 0.024 | 5.499 | 5.78E-33 | -1.479 | 0.005 | Glycosyltransferase | Both |
| BJEIHDPM_01216 | 0.083 | 0.826 | 1.199 | 1.46E-06 | -0.907 | 0.021 | *ureC_2* | Eyach 1.5-2 only |
| BJEIHDPM_00728 | -0.775 | 0.018 | 0.704 | 0.006 | 0.155 | 0.638 | *ugpC_1* | Col-0 only |
| BJEIHDPM_03096 | -0.709 | 0.022 | 0.530 | 0.030 | 0.063 | 0.838 | *cmoB* | Col-0 only |
| BJEIHDPM_03578 | -0.807 | 0.006 | 1.719 | 8.36E-15 | -0.900 | 0.003 | *flgE* | Both |
| BJEIHDPM_00926 | -0.964 | 0.035 | 5.653 | 3.2E-29 | -0.731 | 0.117 | *rmlC_1* | Col-0 only |
| BJEIHDPM_00922 | -0.379 | 0.026 | 6.969 | 3.02E-41 | -0.355 | 0.040 | *rfbB* | Both |
| BJEIHDPM_00925 | -1.200 | 2E-09 | 4.909 | 2.38E-29 | -1.363 | 2.09E-11 | *wfgD* | Both |
| BJEIHDPM_00927 | -2.879 | 7.87E-13 | 4.696 | 8.7E-23 | -2.469 | 1.27E-09 | *tagG_2* | Both |
| BJEIHDPM_00929 | -0.747 | 7.4E-04 | 4.991 | 1.01E-34 | -1.107 | 8.12E-07 | *epsE_4* | Both |
| BJEIHDPM_00928 | -3.293 | 5.98E-14 | 4.377 | 8.78E-20 | -3.892 | 1.87E-16 | *tagH_2* | Both |
| BJEIHDPM_00924 | -1.212 | 8.23E-06 | 6.776 | 5.76E-40 | -1.092 | 7.84E-05 | *rmlA2_1* | Both |
| BJEIHDPM_00535 | -0.200 | 0.526 | 2.040 | 1.74E-17 | -0.909 | 0.005 | *dapL* | Eyach 1.5-2 only |
| BJEIHDPM_03394 | -1.276 | 1.2E-04 | 5.187 | 2.43E-28 | -1.436 | 2.27E-05 | *tagO* | Both |
| BJEIHDPM_03395 | -1.018 | 2.79E-07 | 5.024 | 3.51E-29 | -0.955 | 2E-06 | *pglF* | Both |
| BJEIHDPM_01313 | -1.324 | 1.3E-09 | 4.865 | 7.59E-30 | -0.934 | 2.32E-05 | O-antigen ligase domain-<br>containing protein | Both |
| BJEIHDPM_01315 | -2.185 | 1.55E-06 | 6.460 | 2.76E-33 | -2.926 | 1.52E-09 | Glycosyltransferase | Both |

**Table S2:**
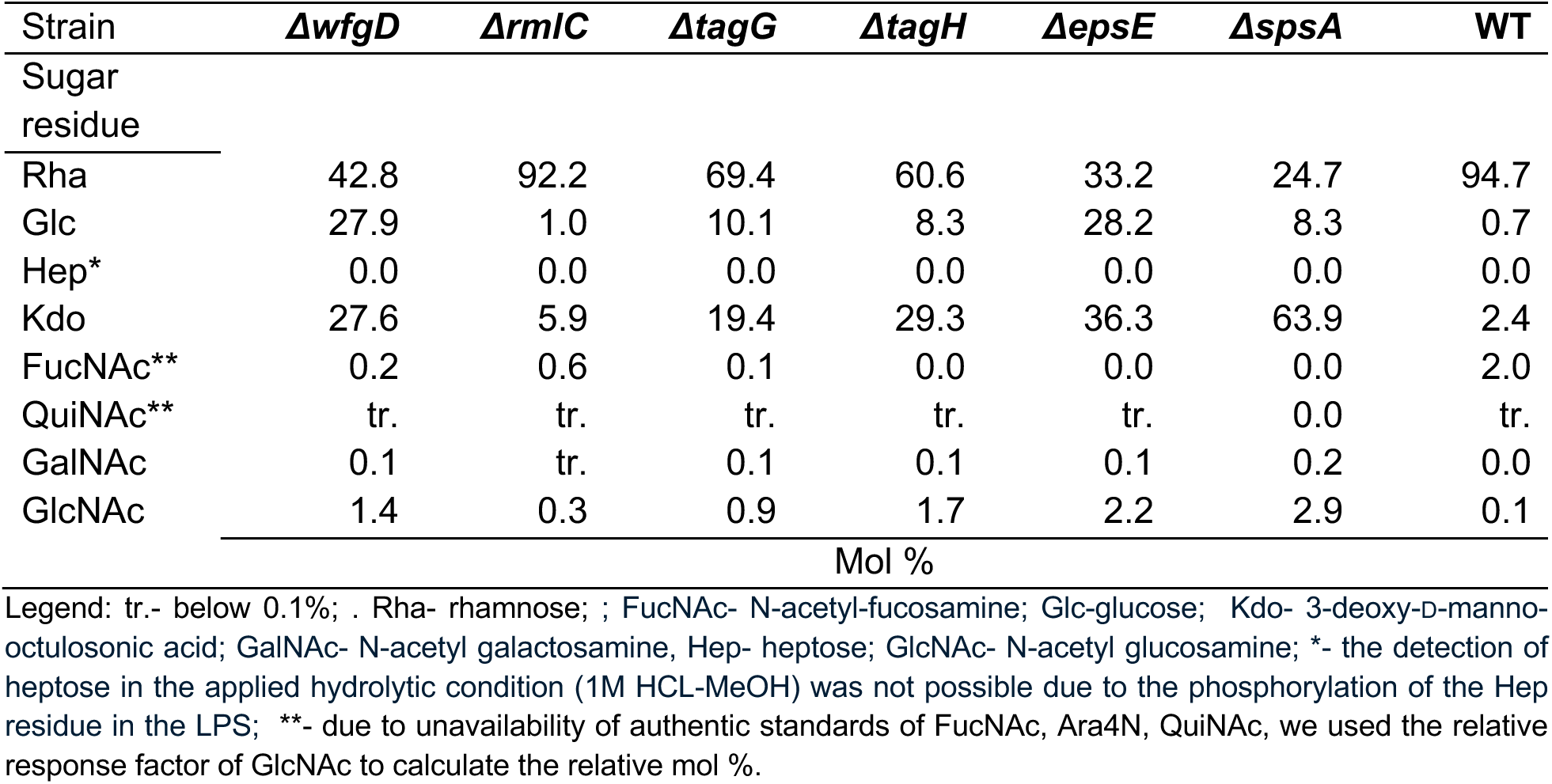
Comparative glycosyl composition analysis of p25.C2 (WT) LPS and the O-antigen mutants. The LPS was analyzed by the GC-MS after acidic methanolysis, and conversion of the monosaccharides to trimethylsilyl (TMS)-methyl glycosides.

| Strain | *ΔwfgD* | *ΔrmIC* | *ΔtagG* | *ΔtagH* | *ΔepsE* | *ΔspsA* | WT |
| --- | --- | --- | --- | --- | --- | --- | --- |
| Sugar residue |  |  |  |  |  |  |  |
| Rha | 42.8 | 92.2 | 69.4 | 60.6 | 33.2 | 24.7 | 94.7 |
| Glc | 27.9 | 1.0 | 10.1 | 8.3 | 28.2 | 8.3 | 0.7 |
| Hep* | 0.0 | 0.0 | 0.0 | 0.0 | 0.0 | 0.0 | 0.0 |
| Kdo | 27.6 | 5.9 | 19.4 | 29.3 | 36.3 | 63.9 | 2.4 |
| FucNAc** | 0.2 | 0.6 | 0.1 | 0.0 | 0.0 | 0.0 | 2.0 |
| QuiNAc** | tr. | tr. | tr. | tr. | tr. | 0.0 | tr. |
| GalNAc | 0.1 | tr. | 0.1 | 0.1 | 0.1 | 0.2 | 0.0 |
| GlcNAc | 1.4 | 0.3 | 0.9 | 1.7 | 2.2 | 2.9 | 0.1 |
| Mol % |  |  |  |  |  |  |  |

Legend: tr.- below 0.1%; . Rha- rhamnose; ; FucNAc- N-acetyl-fucosamine; Glc- glucose; Kdo- 3-deoxy-D-manno-octulosonic acid; GalNAc- N-acetyl galactosamine, Hep- heptose; GlcNAc- N-acetyl glucosamine; \*- the detection of heptose in the applied hydrolytic condition (1M HCL-MeOH) was not possible due to the phosphorylation of the Hep residue in the LPS; \*\*- due to unavailability of authentic standards of FucNAc, Ara4N, QuiNAc, we used the relative response factor of GlcNAc to calculate the relative mol %.

**Table S3:** Statistics associated with the growth curves of p25.C2 WT versus O-antigen mutants. Growth curve parameters *k*, *r* (growth rates), t_gen (generation time) and AUC (area under the curve) were extracted for each curve (Fig. S4). Statistical analysis was performed on the means of three independent biological replicates with 4 technical replicates each. Differences were assessed by one-way analysis of variance (ANOVA1) followed by Tukey’s post hoc test for each parameter (p-values: *** < 0.001, ** < 0.01, * < 0.05).

| PARAMETER | STRAIN | WT | $\Delta epsE$ | $\Delta rmlC$ | $\Delta spsA$ | $\Delta tagG$ | $\Delta tagH$ | $\Delta wfgD$ |
| --- | --- | --- | --- | --- | --- | --- | --- | --- |
| ***k*** | WT |  |  |  |  |  |  |  |
| | $\Delta epsE$ | ** | | | | | | |
| | $\Delta rmlC$ | NS | * | | | | | |
| | $\Delta spsA$ | ** | NS | NS | | | | |
| | $\Delta tagG$ | ** | NS | NS | NS | | | |
| | $\Delta tagH$ | NS | NS | NS | NS | NS | | |
| | $\Delta wfgD$ | ** | NS | * | NS | NS | NS | |
| ***r*** | WT |  |  |  |  |  |  |  |
| | $\Delta epsE$ | NS | | | | | | |
| | $\Delta rmlC$ | NS | NS | | | | | |
| | $\Delta spsA$ | * | NS | ** | | | | |
| | $\Delta tagG$ | NS | NS | NS | * | | | |
| | $\Delta tagH$ | NS | NS | ** | NS | * | | |
| | $\Delta wfgD$ | NS | NS | * | NS | NS | NS | |
| ***t*<sub>gen</sub>** | WT |  |  |  |  |  |  |  |
| | $\Delta epsE$ | NS | | | | | | |
| | $\Delta rmlC$ | NS | NS | | | | | |
| | $\Delta spsA$ | * | NS | * | | | | |
| | $\Delta tagG$ | NS | NS | NS | * | | | |
| | $\Delta tagH$ | NS | NS | * | NS | * | | |
| | $\Delta wfgD$ | NS | NS | NS | NS | NS | NS | |
| **AUC** | WT |  |  |  |  |  |  |  |
| | $\Delta epsE$ | *** | | | | | | |
| | $\Delta rmlC$ | NS | ** | | | | | |
| | $\Delta spsA$ | *** | NS | *** | | | | |
| | $\Delta tagG$ | ** | NS | * | NS | | | |
| | $\Delta tagH$ | *** | NS | ** | NS | NS | | |
| | $\Delta wfgD$ | *** | NS | ** | NS | NS | NS | |

**Table S4:** Hydrogen peroxide minimum inhibitory concentrations (MICs) for WT and O-antigen mutants. MICs were determined as the lowest concentration of hydrogen peroxide that inhibited growth after 24 h in liquid culture, based on OD_600_ measurements collected in triplicate using a Tecan plate reader. Values represent mean ± SD (mM). “MIC/WT (est.)” indicates the relative MIC compared to wild-type; p-values reflect pairwise t-tests between each mutant and WT.

| Strain | MIC (mM, mean ± SD) | MIC/WT (est.) | p-value |
| --- | --- | --- | --- |
| WT | 5.0 ± 0.6 | 1.00 | – |
| *ΔepsE* | 6.9 ± 2.7 | 1.26 | 0.48 |
| *ΔrmlC* | 5.3 ± 0.6 | 1.06 | 0.17 |
| *ΔspsA* | 6.4 ± 2.7 | 1.26 | 0.55 |
| *ΔtagG* | 4.3 ± 0.6 | 0.87 | 0.27 |
| *ΔtagH* | 5.2 ± 0.6 | 1.03 | 0.87 |
| *ΔwfgD* | 6.6 ± 4.3 | 1.18 | 0.72 |

**Table S5:**
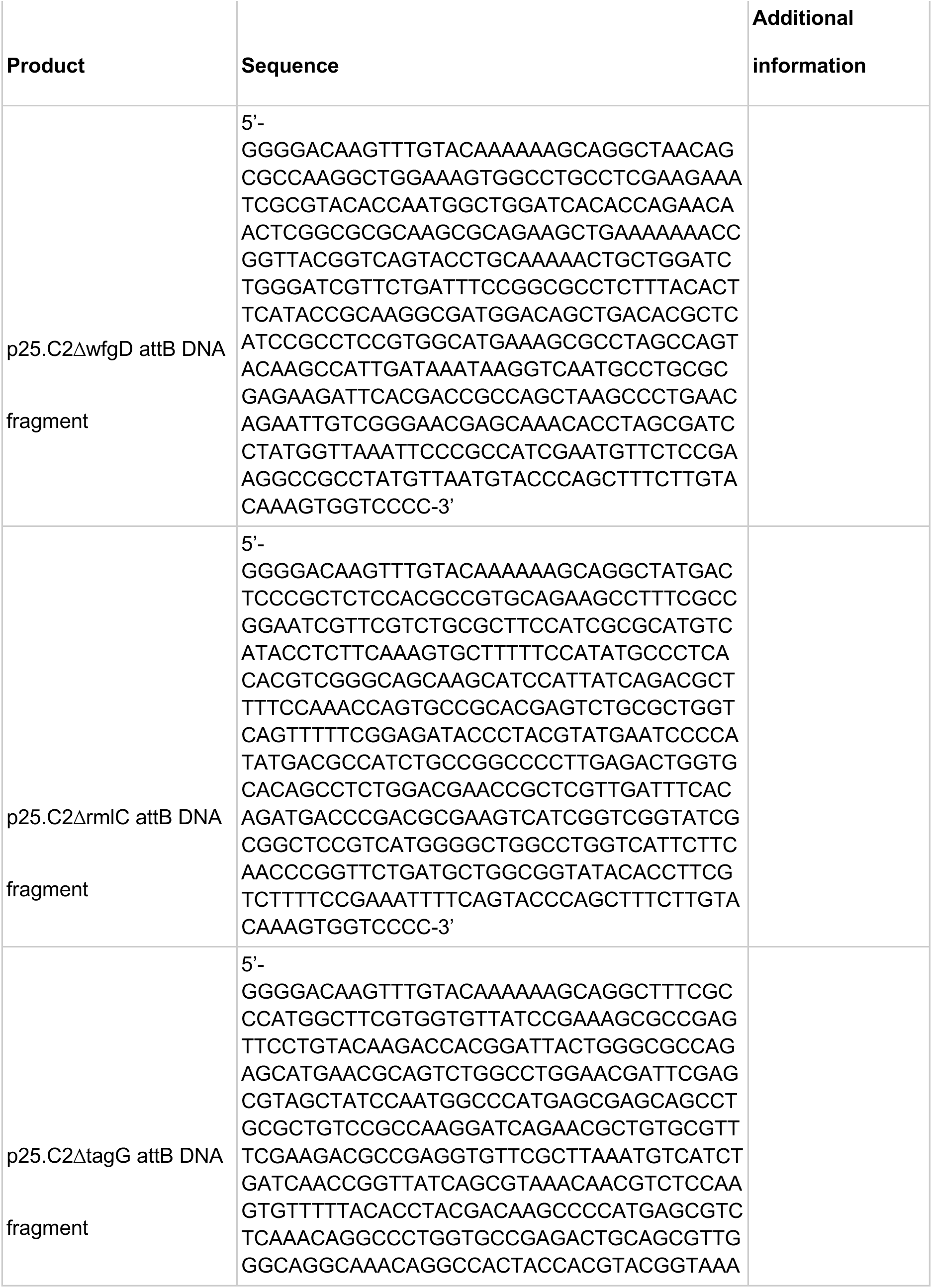
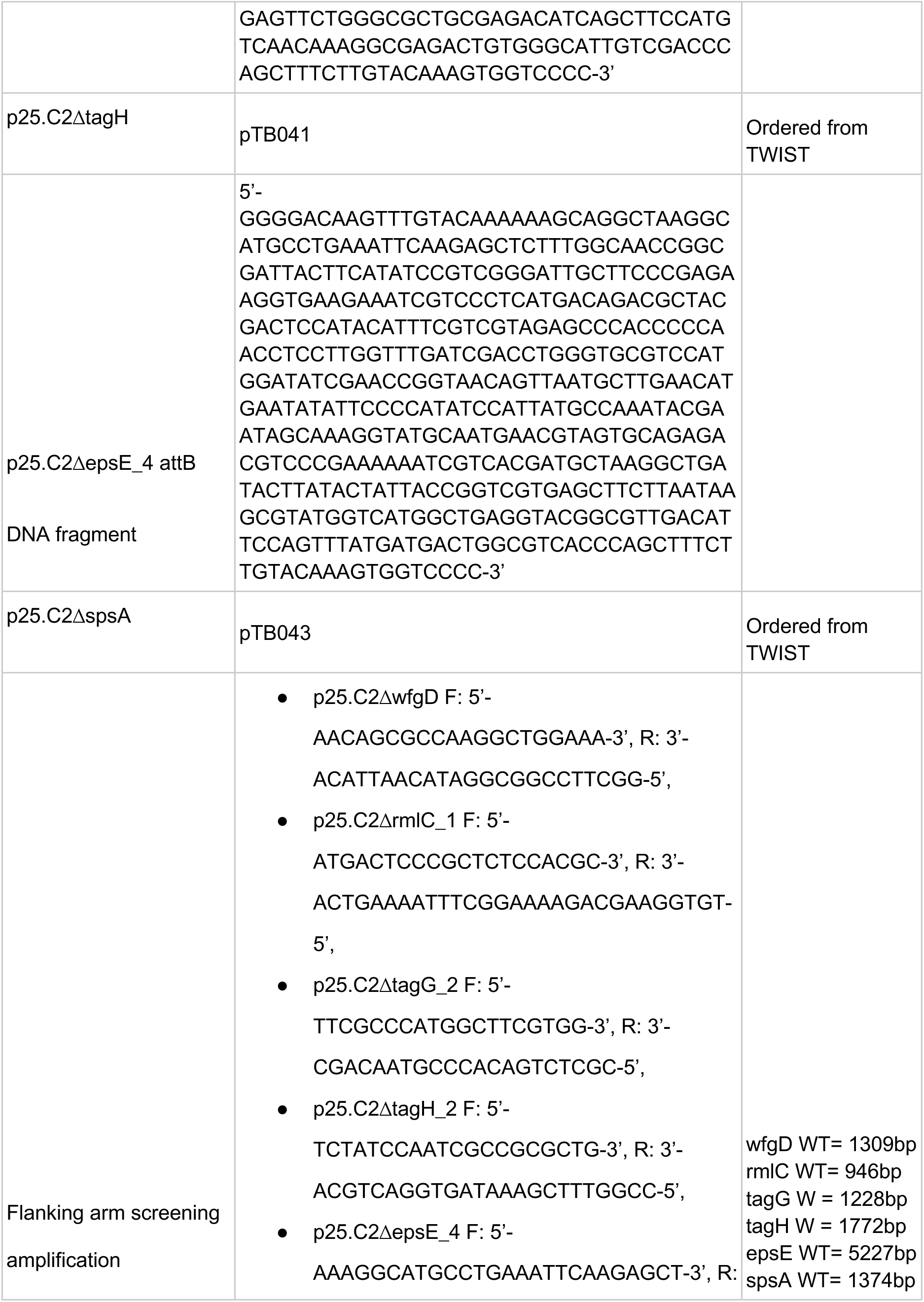
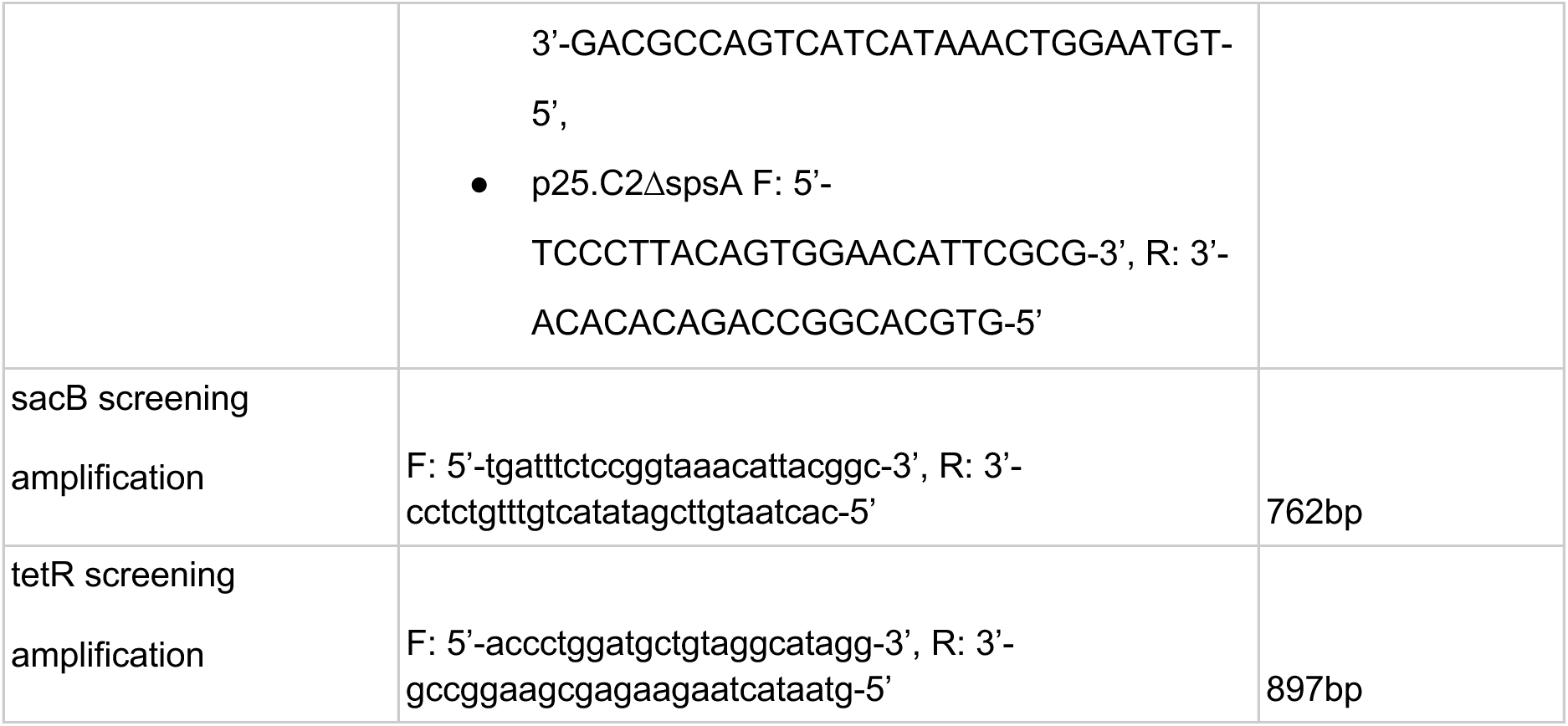
Plasmids used in this study for Gateway cloning.

**Table S6:**
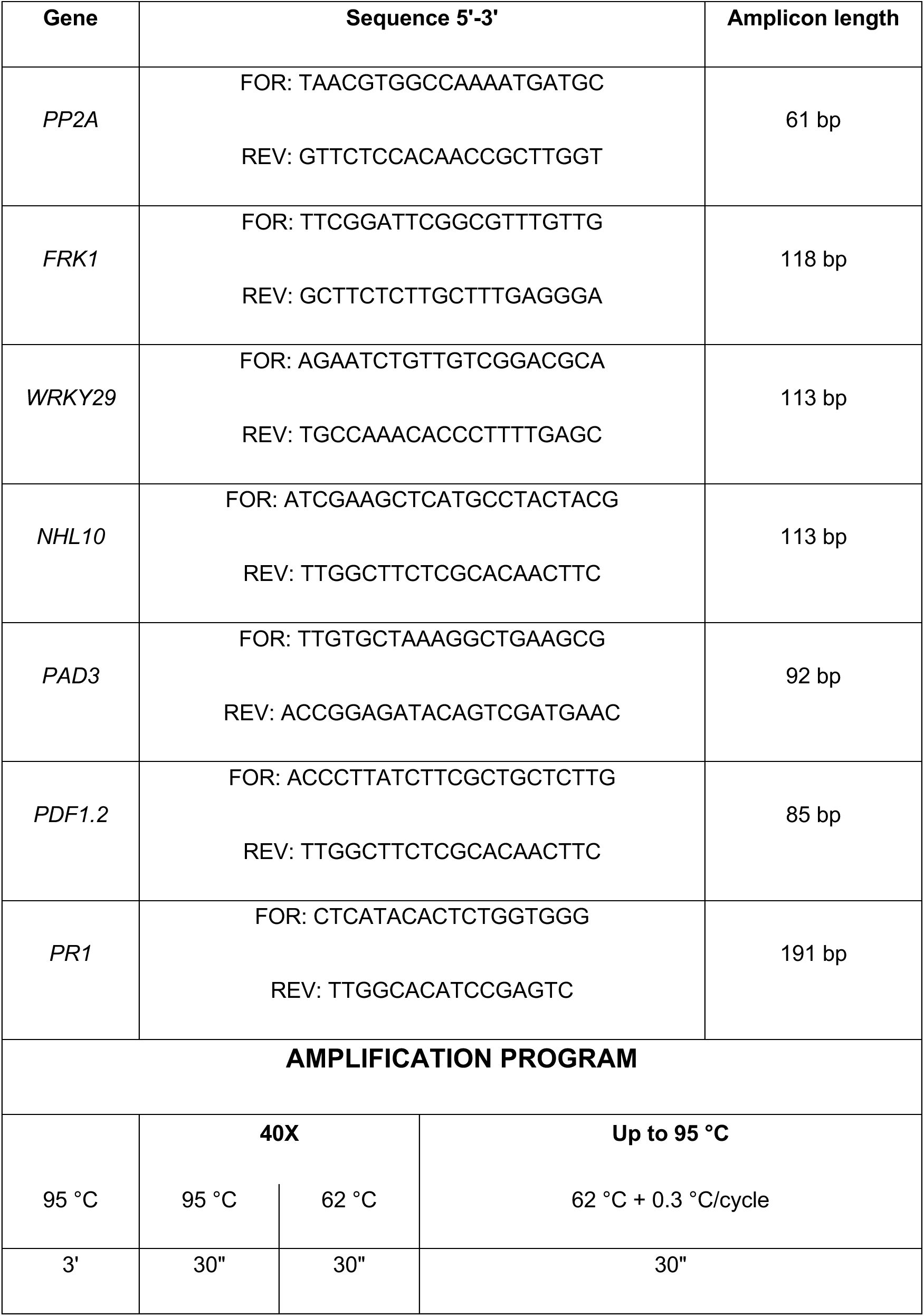
qPCR primers and amplification program used in this study.

**Table S7:** Historical sample metadata. **\* Herbarium: KBI** = Komarov Botanical Institute of RAS (Russia); **KEW** = Kew Botanical Garden (United Kingdom); **LND** = Lund University Botanical Museum (Sweden); **MGC** = Herbario de Málaga (Spain); **NHM** = The Natural History Museum (United Kingdom); **OHN** = Biological Museum Oskarshamn (Sweden); **RJB** = Real Jardin Botanico (Spain); **STG** = Staatliches Museum für Naturkunde Stuttgart (Germany); **TUE** = Herbarium Tubingense University of Tübingen (Germany); **UOS** = University of Oslo (Norway)

| SAMPLE | TYPE | YEAR | COUNTRY | At_proportion | At_covered<br>_proportion | At_averag<br>e_depth | Ps_proportion | Ps_covered<br>_proportion | Ps_avera<br>ge_depth | GROUP | HERBARIU<br>M* | SOURCE |
| --- | --- | --- | --- | --- | --- | --- | --- | --- | --- | --- | --- | --- |
| PL0001 | Plant tissue<br>herbarium | 1966 | Spain | 61.00% | 87.61% | 8.76 | 1.31% | 90.66% | 4.58 | ATUE5 | MGC | This study.<br>ENA:<br>PRJEB98841 |
| PL0026 | Plant tissue<br>herbarium | 2002 | Sweden | 50.00% | 89.86% | 12.01 | 0.50% | 72.29% | 1.58 | ATUE5 | OHN | This study.<br>ENA:<br>PRJEB98841 |
| PL0059 | Plant tissue<br>herbarium | 1985 | Spain | 12.10% | 87.14% | 6.85 | 31.47% | 92.87% | 137.41 | ATUE5 | RJB | This study.<br>ENA:<br>PRJEB98841 |
| PL0073 | Plant tissue<br>herbarium | 1991 | Spain | 56.30% | 88.14% | 8.13 | 1.05% | 83.24% | 2.83 | ATUE5 | RJB | This study.<br>ENA:<br>PRJEB98841 |
| PL0087 | Plant tissue<br>herbarium | NA | Spain | 44.00% | 88.46% | 8.37 | 0.33% | 62.29% | 1.13 | ATUE5 | RJB | This study.<br>ENA:<br>PRJEB98841 |
| PL0102 | Plant tissue<br>herbarium | 1888 | Germany | 25.60% | 89.59% | 7.46 | 6.34% | 95.09% | 31.13 | ATUE5 | TUE | This study.<br>ENA:<br>PRJEB98841 |
| PL0105 | Plant tissue<br>herbarium | 1920 | Sweden | 51.90% | 89.42% | 8.6 | 0.48% | 64.37% | 1.18 | ATUE5 | LND | This study.<br>ENA:<br>PRJEB98841 |
| PL0108 | Plant tissue<br>herbarium | 1946 | Sweden | 19.60% | 89.74% | 9.05 | 0.82% | 80.50% | 4.89 | nonATU<br>E5 | LND | This study.<br>ENA:<br>PRJEB98841 |
| PL0131 | Plant tissue<br>herbarium | 1980 | Spain | 18.00% | 87.70% | 6.55 | 9.59% | 92.67% | 53.07 | ATUE5 | RJB | This study.<br>ENA:<br>PRJEB98841 |
| PL0139 | Plant tissue<br>herbarium | 1996 | Sweden | 13.90% | 79.11% | 2.36 | 0.48% | 70.92% | 1.52 | ATUE5 | OHN | This study.<br>ENA:<br>PRJEB98841 |
| PL0185 | Plant tissue<br>herbarium | 1994 | Germany | 43.20% | 89.06% | 8.35 | 0.60% | 81.74% | 2.7 | ATUE5 | STG | This study.<br>ENA:<br>PRJEB98841 |
| PL0203 | Plant tissue<br>herbarium | 1880 | United<br>Kingdom | 54.80% | 88.49% | 8.06 | 0.50% | 72.39% | 1.68 | nonATU<br>E5 | NHM | This study.<br>ENA:<br>PRJEB98841 |
| PL0210 | Plant tissue<br>herbarium | 1838 | United<br>Kingdom | 30.30% | 84.96% | 4.18 | 3.70% | 87.36% | 10.36 | ATUE5 | NHM | This study.<br>ENA:<br>PRJEB98841 |
| PL0220 | Plant tissue<br>herbarium | 1946 | United<br>Kingdom | 48.60% | 88.47% | 5.84 | 8.37% | 89.32% | 18.44 | ATUE5 | NHM | This study.<br>ENA:<br>PRJEB98841 |
| PL0222 | Plant tissue<br>herbarium | 1887 | United<br>Kingdom | 42.10% | 89.12% | 8.55 | 0.36% | 64.92% | 1.23 | ATUE5 | NHM | This study.<br>ENA:<br>PRJEB98841 |
| PL0224 | Plant tissue<br>herbarium | 1889 | United<br>Kingdom | 20.20% | 89.01% | 6.4 | 1.36% | 93.00% | 7.01 | ATUE5 | NHM | This study.<br>ENA:<br>PRJEB98841 |
| PL0230 | Plant tissue<br>herbarium | 1851 | United<br>Kingdom | 57.30% | 90.32% | 8.68 | 0.73% | 78.60% | 2.01 | ATUE5 | NHM | This study.<br>ENA:<br>PRJEB98841 |
| PL0235 | Plant tissue<br>herbarium | 1905 | United<br>Kingdom | 40.30% | 86.39% | 4.54 | 3.90% | 91.81% | 8.56 | ATUE5 | NHM | This study.<br>ENA:<br>PRJEB98841 |
| PL0240 | Plant tissue<br>herbarium | 1875 | United<br>Kingdom | 66.70% | 88.97% | 7.57 | 2.02% | 84.63% | 7.38 | ATUE5 | NHM | This study.<br>ENA:<br>PRJEB98841 |
| PL0258 | Plant tissue<br>herbarium | 1913 | United<br>Kingdom | 49.90% | 88.39% | 6.42 | 1.15% | 83.22% | 2.75 | nonATUE5 | NHM | This study.<br>ENA:<br>PRJEB98841 |
| PL0027 | Plant tissue<br>herbarium | 1939 | Sweden | 48.30% | 89.78% | 8.85 | 0.33% | 65.78% | 1.27 | ATUE5 | OHN | This study.<br>ENA:<br>PRJEB98841 |
| PL0042 | Plant tissue<br>herbarium | 1982 | Spain | 28.20% | 88.62% | 8.49 | 15.37% | 93.77% | 76.14 | ATUE5 | RJB | This study.<br>ENA:<br>PRJEB98841 |
| PL0046 | Plant tissue<br>herbarium | 1974 | Spain | 49.00% | 88.93% | 12.17 | 4.31% | 89.76% | 19.52 | ATUE5 | RJB | This study.<br>ENA:<br>PRJEB98841 |
| PL0051 | Plant tissue<br>herbarium | 1983 | Spain | 1.50% | 20.73% | 0.26 | 4.89% | 89.92% | 18.44 | ATUE5 | RJB | This study.<br>ENA:<br>PRJEB98841 |
| PL0053 | Plant tissue<br>herbarium | 1987 | Spain | 7.70% | 64.01% | 1.3 | 2.68% | 94.16% | 11.72 | ATUE5 | RJB | This study.<br>ENA:<br>PRJEB98841 |
| PL0065 | Plant tissue<br>herbarium | 2015 | Spain | 39.90% | 88.61% | 12.35 | 0.21% | 68.88% | 1.35 | ATUE5 | RJB | This study.<br>ENA:<br>PRJEB98841 |
| PL0066 | Plant tissue<br>herbarium | 2015 | Spain | 43.40% | 89.49% | 14.32 | 0.32% | 70.50% | 1.61 | nonATUE5 | RJB | This study.<br>ENA:<br>PRJEB98841 |
| PL0068 | Plant tissue<br>herbarium | 1986 | Spain | 17.20% | 89.28% | 8.43 | 1.04% | 91.81% | 12.58 | ATUE5 | RJB | This study.<br>ENA:<br>PRJEB98841 |
| PL0080 | Plant tissue<br>herbarium | 1985 | Spain | 1.90% | 39.76% | 0.58 | 4.23% | 94.48% | 29.67 | ATUE5 | RJB | This study.<br>ENA:<br>PRJEB98841 |
| PL0127 | Plant tissue<br>herbarium | 1979 | Spain | 9.10% | 45.22% | 0.68 | 4.61% | 93.15% | 8.63 | ATUE5 | RJB | This study.<br>ENA:<br>PRJEB98841 |
| PL0137 | Plant tissue<br>herbarium | 1961 | Sweden | 1.30% | 1.89% | 0.02 | 36.32% | 89.29% | 16.67 | ATUE5 | OHN | This study.<br>ENA:<br>PRJEB98841 |
| 27.ESP_1<br>975 | Plant tissue<br>herbarium | 1975 | Spain | 5.50% | 7.22% | 0.64 | 0.76% | 75.00% | 1.71 | ATUE5 | RJB | (Lopez et al.,<br>2025) |
| 30.ESP_1<br>983b | Plant tissue<br>herbarium | 1983 | Spain | 4.70% | 47.14% | 0.81 | 0.54% | 75.39% | 1.74 | ATUE5 | RJB | (Lopez et al.,<br>2025) |
| 33.ESP_1<br>985b | Plant tissue<br>herbarium | 1985 | Spain | 0.80% | 13.26% | 0.16 | 8.86% | 91.39% | 31.62 | ATUE5 | RJB | (Lopez et al.,<br>2025) |
| 34.ESP_1<br>985c_S36 | Plant tissue<br>herbarium | 1985 | Spain | 22.30% | 81.36% | 2.4 | 19.98% | 89.98% | 41.23 | ATUE5 | RJB | (Lopez et al.,<br>2025) |
| 64.GBR_1<br>933b_S36 | Plant tissue<br>herbarium | 1933 | UK | 39.50% | 74.18% | 1.95 | 5.71% | 86.42% | 5.51 | ATUE5 | KEW | (Lopez et al.,<br>2025) |
| 75.LTU_1<br>894_S30 | Plant tissue<br>herbarium | 1894 | Lithuania | 40.20% | 87.96% | 3.56 | 3.67% | 90.64% | 6.43 | ATUE5 | KBI | (Lopez et al.,<br>2025) |
| 76.LTU_2<br>009_S19 | Plant tissue<br>herbarium | 2009 | Lithuania | 0.20% | 3.36% | 0.04 | 26.02% | 89.20% | 65.73 | ATUE5 | RJB | (Lopez et al.,<br>2025) |
| 86.NOR_1<br>911_S7 | Plant tissue<br>herbarium | 1911 | Norway | 31.60% | 83.44% | 3.75 | 0.60% | 67.85% | 1.35 | ATUE5 | UOS | (Lopez et al.,<br>2025) |
| 109.NOR_<br>1990 | Plant tissue<br>herbarium | 1990 | Norway | 4.80% | 30.04% | 0.42 | 11.96% | 91.97% | 26.84 | ATUE5 | UOS | (Lopez et al.,<br>2025) |
| 120.RUS_<br>1860 | Plant tissue<br>herbarium | 1860 | Russia | 35.70% | 70.54% | 1.75 | 5.96% | 89.46% | 5.59 | ATUE5 | KBI | (Lopez et al.,<br>2025) |
| HB0737 | Plant tissue<br>herbarium | 1937 | Germany | 40.80% | 88.15% | 4.52 | 3.95% | 90.08% | 9.5 | ATUE5 | TUE | (Lang et al.,<br>2024) |
| HB0766 | Plant tissue<br>herbarium | 1846 | Germany | 48.90% | 88.31% | 5.87 | 2.85% | 88.03% | 8.16 | ATUE5 | STG | (Lang et al.,<br>2024) |
| HB0808 | Plant tissue<br>herbarium | 1817 | Germany | 55.10% | 88.06% | 5.33 | 0.56% | 59.81% | 1.06 | ATUE5 | STG | (Lang et al.,<br>2024) |
| HB0814 | Plant tissue<br>herbarium | 1882 | Germany | 19.20% | 88.51% | 5.05 | 1.82% | 93.44% | 8.98 | ATUE5 | STG | (Lang et al.,<br>2024) |
| HB0828 | Plant tissue<br>herbarium | 1890 | Germany | 39.10% | 89.47% | 6.34 | 0.52% | 73.53% | 1.74 | nonATU<br>E5 | STG | (Lang et al.,<br>2024) |
| HB0840 | Plant tissue<br>herbarium | 1851 | Germany | 49.70% | 89.45% | 6.09 | 0.73% | 76.28% | 1.78 | ATUE5 | STG | (Lang et al.,<br>2024) |
| HB0841 | Plant tissue<br>herbarium | 1875 | Germany | 49.40% | 89.21% | 5.68 | 0.71% | 73.19% | 1.59 | ATUE5 | STG | (Lang et al.,<br>2024) |
| HB0863 | Plant tissue<br>herbarium | 1954 | Somalia | 23.20% | 76.88% | 3.05 | 0.51% | 65.01% | 1.49 | nonATU<br>E5 | STG | (Lang et al.,<br>2024) |

**Table S8:**
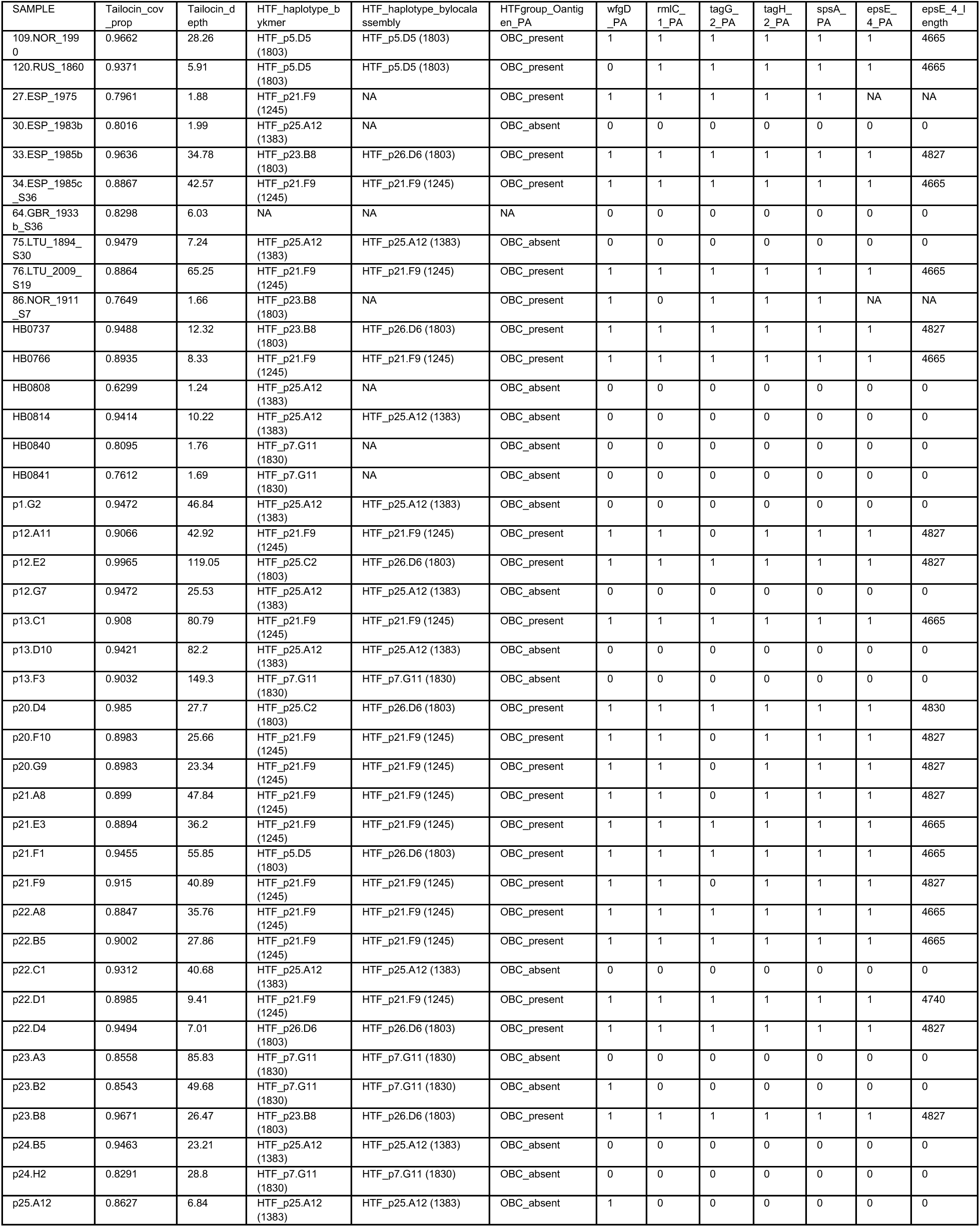
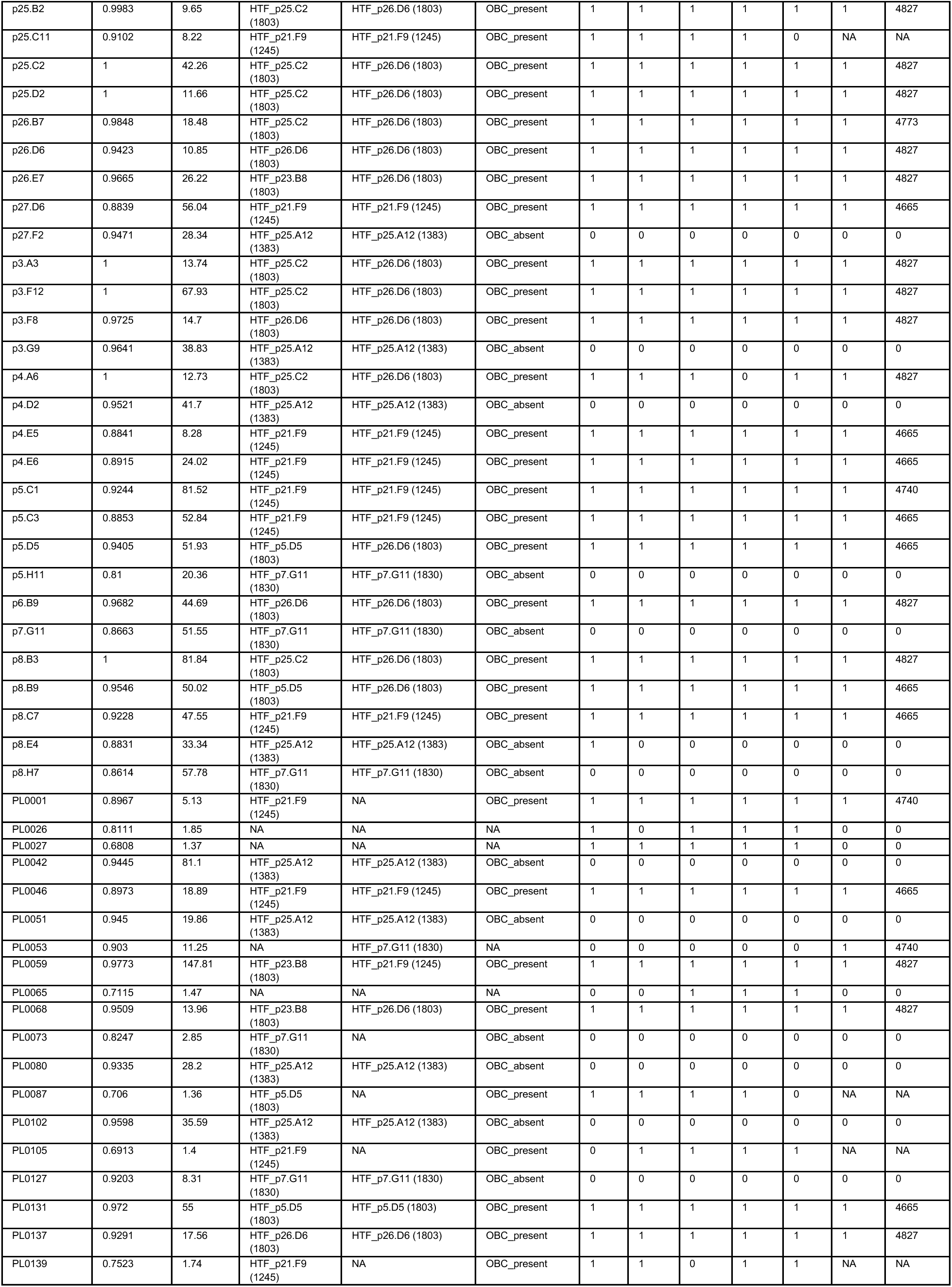
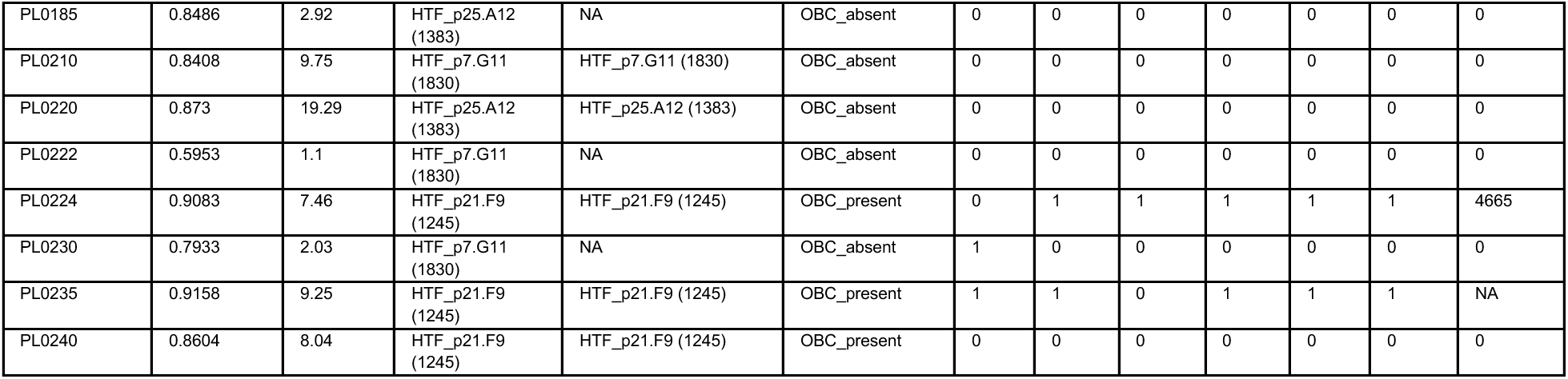
Summary of tailocin coverage, HTF haplotype assignment, and O-antigen presence in historical samples.

## Key Resources Table

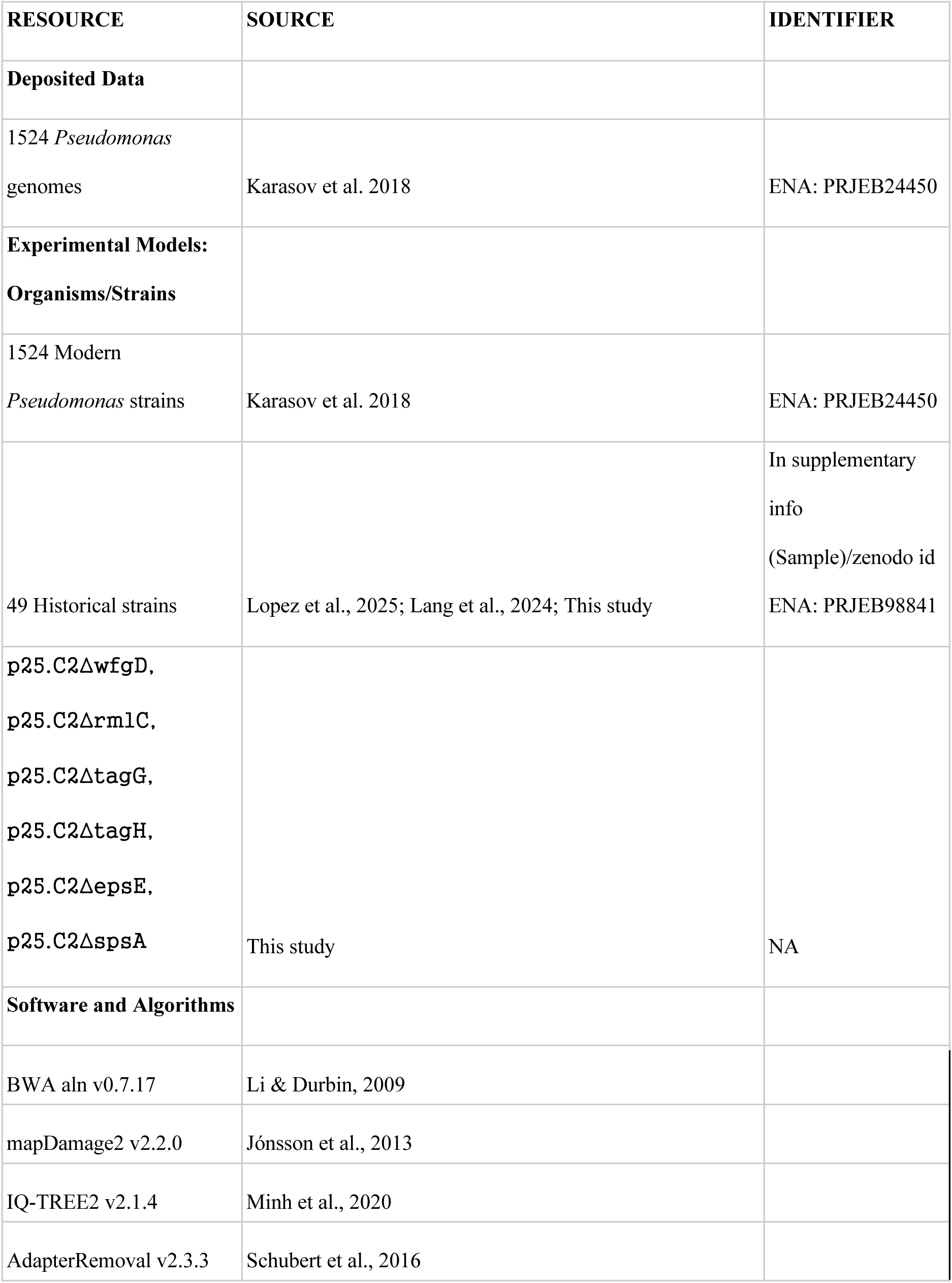
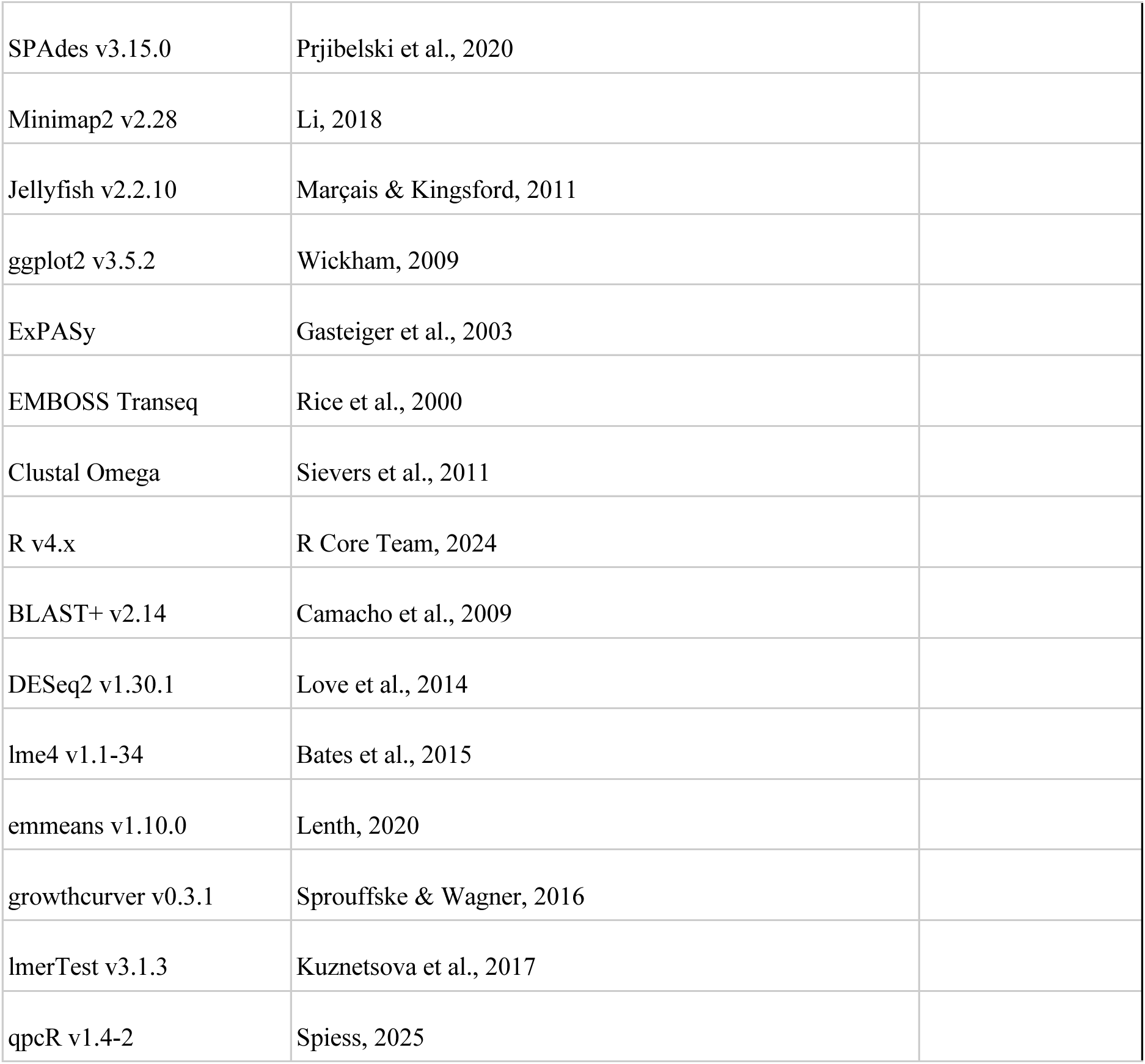

## Modern sample metadata

**Modern sample metadata.**
| SAMPLE | TYPE* | YEAR | COUNTRY | LOCATION | LAT | LON | Pseudomonas_Genome_Coverage_Prop. | Pseudomonas_Genome_Average_Depth_MQ30 |
| --- | --- | --- | --- | --- | --- | --- | --- | --- |
| p12.A11 | Modern | 2018 | Germany | Eyach | 48.446111 | 8.781611 | 0.955211 | 43.2805 |
| p12.A9 | Modern | 2018 | Germany | Eyach | 48.446111 | 8.781611 | 0.385298 | 2.68517 |
| p12.E2 | Modern | 2018 | Germany | Eyach | 48.446111 | 8.781611 | 0.946364 | 124.361 |
| p12.F2 | Modern | 2018 | Germany | Det-2 | 48.556255 | 9.135424 | 0.941151 | 61.1632 |
| p12.G7 | Modern | 2018 | Germany | Det-2 | 48.556255 | 9.135424 | 0.942483 | 26.1727 |
| p12.H7 | Modern | 2018 | Germany | Det-2 | 48.556255 | 9.135424 | 0.703408 | 7.44366 |
| p13.C1 | Modern | 2018 | Germany | Eyach | 48.446111 | 8.781611 | 0.919667 | 91.4711 |
| p13.C7 | Modern | 2018 | Germany | Eyach | 48.446111 | 8.781611 | 0.915668 | 30.2041 |
| p13.D10 | Modern | 2018 | Germany | Eyach | 48.446111 | 8.781611 | 0.917782 | 83.4618 |
| p13.D5 | Modern | 2018 | Germany | Eyach | 48.446111 | 8.781611 | 0.373545 | 3.2645 |
| p13.F1 | Modern | 2018 | Germany | Eyach | 48.446111 | 8.781611 | 0.548638 | 9.89508 |
| p13.F3 | Modern | 2018 | Germany | Eyach | 48.446111 | 8.781611 | 0.930129 | 180.182 |
| p13.F5 | Modern | 2018 | Germany | Eyach | 48.446111 | 8.781611 | 0.486993 | 7.68403 |
| p20.B10 | Modern | 2018 | Germany | Eyach | 48.446111 | 8.781611 | 0.362051 | 6.0333 |
| p20.D4 | Modern | 2018 | Germany | Eyach | 48.446111 | 8.781611 | 0.965996 | 26.275 |
| p20.F10 | Modern | 2018 | Germany | Eyach | 48.446111 | 8.781611 | 0.951949 | 27.3486 |
| p20.G9 | Modern | 2018 | Germany | Eyach | 48.446111 | 8.781611 | 0.951606 | 25.8164 |
| p21.A8 | Modern | 2018 | Germany | Eyach | 48.446111 | 8.781611 | 0.951989 | 50.4423 |
| p21.E3 | Modern | 2018 | Germany | Eyach | 48.446111 | 8.781611 | 0.912764 | 42.5976 |
| p21.F1 | Modern | 2018 | Germany | Eyach | 48.446111 | 8.781611 | 0.921125 | 54.2176 |
| p21.F9 | Modern | 2018 | Germany | Eyach | 48.446111 | 8.781611 | 0.951456 | 43.9553 |
| p22.A8 | Modern | 2018 | Germany | Eyach | 48.446111 | 8.781611 | 0.910773 | 42.5346 |
| p22.B5 | Modern | 2018 | Germany | Eyach | 48.446111 | 8.781611 | 0.908795 | 31.8141 |
| p22.C1 | Modern | 2018 | Germany | Eyach | 48.446111 | 8.781611 | 0.910697 | 44.4703 |
| p22.D1 | Modern | 2018 | Germany | Eyach | 48.446111 | 8.781611 | 0.93509 | 9.86993 |
| p22.D4 | Modern | 2018 | Germany | Eyach | 48.446111 | 8.781611 | 0.901368 | 6.48617 |
| p23.A3 | Modern | 2018 | Germany | Det-2 | 48.556255 | 9.135424 | 0.922147 | 90.3312 |
| p23.A5 | Modern | 2018 | Germany | Det-2 | 48.556255 | 9.135424 | 0.355851 | 5.69028 |
| p23.B4 | Modern | 2018 | Germany | Det-2 | 48.556255 | 9.135424 | 0.353754 | 10.3529 |
| p23.B8 | Modern | 2018 | Germany | Pfrondorf2 | 48.561087 | 9.109294 | 0.915323 | 25.8754 |
| p24.B5 | Modern | 2018 | Germany | Eyach | 48.446111 | 8.781611 | 0.906998 | 24.3191 |
| p24.H2 | Modern | 2018 | Germany | Eyach | 48.446111 | 8.781611 | 0.919194 | 36.961 |
| p25.A12 | Modern | 2018 | Germany | Eyach | 48.446111 | 8.781611 | 0.904389 | 8.27369 |
| p25.B2 | Modern | 2018 | Germany | Eyach | 48.446111 | 8.781611 | 0.99527 | 9.73957 |
| p25.C11 | Modern | 2018 | Germany | Eyach | 48.446111 | 8.781611 | 0.936486 | 7.67613 |
| p25.C2 | Modern | 2018 | Germany | Eyach | 48.446111 | 8.781611 | 0.999957 | 38.9734 |
| p25.D2 | Modern | 2018 | Germany | Eyach | 48.446111 | 8.781611 | 0.997331 | 11.0366 |
| p26.B7 | Modern | 2018 | Germany | Pfrondorf2 | 48.561087 | 9.109294 | 0.918811 | 18.5831 |
| p26.C10 | Modern | 2018 | Germany | Det-2 | 48.556255 | 9.135424 | 0.532614 | 7.27294 |
| p26.D6 | Modern | 2018 | Germany | Eyach | 48.446111 | 8.781611 | 0.907977 | 10.1768 |
| p26.E7 | Modern | 2018 | Germany | Pfrondorf2 | 48.561087 | 9.109294 | 0.915118 | 26.4647 |
| p26.F6 | Modern | 2018 | Germany | Pfrondorf2 | 48.561087 | 9.109294 | 0.263831 | 1.09767 |
| p27.C5 | Modern | 2018 | Germany | Pfrondorf2 | 48.561087 | 9.109294 | 0.33096 | 4.98267 |
| p27.D6 | Modern | 2018 | Germany | Pfrondorf2 | 48.561087 | 9.109294 | 0.915116 | 62.9111 |
| p27.F2 | Modern | 2018 | Germany | Pfrondorf2 | 48.561087 | 9.109294 | 0.908864 | 30.4347 |
| p3.A3 | Modern | 2018 | Germany | Eyach | 48.446111 | 8.781611 | 0.999479 | 12.1995 |
| p3.F12 | Modern | 2018 | Germany | Eyach | 48.446111 | 8.781611 | 0.999766 | 64.6266 |
| p3.F8 | Modern | 2018 | Germany | Eyach | 48.446111 | 8.781611 | 0.910145 | 14.5013 |
| p3.G11 | Modern | 2018 | Germany | Eyach | 48.446111 | 8.781611 | 0.405107 | 35.7165 |
| p3.G9 | Modern | 2018 | Germany | Eyach | 48.446111 | 8.781611 | 0.936757 | 40.7657 |
| p4.A6 | Modern | 2018 | Germany | Eyach | 48.446111 | 8.781611 | 0.999359 | 11.257 |
| p4.A8 | Modern | 2018 | Germany | Eyach | 48.446111 | 8.781611 | 0.317566 | 8.57523 |
| p4.C5 | Modern | 2018 | Germany | Eyach | 48.446111 | 8.781611 | 0.281958 | 2.19208 |
| p4.D11 | Modern | 2018 | Germany | Eyach | 48.446111 | 8.781611 | 0.329928 | 7.72875 |
| p4.D2 | Modern | 2018 | Germany | Eyach | 48.446111 | 8.781611 | 0.903089 | 43.5342 |
| p4.E5 | Modern | 2018 | Germany | Eyach | 48.446111 | 8.781611 | 0.905147 | 9.07888 |
| p4.E6 | Modern | 2018 | Germany | Eyach | 48.446111 | 8.781611 | 0.907851 | 26.4274 |
| p4.H3 | Modern | 2018 | Germany | Det-2 | 48.556255 | 9.135424 | 0.359271 | 11.1964 |
| p5.A5 | Modern | 2018 | Germany | Eyach | 48.446111 | 8.781611 | 0.569164 | 20.9644 |
| p5.C1 | Modern | 2018 | Germany | Eyach | 48.446111 | 8.781611 | 0.940631 | 83.5842 |
| p5.C3 | Modern | 2018 | Germany | Eyach | 48.446111 | 8.781611 | 0.911686 | 61.8474 |
| p5.D5 | Modern | 2018 | Germany | Eyach | 48.446111 | 8.781611 | 0.908126 | 47.1102 |
| p5.F2 | Modern | 2018 | Germany | Eyach | 48.446111 | 8.781611 | 0.355109 | 10.8067 |
| p5.H11 | Modern | 2018 | Germany | Eyach | 48.446111 | 8.781611 | 0.912223 | 24.6014 |
| p6.A10 | Modern | 2018 | Germany | Eyach | 48.446111 | 8.781611 | 0.967914 | 86.1443 |
| p6.B5 | Modern | 2018 | Germany | Eyach | 48.446111 | 8.781611 | 0.377432 | 8.17665 |
| p6.B9 | Modern | 2018 | Germany | Eyach | 48.446111 | 8.781611 | 0.915641 | 40.9034 |
| p6.D10 | Modern | 2018 | Germany | Eyach | 48.446111 | 8.781611 | 0.362251 | 6.29957 |
| p6.E9 | Modern | 2018 | Germany | Eyach | 48.446111 | 8.781611 | 0.348102 | 5.45088 |
| p6.F1 | Modern | 2018 | Germany | Det-2 | 48.556255 | 9.135424 | 0.441901 | 14.1066 |
| p6.G2 | Modern | 2018 | Germany | Det-2 | 48.556255 | 9.135424 | 0.591204 | 22.8581 |
| p6.G3 | Modern | 2018 | Germany | Det-2 | 48.556255 | 9.135424 | 0.374359 | 12.0216 |
| p7.F2 | Modern | 2018 | Germany | Eyach | 48.446111 | 8.781611 | 0.420044 | 6.6193 |
| p7.G11 | Modern | 2018 | Germany | Eyach | 48.446111 | 8.781611 | 0.922476 | 63.4571 |
| p8.B3 | Modern | 2018 | Germany | Eyach | 48.446111 | 8.781611 | 0.999766 | 72.1507 |
| p8.B9 | Modern | 2018 | Germany | Eyach | 48.446111 | 8.781611 | 0.921803 | 46.3749 |
| p8.C7 | Modern | 2018 | Germany | Eyach | 48.446111 | 8.781611 | 0.933317 | 51.4209 |
| p8.D11 | Modern | 2018 | Germany | Eyach | 48.446111 | 8.781611 | 0.432979 | 9.92024 |
| p8.E4 | Modern | 2018 | Germany | Eyach | 48.446111 | 8.781611 | 0.917628 | 39.6076 |
| p8.G10 | Modern | 2018 | Germany | Det-2 | 48.556255 | 9.135424 | 0.460138 | 12.253 |
| p8.G2 | Modern | 2018 | Germany | Det-2 | 48.556255 | 9.135424 | 0.450816 | 8.26752 |
| p8.H7 | Modern | 2018 | Germany | Det-2 | 48.556255 | 9.135424 | 0.899835 | 67.821 |
| p9.C4 | Modern | 2018 | Germany | Eyach | 48.446111 | 8.781611 | 0.946827 | 40.897 |
| p9.H10 | Modern | 2018 | Germany | Eyach | 48.446111 | 8.781611 | 0.469617 | 11.8808 |

